# Korean endemic species no more: on the occurrence of *Pelophylax chosenicus* in China

**DOI:** 10.1101/2022.03.23.485556

**Authors:** Amaël Borzée, Yucheol Shin, Yoonhyuk Bae, Siti N. Othman

**Affiliations:** Laboratory of Animal Behaviour and Conservation, College of Biology and the Environment, Nanjing Forestry University, Nanjing, People’s Republic of China; Department of Biological Sciences, College of Natural Science, Kangwon National University, Chuncheon, Republic of Korea

**Keywords:** anuran surveys, landscape modeling, northeast Asia, range extension, species identification, water frog

## Abstract

Understanding species distribution is the first requirement to work on their behaviour, conservation, and phylogeography. Over the last decades, the number of species described on the Korean peninsula has significantly increased, but areas around the boundaries of the Korean peninsula still have to be surveyed for the presence of these species, especially where the environment is similar and connected. Here, we conducted surveys in the continuous landscapes of the Republic of Korea, the Democratic People’s Republic of Korea and the People’s Republic of China to determine the range of the Gold-spotted pond frog, *Pelophylax chosenicus*. The surveys were conducted between 2016 and 2020 through visual and call encounters. We also used molecular tools to confirm the species identity of the northernmost population, in China, sequencing the mitochondrial DNA *16S* gene fragment. We then determined the importance of landscape types for the species, and especially rice paddies, and used landscape models to define suitable habitats across the region. We found the species to be widespread in low elevation wetlands along the coast of the Yellow Sea, with two isolated populations on the south-east of the Korean Peninsula, and the northernmost population in the vicinity of Dandong in PR China. As this species is listed as threatened in the Republic of Korea, knowing its exact distribution will be important for conservation practices, and this first record for PR China provides a baseline for further surveys.

## Introduction

Resolving the distribution of species is a key requirement to understand their diversity and behavioural ecology, but also the threats they face (Manne and Pimm 2001; Richardson and Whittaker 2010). Historically, conducting field surveys was one of the first research efforts to study biodiversity (Wallace 1876), and determining the occurrence of species is needed to understand and protect biodiversity (Wang et al. 2021). Occurrence surveys are generally based on field surveys, automated monitoring, surveys via citizen science, and literature reviews (Fletcher Jr et al. 2019). The raw data can then be complemented through the use of ecological models to determine the suitable habitat for the species (Araújo et al. 2019). This is especially true in the case of species ranging across an inaccessible area, such as species distributed across the Korean Peninsula and northeast People’s Republic of China (hereafter China), for which the distribution in the Democratic People’s Republic of Korea (hereafter DPR Korea) can only be rarely surveyed, but can be modelled adequately (Borzée et al. 2021).

The genus *Pelophylax* is widespread in Eurasia, likely originating from the western Palearctic (Pyron 2014), with the East Asian clades having diverged about 6.2 mya (Liu et al. 2010). The genus is almost continuously distributed across the Palearctic as some species are also found in the dry landscapes of central Asia. In northeast Asia, the two main clades, the *Pelophylax nigromaculatus* and *Pelophylax plancyi* complexes (following Liu et al. 2010) have diverged about 4.8 mya, but they have a history of hybridisation, with cytonuclear discordance in numerous populations, likely in relation with glacial cycling (Komaki et al. 2015). The genus is widespread across landscapes as species have broad ecological requirements, some species are able to breed in most type of wetlands (Garcia et al. 2017; e.g. *P. nigromaculatus* in northeast Asia; Borzée et al. 2021), and they are good invaders (Wang et al. 2016; Wang et al. 2017; Bae et al. 2022). Niche segregation between the two species complexes is also likely to occur based on prey preference and microhabitat use (Eom et al. 2007; Borzée et al. 2019b; Nakanishi et al. 2020), with the *P. plancyi* complex expected to have more stringent ecological requirements. *Pelophylax chosenicus* is closely related to *P. plancyi*, and the time of divergence between the two clades is so far unresolved, although it is likely to be relatively recent (Liu et al. 2010). However, the two species are distinct genetically (mtDNA; Ryu and Hwang 2011) and morphologically distinct, with male *P. plancyi* having clearly visible air sacs (Fei et al. 2012) and male *P. chosenicus* generally not having air sacs, or having internal air sacs (Kim et al. 2019), despite some exceptions possibly related to hybridisation with *P. nigromaculatus* as males of the latter species have large air sacs and the two species hybridise (Komaki et al. 2015). *Pelophylax chosenicus* is distributed along the western coast of the Korean Peninsula, until an unknown northern boundary. Following multiple short notes on northward range extension (Borzée et al. 2017a; Borzée et al. 2018b; Son and Borzée 2020), and modeling showing that the species likely extends further north than surveyed (Borzée et al. 2021), more information is needed on the distribution of the species, including the coastal area in China. Therefore, this study was designed to understand the distribution of *P. chosenicus* through field surveys and modelling, relying on molecular tools to identify the northernmost population at the species level.

## Material and methods

### Species introduction

Pelophylax chosenicus was described in 1926 (Okada 1926) and the species can be estimated to have been abundant at the beginning on the 20th century through the publications available (Okada 1928; Okada 1931; Shannon 1956). Hybridisation with sympatric *Pelophylax* species has been known early on, as it was one of the first focus of research on the species (Nishioka 1972; Nishioka and Okumoto 1983). The first report on the behavioural ecology of the species was first published in 1998 (Yoon et al. 1998), with an increasing understanding of the behavioural ecology of the species (Do et al. 2021). The need for conservation assessments and programs was however rapidly understood (Shim 2003; Ra et al. 2008a; Park et al. 2009).

### Field surveys

Surveys were conducted between 2015 and 2021, mixing both normalised transects and opportunistic observations. Normalised surveys were conducted separately in the Republic of Korea (R Korea hereafter), in DPR Korea and in China. The surveys in R Korea were conducted in conjunction with surveys conducted for Hylid populations (Borzée et al. 2017b; Borzée et al. 2018a) as the target species, *Pelophylax chosenicus*, generally shares its breeding habitat with *Dryophytes suweonensis* and *D. flaviventris* (Borzée et al. 2020b). The surveys were conducted between late April and early July, matching with the breeding season of the species (Seo et al. 2014). Survey sites were broadly selected to match with the habitat types where the species occurs, including both natural and agricultural wetlands. Each site was surveyed through an aural transect, and the presence and absence of the species was binary encoded for each site (0 = absence; 1 = presence). The time of day and weather patterns were selected to match with the activity windows of the species (Yang et al. 2000). The exact details of the protocol can be found in Borzée et al. (2017c). In addition, opportunistic survey transects were conducted across the nation, always matching with the breeding season of the species and in potentially adequate habitats (survey sites in Figure 1 and GPS coordinates in Supplementary file 1).

**Figure 1.**
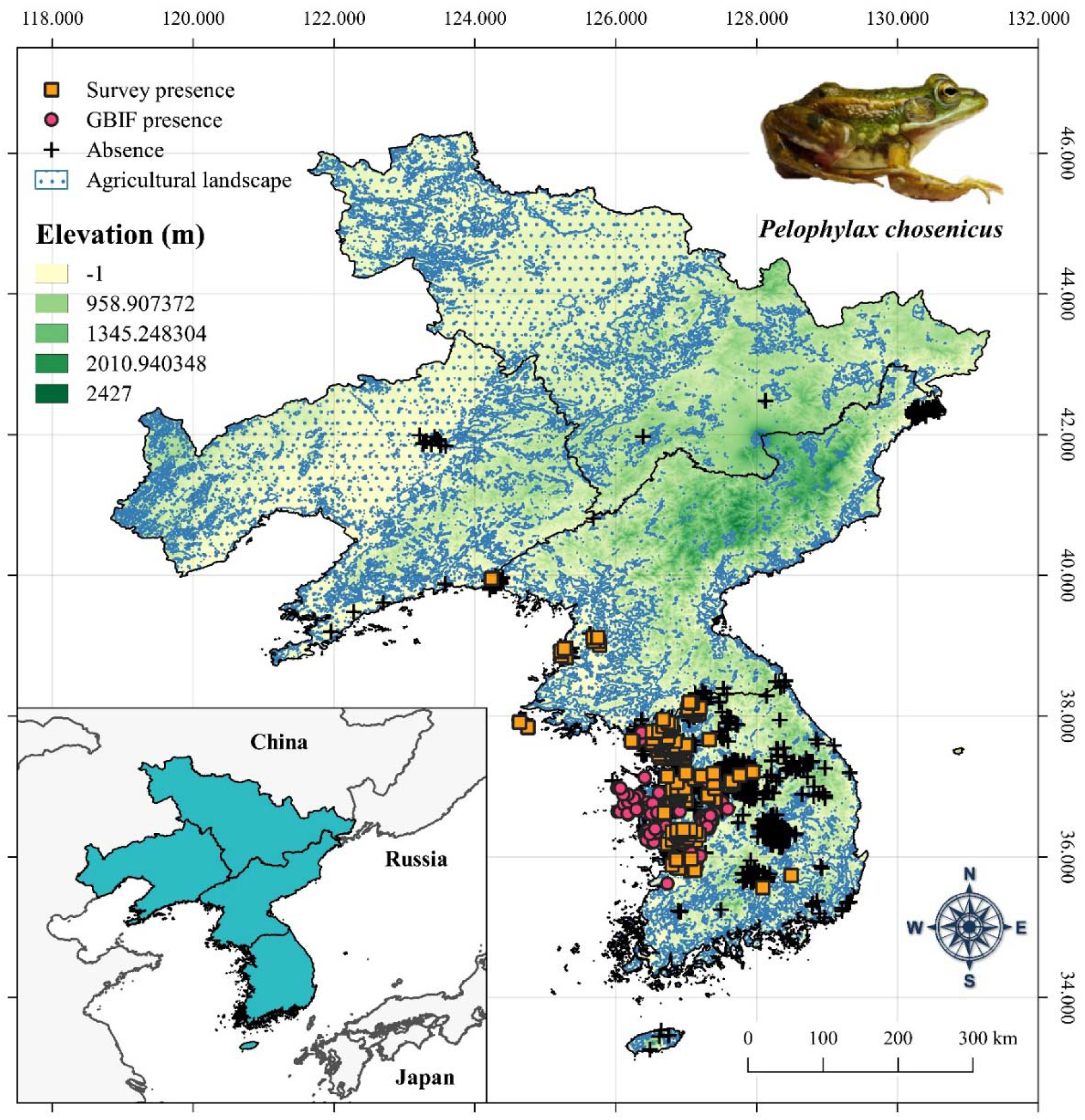
Spatial distribution of presence (*n* = 403; 273 field survey point and 130 from GBIF and one museum datapoint) and putative absence points from field surveys (*n* = 1280) for *Pelophylax chosenicus* across the study area. The presence points are divided into unique survey-based data and occurrence data derived from the Global Biodiversity Information Facility (GBIF.org; DOI: 10.15468/dl.6kvymf). The agricultural landscapes are represented as areas with agricultural landscape cover greater than 50%. The location of study area (blue polygon) in northeast Asia is shown in the inset map. Map computed with QGIS3 (QGIS Development Team (2022). QGIS Geographic Information System. Open Source Geospatial Foundation Project. http://qgis.osgeo.org).

**Figure 2.**
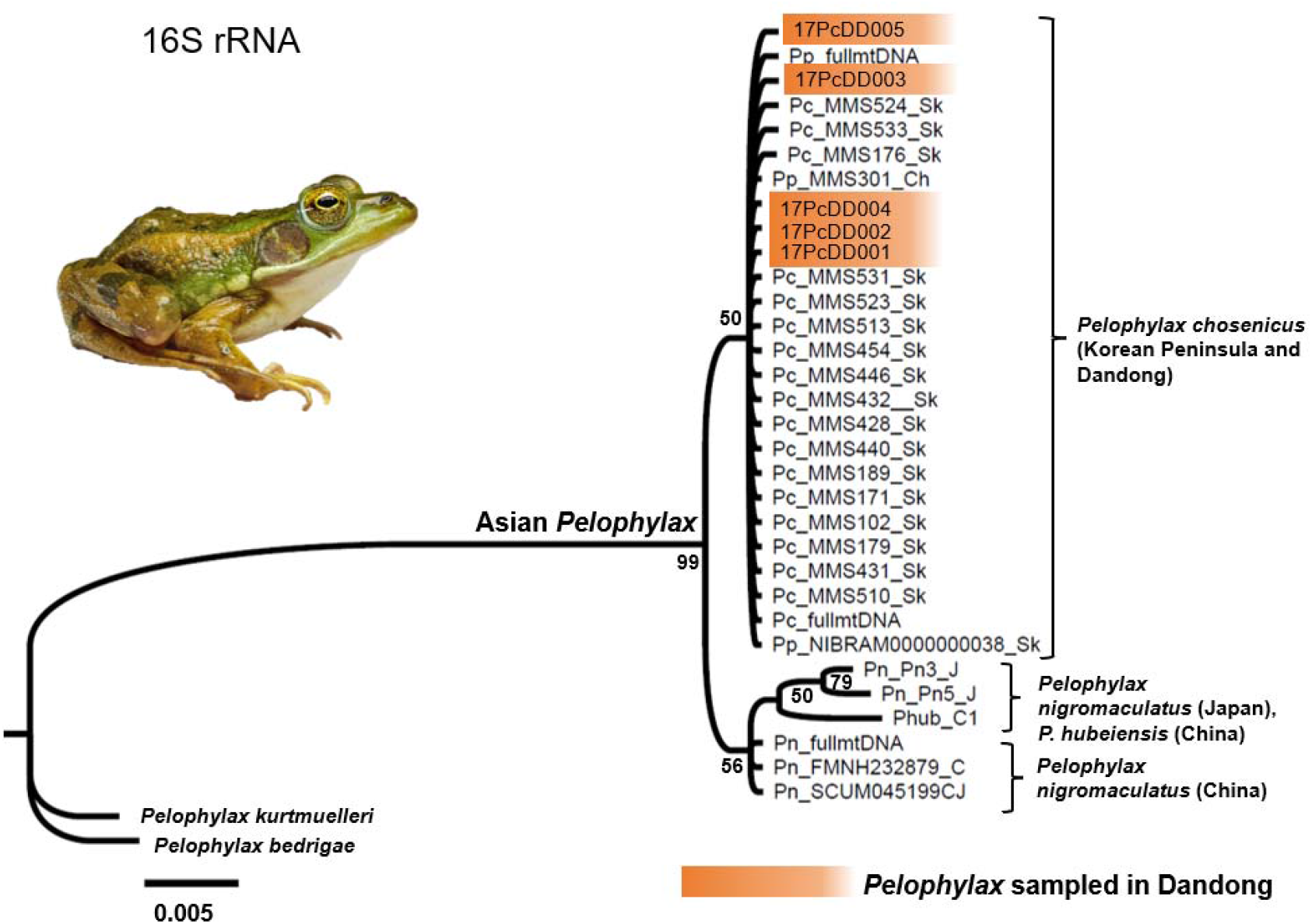
Phylogenetic tree for *Pelophylax* individuals sampled in Dandong in relation with sympatric species. Phylogeny reconstructed using the 16S rRNA marker (554 bp; *N* = 34). The percentage of posterior probability is indicated for each node.

Surveys in DPR Korea were conducted opportunistically in 2017 and 2018, followed by surveys in 2019 relying on the same protocol as the one used in R Korea. The methodology and datapoints are presented in a country wide review (Borzée et al. 2021).

Surveys in China were conducted in June 2018, following the same protocol as the one used in R Korea, but within an area geographically restricted to the regions surrounding the city of Dandong. Additional opportunistic survey transects were conducted at four sites along the coast and further along the border between China and DPR Korea (Figure 1 and GPS coordinates in Supplementary file 1).

For all transects, the GPS coordinates recorded were the ones where the transect started, and as no transect was longer than 500 m it did not impact subsequent analyses. In total, we conducted 1674 surveys. While the probability of detecting the species should be close to the highest value possible with the protocol used (Petitot et al. 2014; Başkale and Çapar 2016), and specifically spending at least 5 min at each site, we consider each transect where the species was not detected as a putative absence point later in the manuscript. The dataset was also complemented by a museum datapoint (Shin et al. 2020), resulting in an additional presence point (37.6678°N, 127.3264°E).

### Molecular identification

To further verify the occurrence of *Pelophyalx chosenicus* in China, we identified the sampled individuals during our survey in Dandong, China through barcoding and phylogenetic analyses. All 5 individuals were captured on the spot during the survey and orally swabbed (Dryswab™ ENT, Medical Wire, Wiltshire, UK). The individuals were released back to the sampling point within 5 min, and the buccal swabs were kept in dry conditions with desiccation beads until DNA extraction. We extracted genomic DNA using the DNeasy Blood and Tissue Extraction Kit (Qiagen, Hilden, Germany), following the manufacturer’s protocol. We determined the concentration of each DNA sample using a NanoDrop™ 2000/2000c.

We then amplified the mitochondrial 16S rRNA gene fragment using a Polymerase Chain Reaction (PCR). We selected this marker as it is commonly used for barcoding of northeast Asian ranids, but we excluded the gene fragments Cytochrome b (Cyt*b*) and 12S rRNA as preliminary analyses showed that they are not phylogenetically informative. The primer pair targeted a 382 bp-long gene fragment (Sumida et al. 2002) with the following primers FS01 (5’-AAC GCT AAG ATG AAC CCT AAA AAG TTC T-3’) and R16 (5’-ATA GTG GGG TAT CTA ATC CCA GTT TGT TTT-3’). We prepared the PCR reaction with a total volume of 20 μL per tube for each template of DNA, and each samples contained 35 to 50 ng/μL of DNA. The final concentrations of our PCR reactions were 0.125 μM for each forward and reverse primer, 1x Ex taq Buffer (Takara; Shiga, Japan), 1.875 mM of magnesium chloride (MgCl_2_), 0.2 mM of dNTPs Mix (Takara; Shiga, Japan), 0.1 unit/μL of Ex taq (HR001A, Takara; Shiga, Japan), and double distilled water added to make up the final volume. We performed the PCR amplifications using a SimpliAmpTM Thermal Cycler (Applied Biosystems, USA), under the following thermal conditions: initial denaturation for 5 min at 95°C; 35 cycles of denaturation at 94°C for 1 min, annealing at 55°C for 1 min, extension at 72°C for 1 min, followed by a final extension at 72°C for 10 min. To screen the PCR products, we ran an electrophoresis on a 1.5% agarose gel loaded with 3 μL of PCR amplicons and 1 uL of TopGreen Nucleic Acid 6x Loading Dye (GenomicBase, Republic of Korea). All PCR amplicons were sent for purification and Sanger sequencing for both forward and reverse directions to Cosmogenetech (Cosmogenetech Co., Ltd., Seoul, Republic of Korea). We deposited all generated sequences in GenBank (accession numbers in Supplementary file 2).

To confirm the identity of the individuals sampled in Dandong, we then reconstructed the phylogeny of the species using the 16 rRNA gene fragment, including the sympatric species of the region. To do so, we first retrieved 29 homologous sequences from BLASTn and GeneBank (accession numbers in Supplementary file 2; originating from Jiang and Zhou 2005; Frost et al. 2006; Ryu and Hwang 2011; Jeong et al. 2013; Hofman et al. 2016; Yuan et al. 2016; Jiang et al. 2017; Tokumoto et al. 2019). We used *P. bedrigae* and *P. kurtmuelleri* as outgroups. We analysed and trimmed the DNA sequences using Geneious Prime version 2022.1 (http://www.*geneious*.com, Kearse et al. 2012), and aligned sequences using Clustal Omega (Sievers et al. 2011) with its default setting under the same Geneious Prime programme (Table S1). The trimmed sequences and final alignments used for the phylogenetic analyses were 554 bp for 16S rRNA gene fragment (*n* = 34).

Finally, we reconstructed the phylogeny using the Bayesian Inference (BI) method in Mr Bayes version 3.2.6 (Huelsenbeck and Ronquist 2001). We determined the best evolutionary model using Partition Finder version 2.1.1 (Lanfear et al. 2017) and obtained K80 + I as best model for the non-coding 16S rRNA gene fragment. In Mr. Bayes, we entered the input file containing the 16S rRNA gene alignments and ran the analysis for ten million generations. We sampled the Markov chain every 1000 generations and discarded the first 10% of the generations. We ensured the analyses converged, indicated by an average standard deviation of split frequencies <0.05. To verify whether the tree reached its stationary state, we evaluated the meaningfulness of each parameter by the value of effective sample size (ESS) of each posterior distribution (above 200) using TRACER version 1.7.1 (Rambaut et al. 2018).

### Landscape features

To analyse the landscape features used by *P. chosenicus*, we combined the data obtained from our field surveys with records from the Global Biodiversity Information Facility (GBIF; DOI: 10.15468/dl.6kvymf.), collected through the *megaSDM* package in R (Shipley et al. 2022), by running the ‘OccurrenceCollection’ function. The combined datasets resulted in a total of 403 occurrence points (Figure 1). In addition to using presence points, we used the 1280 putative absence points recorded from our surveys to compare landscape features between presence and absence points (Figure 1). Next, we spatially thinned the occurrence data using a thinning distance of 500 m, resulting in 273 presence points and 922 putative absence point. Finally, we extracted landscape types at these spatially thinned presence and absence sites. To do so, we downloaded a 500-m resolution global land cover raster from the Global Land Cover by National Mapping Organizations (GLCNMO version 3; Kobayashi et al. 2017; available from: https://globalmaps.github.io/glcnmo.html). The raster cells for this dataset were classified into 14 different landscape types, including several landcover types that are suitable habitats for *P. chosenicus* (e.g., natural and agricultural wetlands; data in Supplementary file 1). Finally, we conducted a Chi-square goodness of fit test to determine the relation between the type of landscape for each of the presence (*n* = 273) and putative absence datasets (*n* = 922). Further, due to the expected reliance of the species on rice paddies, we binary encoded the presence of rice paddies for each site (data in Supplementary file 1), and tested for a difference between species occurrence and rice paddies through a third chi-square test.

### Ecological niche modeling

To generate ecological niche models (ENMs) for *P. chosenicus*, we initially considered a set of 27 environmental variables. These include 19 bioclimatic variables and elevation data downloaded from the WorldClim 2.0 database (Fick and Hijmans 2017; https://www.worldclim.org/) and six consensus land cover variables (cultivated area, forest, herbaceous vegetation, open water bodies, shrub, urban areas) and slope data downloaded from EarthEnv (Tuanmu and Jetz 2014; Amatulli et al. 2018; http://www.earthenv.org/). The selection of the six land cover layers is based on the habitat feature analyses of presence points. All environmental layers were downloaded as 1-km spatial resolution rasters.

To define the model calibration area (Soberón and Peterson 2005), we generated a 50-km radius circular buffer (∼ 0.45 dd) around each occurrence point using the ‘BackgroundBuffers’ function of the *megaSDM* package (Shipley et al., 2021). We then masked the environmental layers to the boundaries of the calibration area using R packages *raster* (Hijmans 2021) and *rgdal* (Bivand et al. 2021), and used the masked rasters for downstream niche modeling.

We further spatially thinned the initial occurrence dataset (*n* = 405) using the thinning distance of 1km. This resulted in a total of 273 occurrence points used for model calibration. To generate the background dataset, we implemented a target group background sampling to account for the differences in spatial sampling efforts between China, DPR Korea and R Korea (Barber et al. 2022). To do so, we first downloaded all amphibian occurrence points recorded from within the boundaries of the calibration area from the Global Biodiversity Information Facility (GBIF) via the *megaSDM* package (no DOI assigned to the dataset when extracted through the package). Then, we converted these points to a density raster representing the spatial sampling efforts of amphibian occurrence points within the modeling extent. We used this raster for spatial bias correction in the model calibration process.

Using the R package *SDMtune* (Vignali et al. 2020), we implemented a data-driven variable selection approach to select environmental variables with high importance and low collinearity. First, using 80% of the occurrence points (*n* = 218), 8,000 target group background points, and all 27 environmental variables masked to the calibration area, we trained a Maximum Entropy (MaxEnt) model in default settings and 10 permutations. Next, we used the ‘varImp’ function to check the importance of each variable. We then used the ‘reduceVar’ function to generate a reduced-variable model, applying a percent contribution threshold of 1 and removing variables below this threshold. Next, we used the ‘varSel’ function on the reduced-variable model to further remove highly collinear variables. The threshold for this step was the Spearman’s correlation test coefficient of 0.7. In both steps of variable reduction, we used True Skill Statistic (TSS; Allouche et al. 2006) calculated from the testing dataset to compare the performance of models before and after variable removal. After this step, the following eight environmental variables were retained for model calibration: isothermality (bio3), mean temperature of wettest quarter (bio8), precipitation of driest month (bio14), cultivated landscape cover, herbaceous vegetation cover, open water bodies, urban areas cover, and slope.

Next, we optimized the MaxEnt model parameters using the *ENMeval* package (version 2.0.0; Kass et al. 2021). Using the eight selected environmental variables, all occurrence points, and 10,000 target group background points as inputs and applying a 10-fold cross-validation, we tested the combinations of six MaxEnt feature classes (L, LQ, H, LQH, LQHP, LQHPT) and regularization multipliers (RM) ranging from 0.5 to 8 at the increment of 1.

Among the models output from the *ENMeval* run, we selected the optimal model generated with an H feature combined with a regularization multiplier of 0.5, as this model had both the highest validation AUC and the lowest 10% omission rate (Kass et al. 2021).

Based on the selected optimal model parameters, we fitted and projected the final ENM in 10-fold cross-validation using MaxEnt version 3.4.1 via the *dismo* R package (Hijmans et al. 2020). We also used cloglog output and used jackknife tests to assess variable importance. The ENMs were first trained on the calibration area, and then projected across the entire study area. We evaluated the final model based on the following criteria: 1) area under the Receiver Operating Characteristic curve (AUC), 2) True Skill Statistic (TSS; Allouche et al. 2006), using Maximum Test Sensitivity plus Specificity threshold (= 0.4154) for calculation, 3) AUC_DIFF_ (difference between training and validation AUC; Warren and Seifert 2011), 4) comparisons with known distribution of *P. chosenicus* (NIBR 2019), and 5) comparisons with null models (Bohl et al. 2019; Kass et al. 2021). For comparisons with null models, we used the ‘ENMnulls’ function of the *ENMeval* package (Kass et al. 2021) to simulate 100 iterations of null models with the same parameterization as the empirical model (H feature with RM value of 0.5). The AUC, TSS, and comparisons with null models were used to assess model’s predictive abilities, AUC_DIFF_ was used to assess the degree of model overfitting, and comparisons with known distributions were used to visually assess the consistency between the predicted niches and actual distribution.

Next, to identify potential areas of presence for *P. chosenicus* across our study range, we applied one “liberal” threshold (= 0.2578; 10-percentile training presence; P10) and one “strict” threshold (= 0.4154; Maximum Test Sensitivity plus Specificity; MTSS) to convert the continuous suitability outputs of ENMs into binary presence/absence maps. To generate binary maps, we used the ‘ecospat.binary.model’ function of the *ecospat* R package (Di Cola et al. 2017). We used R version 4.1.0 (R Core Team 2021) for all the analyses conducted herein.

## Results

### Field surveys and molecular identification

The field surveys resulted in 1195 independent datapoints (thinned from the 1674 surveys conducted), including 394 presence points (273 after thinning; complemented with 130 datapoints from GBIF (after thinning) and one datapoint from the museum), and 1280 putative absence points (922 after thinning; Figure 1). In R Korea, we found the species to be ranging from Buan in the south, along with two inland populations, continued northwards through a continuous distribution on the lowlands of the nation, interrupted only by the Chilgap hills and the Seoul-Incheon urban area. All presence datapoints collected in DPR Korea were in the lowlands, and the species is expected to be similarly continuously distributed along the west coast of the Korean Peninsula until the border with China. In China, the only presence points recorded were in the vicinity of Dandong, close to the border with DPR Korea.

The analysis for the five individuals sampled in Dandong using the 16 rRNA gene fragment showed a high genetic homogeneity with *P. chosenicus*, with a percentage of sequences identity ≥ 98% (Blast comparison with GenBank accession numbers JF730436; JF730436; JF730436; JF730436; JF730436). In addition, all five individuals were placed within the monophyletic *P. chosenicus* clade in the BI tree (*n* = 36; PP: 50%).

### Landscape features

Based on 273 presence points, the highest number of occurrences was recorded in agricultural landscapes (*n* = 146; cropland and rice paddies combined), followed by urban areas (*n* = 53), deciduous broadleaf forests (*n* = 15), open canopy forests (*n* = 12), vegetation mosaics (*n* = 11), mixed forests (*n* = 9), water bodies (*n* = 9), evergreen needleleaf forests (*n* = 5), shrubs (*n* = 5), deciduous needleleaf forests (*n* = 4), herbaceous vegetation (*n* = 3), and sparse vegetation (*n* = 1). The distribution of occurrence points was significantly different across land cover types (chi-square test; χ^2^ = 406.38; df = 12; *p* < 0.001; Figure 3).

**Figure 3.**
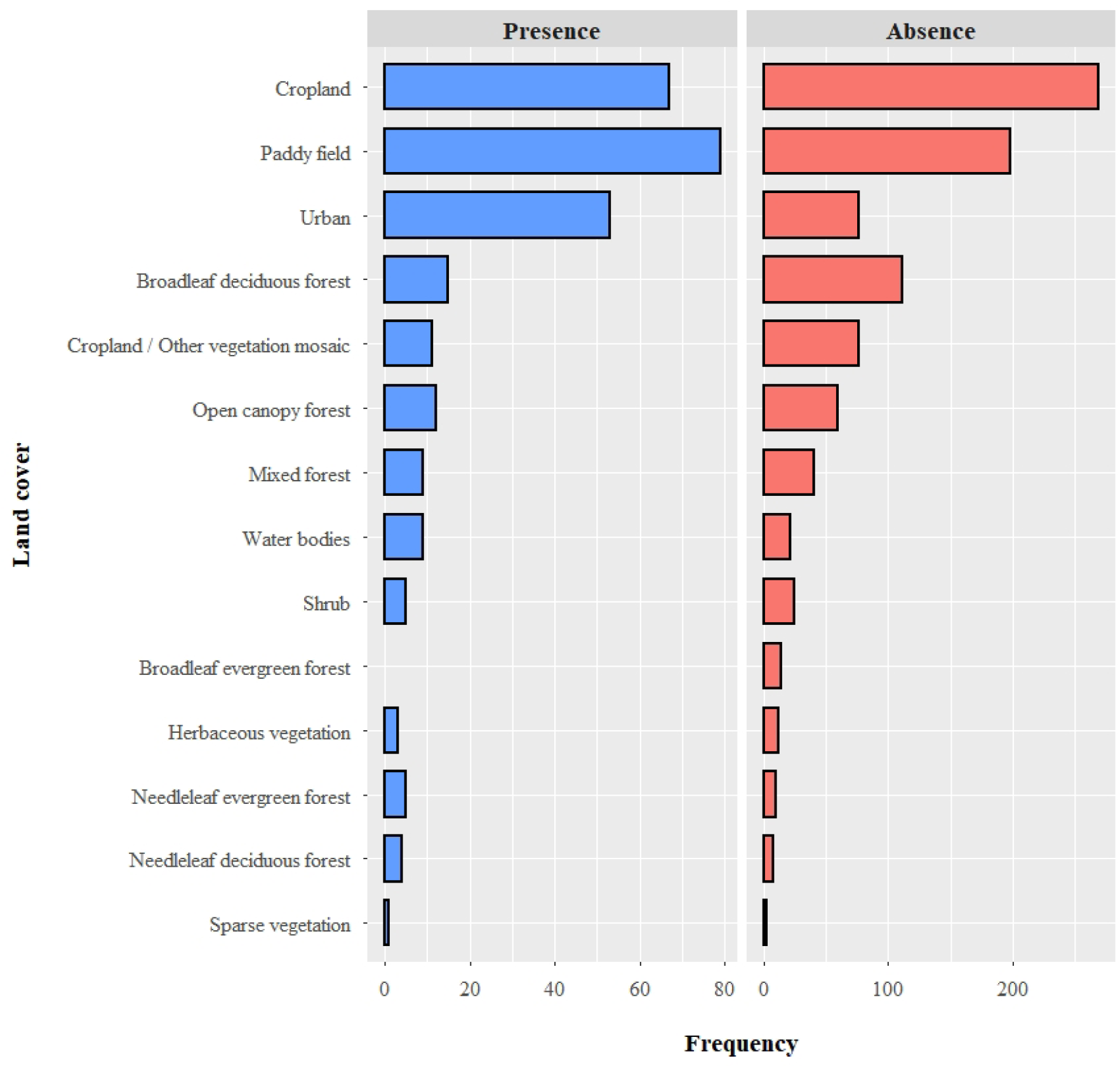
Landscape characteristics at presence (*n* = 273) and putative absence (*n* = 922) sites for *Pelophylax chosenicus*. The distribution of presence and putative absence sites were significantly different across land cover types (chi-square tests; *p* < 0.001). For both presence and absence datasets, agricultural landscapes (croplands and rice paddies combined) account for the highest recorded frequency among landscape types, but the frequency of rice paddies is significantly different for presence and putative absence (chi-square test; *p* < 0.001).

For the 922 putative absence points (after thinning), the highest number of absence points was recorded in agricultural landscapes (*n* = 467), followed by deciduous broadleaf forest (*n* = 111), vegetation mosaics (*n* = 76), urban areas (*n* = 76), open canopy forests (*n* = 60), mixed forests (*n* = 40), shrubs (*n* = 25), water bodies (*n* = 21), evergreen broadleaf forests (*n* = 14), herbaceous vegetation (*n* = 12), evergreen needleleaf forests (*n* = 10), deciduous needleleaf forests (*n* = 8), and sparse vegetation (*n* = 2). The distribution of putative absence points was significantly different across land cover types (χ^2^ = 1237.4; df = 13; *p* < 0.001; Figure 3).

Finally, we found that 70 out of the 273 presence points (25.67%) were located in rice paddies, while 198 out of the 922 putative absence points (21.47%) were located in rice paddies, a significant difference between the presence and putative absence points (χ2 = 343.83; df = 1; *p* < 0.001).

### Ecological niche modeling

The two statistical evaluation metrics indicated high predictive performance of the final ENM (AUC = 0.820 ± 0.027 SD; TSS = 0.553 ± 0.008 SD), while the AUC_DIFF_ value (= 0.023) indicated low model overfitting. In addition, comparisons of validation AUC scores with 100 null ENMs indicated a significantly higher predictive performance of the empirical models (Mann-Whitney *U*-test: *W* = 4800; *p* < 0.001; Figure 4). Furthermore, the plotted continuous habitat suitability model and binary presence/absence maps generated from both P10 (“liberal”) and MTSS (“strict”) thresholds were consistent with the known distribution of *P. chosenicus* across the Korean Peninsula (NIBR 2019). Therefore, we considered the final ENM to be suitable as a representation of habitat suitability for *P. chosenicus* across the extent of our study.

**Figure 4.**
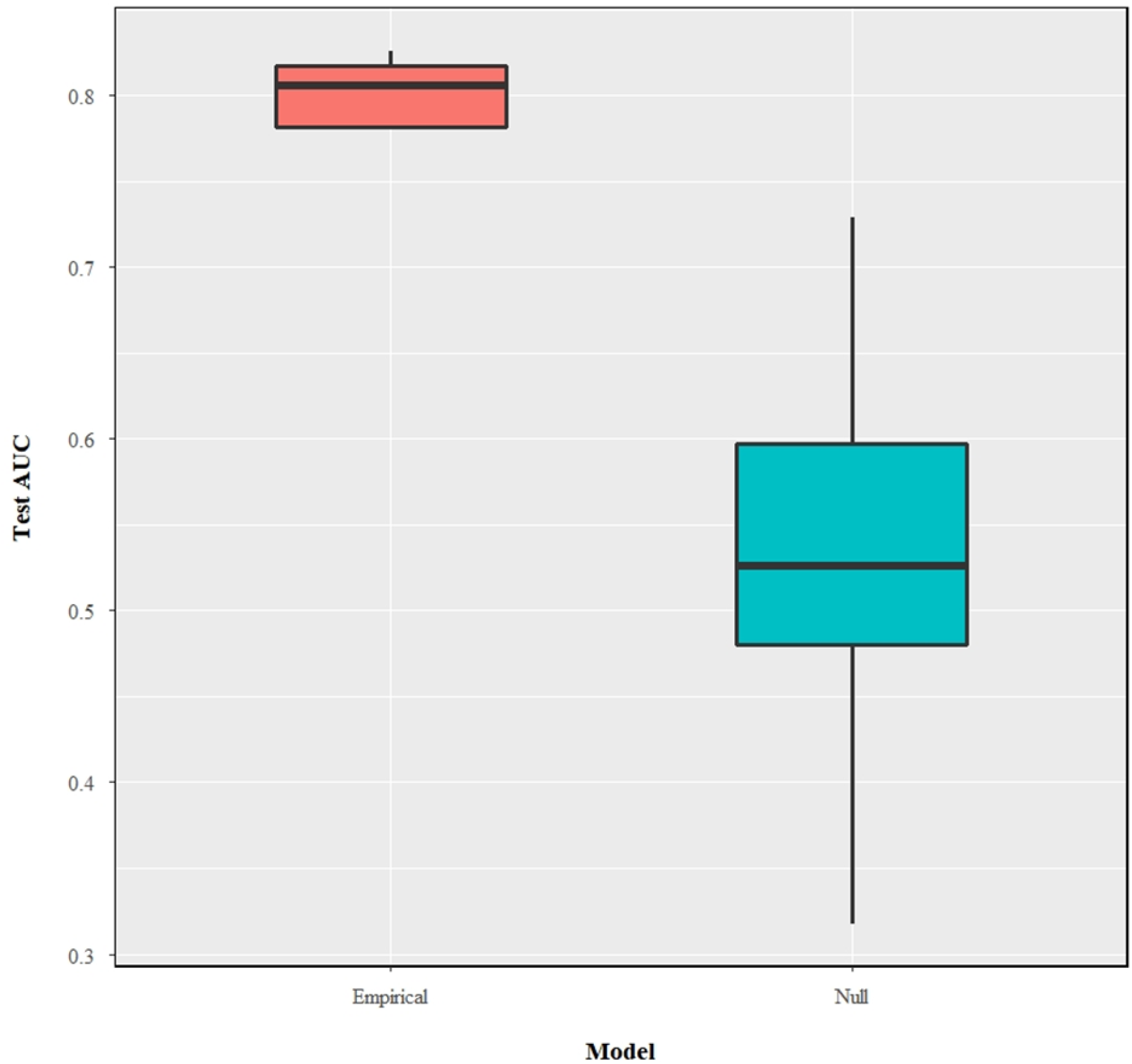
The distribution of test AUC values between the empirical MaxEnt models of *Pelophylax chosenicus* (*n* = 48) and null models (*n* = 100). The empirical models have significantly higher test AUC values compared to the null models (Mann-Whitney *U*-test: *W* = 4800; *p* < 0.001).

Based on percent contribution, isothermality (bio3) had the highest contribution to the model, followed by slope, precipitation of driest month (bio14), mean temperature of wettest quarter (bio8), cultivated landscape cover, urban landscape cover, herbaceous vegetation cover, and the cover of open water bodies (Table 1). Based on permutation importance, isothermality (bio3) had the highest contribution, followed by precipitation of driest month (bio14), mean temperature of wettest quarter (bio8), cultivated landscape cover, slope, herbaceous vegetation cover, the cover of open water bodies, and urban landscape cover (Table 1). According to the response curves, the suitable habitats for *P. chosenicus* across our study area are characterized by isothermality between 22 and 28, mean temperature of wettest quarter above 22°C, precipitation of driest months above 25mm, slope below approximately 15°, cultivated landscape cover greater than 40%, herbaceous vegetation cover approximately below 52%, cover of open water bodies greater than 20%, and cover of urban areas approximately below 85% (Figure 5).

**Table 1.** Percent contribution and permutation importance of environmental variables used to predict current habitat suitability of *Pelophylax chosenicus*. Based on 273 occurrence points compiled from survey results and Global Biodiversity Information Facility (GBIF; data in Supplementary file 1).

| Variable | Percent contribution | Permutation importance |
| --- | --- | --- |
| Isothermality (bio3) | 33.3 | 46.2 |
| Slope | 31.9 | 2 |
| Precipitation of driest month (bio14) | 20.5 | 23.1 |
| Mean temperature of wettest quarter (bio8) | 7.1 | 22.7 |
| Cultivated landscape | 3.8 | 5.1 |
| Urban landscape | 2.3 | 0.2 |
| Herbaceous vegetation | 0.7 | 0.3 |
| Open water bodies | 0.5 | 0.3 |

**Figure 5.**
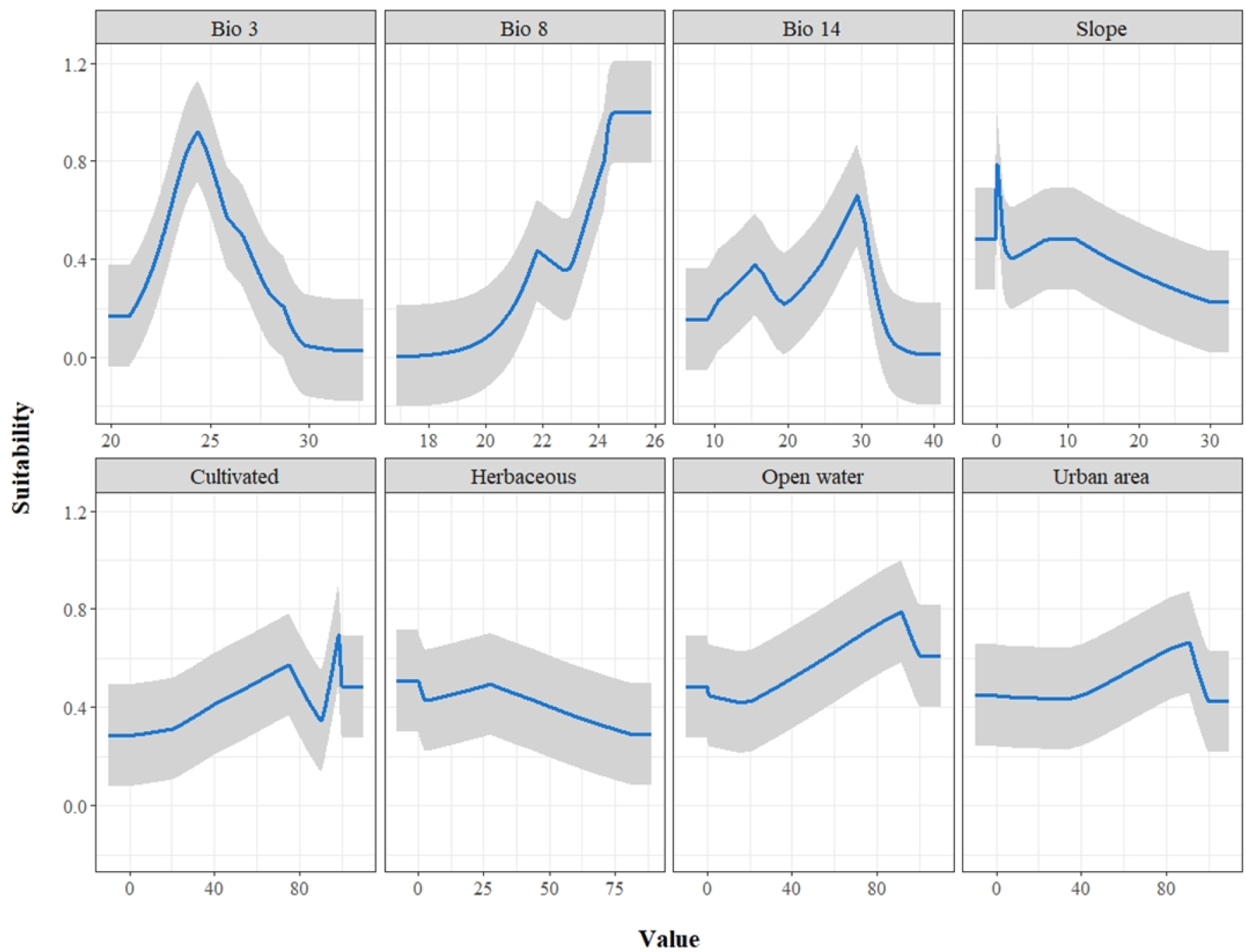
Response curves of *Pelophylax chosenicus* to environmental variables used in ecological niche modeling. See Table 1 for contribution value of each variable. Based on 273 occurrence points compiled from surveys results and Global Biodiversity Information Facility (GBIF; data in Supplementary file 1).

Both continuous and binary MaxEnt outputs show areas of high habitat suitability across the western coast of the Korean Peninsula. The areas of suitable habitat predicted by the ENM were highly consistent with the known distribution across R Korea (NIBR 2019; Figure 6; Figure 7), and while the distribution of the species is incompletely known for DPR Korea, the predicted distribution is highly consistent with a previous model-based range predictions of *P. chosenicus* for the country (Borzée et al. 2021). Our ENM predictions show that the narrow areas of highly suitable habitat along the western coast of the Korean Peninsula continues into the southern coastal regions of Dandong in the Liaoning Province (Figure 6). Across the central Liaoning Province, there is another broad area of predicted suitable habitat continuing into northern Jilin Province. Both P10 (“liberal”) and the MTSS (“strict”) thresholds consistently predicted such patterns of suitable habitats, while the former tended to predict broader areas to be suitable for *P. chosenicus* (Figure 7).

**Figure 6.**
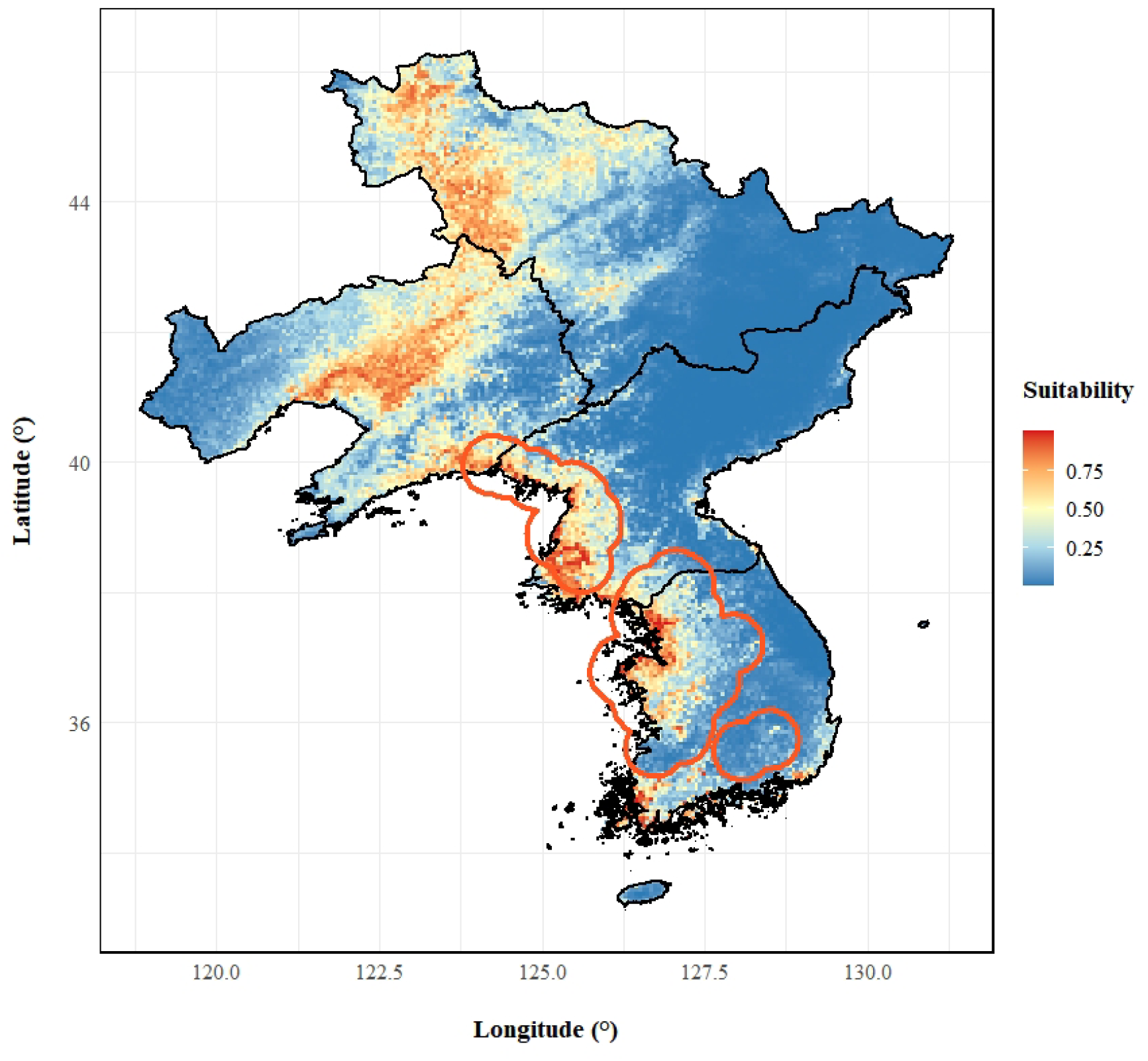
The MaxEnt habitat suitability model for *Pelophylax chosenicus* based on eight environmental variables and 273 occurrence points (AUC = 0.820 ± 0.027 SD; TSS = 0.553 ± 0.008 SD; AUC_DIFF_ = 0.023). The model was first trained on the calibration area (circular polygon) and then projected across the entire study extent encompassing the Korean Peninsula, Jilin province, and Liaoning province.

**Figure 7.**
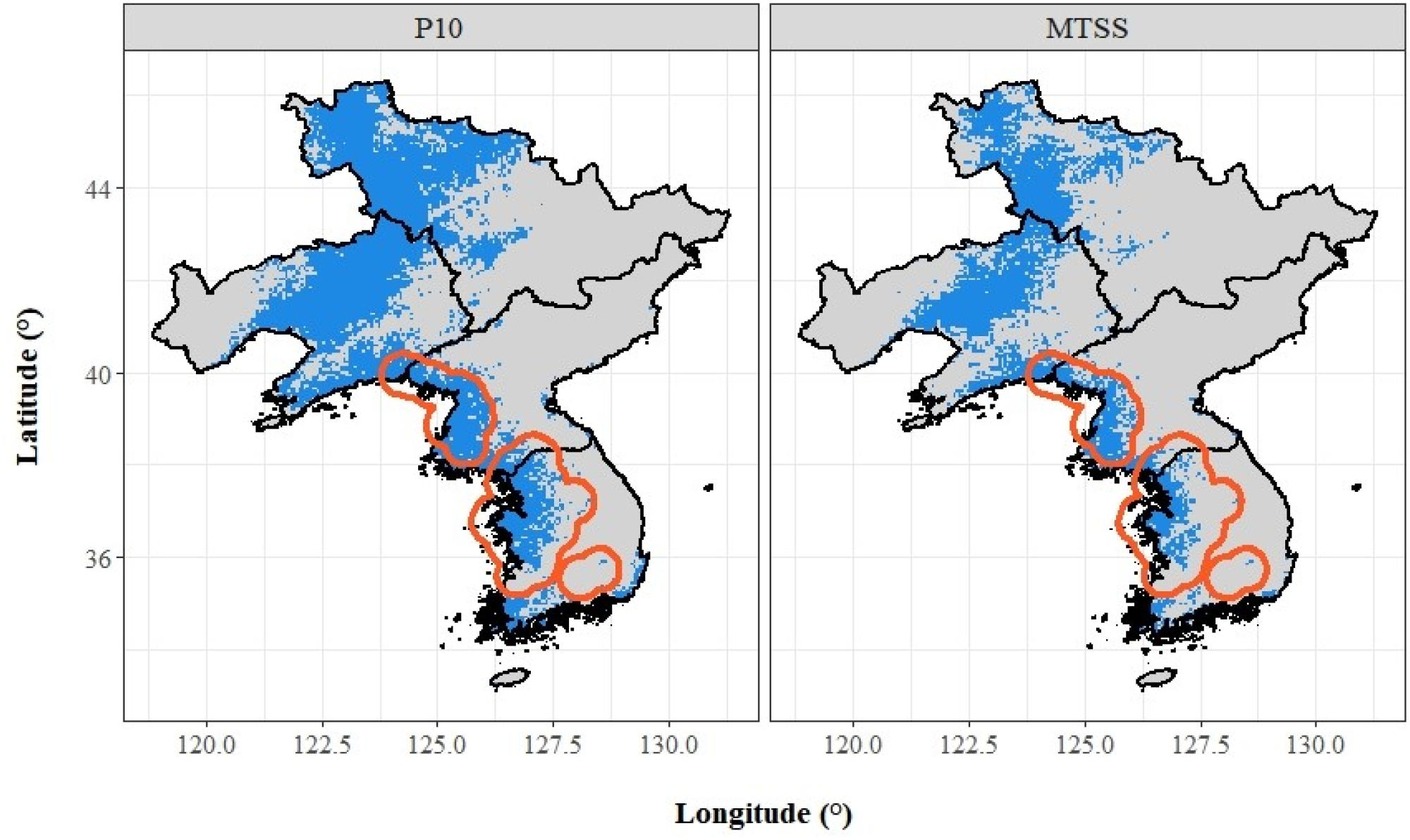
Binary presence/absence maps for *Pelophylax chosenicus*. Generated from MaxEnt model outputs by applying one “liberal” threshold (= 0.2578; 10 percentile training presence; P10) and one “strict” threshold (= 0.4154; Maximum Test Sensitivity plus Specificity; MTSS). While both thresholds generally predicted similar areas of presence both within the calibration area (circular polygon) and across the entire study area, the more liberal P10 threshold predicted broader areas of presence.

## Discussion

Here we report for the first time the presence of *Pelophylax chosenicus* in China, identified through molecular tools. Here, we do not clarify the relationship between *P. chosenicus* and its sister clade *P. plancyi*, and either of the two clades within *P. chosenicus* (Min et al. 2008) could be the one found at the site. We then identify suitable habitats throughout eastern DPR Korea using landscape modeling. Finally, we confirm reports of two inland populations in R Korea, one south of Daegu and the other in the vicinity of Hapcheon. The southernmost population reported in Buan is likely not the southernmost one based on modelling results, despite a distribution generally matching that of the Korean endemic *Dryophytes flaviventris* (Borzée et al. 2020a). The presence of the two inland populations in R Korea is also in agreement with the modeling results and confirm the likely presence of other inland populations throughout the Korea Peninsula, and maybe further north in Liaoning (Figure 1). This is especially likely in view of the modeling results predicting the presence of the species in the plains west of the Bohai Bay. However, further surveys are needed as the species would have to have dispersed across the hills between Dandong and Shenyang, a range delineation boundary for some species, such as *Rana huanrenensis* (IUCN SSC Amphibian Specialist Group 2019b). This type of movement is not impossible as dispersal across the Yellow Sea happened in some species, including *Rana coreana* (IUCN SSC Amphibian Specialist Group 2019a). Both surveys and modelling results support the absence of *P. chosenicus* on the east coast of the Korean Peninsula and further north, in line with the data already published (NIBR 2019) and available through the citizen science platform iNaturalist (https://www.inaturalist.org/observations?place_id=any&subview=map&taxon_id=66318), a credible platform for amphibian species identification (Borzée et al. 2019a). Further surveys in eastern Korea, Jilin and the Russian Primorye are thus unlikely to detect the presence of the species, but additional surveys are needed in Jeolla Province in R Korea, along the whole west coast of DPR Korea, and in all the lowlands of Liaoning in China.

In terms of landscape use, *P. chosenicus* was found to be significantly more often present in croplands, deciduous broadleaf forests and urban areas. It is important to understand the link between the drivers of settlement for historical human populations, their transformation into current urban area, and the match with the lowlands used by species, similarly to *Dryophytes suweonensis* (Borzée et al. 2017b), but also amphibians in general (Small and Cohen 2004), and the connection with high biodiversity (Dambrine et al. 2007). The sites with putative absence were generally agricultural landscapes, followed by deciduous broadleaf forest, vegetation mosaics and urban areas. The overlap in the landscape features where the species was found and putatively absent is representative of the broad scale of the sampling effort, within and outside of the range of the species, and highlights that landscape variables are not limiting the distribution of the species. In contrast, the distribution of *P. chosenicus* is likely restricted by other variables such as elevation or climate, similarly to other anuran species (Otto et al. 2007). The significant difference in presence and putative absence in relation to rice paddies is important as the species relies on this type of landscape for its survival, as expected due to the high rate of conversion of natural wetlands into agricultural wetlands in R Korea (Lee and Miller-Rushing 2014; Park et al. 2021b; Borzée et al. 2019c), but it also relies on other types of landscapes, and even wetlands in urban areas (Sung et al. 2007; Ra et al. 2008b; Ra et al. 2010).

In general, our modelling results were highly supported, and the empirical models were better supported than the null models, highlighting the validity of our results. In addition, the two binary thresholds used did not show extensive variations in the suitable habitat for the species. Isothermality was the most important variables, representative of daily temperature fluctuations in comparison with yearly variations, and highlighting that the range of the species was not determined by landscape use only, similarly to the Chinese giant salamander as its range is also restricted by isothermality (Zhang et al. 2020). Slope was the second most important variable (<15°) for the occurrence of the species, generally matching with lowlands in our focal landscape, a variable also associated with the occurrence of the syntopic *D. suweonensis* and *D. flaviventris* (Borzée et al. 2020a), and also associated with the Chinese giant salamander (Zhang et al. 2020). Next in the list of important variables was the precipitation of the driest month, with the presence of the species being associated with rain regimes matching with light rain (> 25mm), followed by a mean temperature of the wettest quarter (>22°C), matching with relatively high temperature for temperate areas, and the breeding season of the species (Seo et al. 2014). Finally, vegetation cover with cultivated landscape over >40%, herbaceous vegetation cover approximately below 52% and cover of urban areas approximately below 85% were related to the presence of the species. These last variables also highlight that while the species can live in agricultural wetlands, likely due to their ancestral presence in habitats being transformed, they are also present in other areas.

The strong overlap with urban areas could be related to the ability of the species to cope with development, although it is more likely related to the expectation that *P. chosenicus* can be relatively long lived (15 years for the closely related *P. nigromaculatus*, Kashiwagi et al. 2005), in connection with the fact that development is recent (Lee and Miller-Rushing 2014; Lee et al. 2018); and that individuals in urban areas are remnant populations as the species is sensitive to pollution (Byeon 2010; Borzée et al. 2018c).

While our results can be seen in a positive light as we define a broader range than previously known for *P. chosenicus*, these are not genuine improvements in the status of the species, but additional information on a species that is listed as Vulnerable and decreasing in the IUCN Red List of Threatened Species (IUCN SSC Amphibian Specialist Group 2021). The species faces multiple threats (Cheong et al. 2009; Ra and Park 2011; Ra et al. 2016), but habitat loss remains the main threat as most low elevation wetlands have been converted to rice agriculture, and are now being transformed into dry agriculture or tertiary production for economic reasons (e.g. in the boundaries of Seoul; Park et al. 2021b). However, the threats need to be better understood as the species is not strictly bound to rice paddies, unlike other threatened species in the landscape (NIBR 2019), but it is sensitive to agrochemical pollution (Borzée et al. 2018c). In addition, the species is under threat due to climate change both because of the shift in climate at current natural habitats (Gerick et al. 2014; Kim et al. 2021), but also because of the delay in the flooding of rice paddies where the species breeds in agricultural wetlands (Chuang et al. 2018). It is important to note that conservation efforts are ongoing in R Korea, with a translocation program initiated by the National Institute of Ecology of Korea (Park et al. 2021a), and several conservation plans and recommendation available (Yang and Koo 2016; Ra et al. 2019; Yoo et al. 2019). In addition, the species is not likely to be threatened in PDR Korea. Further surveys are needed in China to better understand the distribution of the species, and the threats in the nation.

## Acknowledgement

We are grateful to Yikweon Jang for the help provided over the years and to Desiree Andersen for the help during fieldwork in China. This work was supported by the Foreign Youth Talent Program (QN2021014013L) from the Ministry of Science and Technology to AB.

## Authors contribution

Conceptualization: AB; Methodology: AB, YS, YB, SNO; Software: AB, YS, SNO; Validation: AB, YS, YB, SNO; Formal analysis: AB, YS, SNO; Resources: AB; Writing - Original Draft: AB, YS, SNO; Writing - Review & Editing: AB, YS, YB, SNO; Funding acquisition: AB.

## Data availability

All the data used in this study can be found in the Supplementary files 1 (landscape modelling) and 2 (molecular analyses), and in the Global Biodiversity Information Facility (gbif.org; DOI: 10.15468/dl.6kvymf). The complete MaxEnt distribution modeling workflow in R is available from GitHub (https://github.com/yucheols/Pelophylax_chosenicus_SDM). The sequences generated from this study has been deposited to the GenBank (https://www.ncbi.nlm.nih.gov/) under the accession numbers ON054286 through ON054290.

## References

Allouche O, Tsoar A, and Kadmon R (2006) Assessing the accuracy of species distribution models: prevalence, kappa and the true skill statistic (TSS). Journal of Applied Ecology 43: 1223–1232. 10.1111/j.1365-2664.2006.01214.x

Amatulli G, Domisch S, Tuanmu M-N, Parmentier B, Ranipeta A, Malczyk J, and Jetz W (2018) A suite of global, cross-scale topographic variables for environmental and biodiversity modeling. Scientific Data 5: 180040. 10.1038/sdata.2018.40

Araújo MB, Anderson RP, Barbosa AM, Beale CM, Dormann CF, Early R, Garcia RA, Guisan A, Maiorano L, and Naimi B (2019) Standards for distribution models in biodiversity assessments. Science Advances 5: eaat4858. 10.1126/sciadv.aat4858

Bae Y, Park J, Othman SN, Jang Y, and Borzée A (2022) Record of invasive Rana huanrenensis Fei, Ye, and Huang, 1990 and Pelophylax nigromaculatus (Hallowell, 1861) on Ulleung Island, Republic of Korea. BioInvasion Records 11: 278–286. 10.3391/bir.2022.11.1.28

Barber RA, Ball SG, Morris RKA, and Gilbert F (2022) Target-group backgrounds prove effective at correcting sampling bias in Maxent models. Diversity and Distributions 28: 128–141. 10.1111/ddi.13442

Başkale E, and Çapar D (2016) Detection probability and habitat selection of the Beyşehir Frog, Pelophylax caralitanus (Arikan 1988), in southwestern Anatolia, Turkey. Russian Journal of Herpetology 23: 205–214.

Bivand R, Keitt T, and Rowlingson B (2021) rgdal: Bindings for the ‘Geospatial’ Data Abstraction Library. R package version 1.5-23: https://CRAN.R-project.org/package=rgdal.

Bohl CL, Kass JM, and Anderson RP (2019) A new null model approach to quantify performance and significance for ecological niche models of species distributions. Journal of Biogeography 46: 1101–1111. 10.1111/jbi.13573

Borzée A, Andersen D, and Jang Y (2018a) Population trend inferred from aural surveys for calling anurans in Korea. PeerJ 6: e5568. 10.7717/peerj.5568

Borzée A, Bae YH, Groffen J, and Jang Y (2017a) Pelophylax chosenicus (Gold-spotted Pond Frog). Herpetological Review 48: 79–80.

Borzée A, Bae YH, and Seliger B (2018b) Pelophylax chosenicus (Gold-spotted Pond Frog). Herpetological Review 49: 709.

Borzée A, Baek HJ, Lee CH, Kim DY, Song J-Y, Suh J-H, Jang Y, and Min M-S (2019a) Scientific publication of georeferenced molecular data as an adequate guide to delimit the range of Korean Hynobius salamanders through citizen science. Acta herpetologica 14: 27–33. 10.13128/Acta_Herpetol-24102

Borzée A, Kim K, Heo K, Jablonski PG, and Jang Y (2017b) Impact of land reclamation and agricultural water regime on the distribution and conservation status of the endangered Dryophytes suweonensis. PeerJ 5: e3872. 10.7717/peerj.3872

Borzée A, Kosch TA, Kim M, and Jang Y (2017c) Introduced bullfrogs are associated with increased Batrachochytrium dendrobatidis prevalence and reduced occurrence of Korean treefrogs. PLOS One 12: e0177860. 10.1371/journal.pone.0177860

Borzée A, Kyong CN, Kil HK, and Jang Y (2018c) Impact of water quality on the occurrence of two endangered Korean anurans: Dryophytes suweonensis and Pelophylax chosenicus. Herpetologica 74: 1–7. 10.1655/Herpetologica-D-17-00011

Borzée A, Litvinchuk SN, Ri K, Andersen D, Nam TY, Jon GH, Man HS, Choe JS, Kwon S, Othman SN, Messenger K, Bae Y, Shin Y, Kim A, Maslova I, Luedtke J, Hobin L, Moores N, Seliger B, Glenk F, and Jang Y (2021) Update on distribution and conservation status of amphibians in the Democratic People’s Republic of Korea: conclusions based on field surveys, environmental modelling, molecular analyses and call properties. Animals 11: 2057. 10.3390/ani11072057

Borzée A, Messenger KR, Chae S, Andersen D, Groffen J, Kim YI, An J, Othman S, Ri K, Nam TY, Bae Y, Ren J-L, Li J-T, Chuang M-F, Yi Y, Yucheol Shin, Kwon T, Jang Y, and Min M-S (2020a) Yellow sea mediated segregation between North East Asian Dryophytes species. Plos One 15: e0234299. 10.1371/journal.pone.0234299

Borzée A, Oh S, Sin E, and Jang Y (2020b) Spring voices in Korean rice fields: the effect of abiotic variables and syntopic calls on the calling activity of the treefrog Dryophytes suweonensis. Asian Herpetological Research 11: 335–341. 10.16373/j.cnki.ahr.190046

Borzée A, Ren J, Li J-T, Groffen J, Jang Y, and Messenger RK (2019b) Microhabitat segregation between Black-spotted Pond Frogs, Pelophylax nigromaculatus, and Gold-striped Pond Frogs, P. plancyi (Anura: Ranidae). Reptiles & Amphibians: Conservation and Natural History 26: 119–120.

Borzée A, Struecker M-Y, Yi Y, Kim D, and Kim H (2019c) Time for Korean wildlife conservation. Science 363: 1161–1162. 10.1126/science.aaw9023

Byeon CW (2010) Water purification and ecological restoration effects of the Keumeo stream sustainable structured wetland biotop (SSB) System established on the floodplain of Kyungan stream. Journal of the Korean Society of Environmental Restoration Technology 13: 23–35.

Cheong SW, Sung HC, Park DS, and Park SR (2009) Population viability analysis of a Gold-spotted Pond Frog (Rana chosenica) population: implications for effective conservation and re-introduction. Korean Journal of Environmental Biology 27: 73–81.

Chuang MF, Borzée A, and Jang Y (Year). “Impact of environmental variables on the breeding phenology of a South Korean treefrog, Dryophytes suweonensis”, in: International Long Term Ecological Research Network).

Dambrine É, Dupouey JL, Laüt L, Humbert L, Thinon M, Beaufils T, and Richard H (2007) Present forest biodiversity patterns in France related to former Roman agriculture. Ecology 88: 1430–1439. 10.1890/05-1314

Di Cola V, Broennimann O, Petitpierre B, Breiner FT, D’amen M, Randin C, Engler R, Pottier J, Pio D, Dubuis A, Pellissier L, Mateo RG, Hordijk W, Salamin N, and Guisan A (2017) ecospat: an R package to support spatial analyses and modeling of species niches and distributions. Ecography 40: 774–787. 10.1111/ecog.02671

Do MS, Son SJ, Choi G, Yoo N, Koo KS, and Nam HK (2021) Anuran community patterns in the rice fields of the mid-western region of the Republic of Korea. Global Ecology and Conservation 26: e01448. 10.1016/j.gecco.2020.e01448

Eom JH, Lee JH, Ra NY, and Park DS (2007) Preferred feeding sites and prey of the adult Gold-spotted pond frog, Rana plancyi chosenica. Journal of Ecology and Environment 30: 357–361. 10.5141/JEFB.2007.30.4.357

Fei L, Changyuan Y, and Jianping J (2012) Colored atlas of Chinese amphibians and their distributions. Sichuan science and technology press, Chendu, People’s Republic of China.

Fick SE, and Hijmans RJ (2017) WorldClim 2: new 1km spatial resolution climate surfaces for global land areas. International Journal of Climatology 37: 4302–4315. 10.1002/joc.5086

Fletcher Jr RJ, Hefley TJ, Robertson EP, Zuckerberg B, Mccleery RA, and Dorazio RM (2019) A practical guide for combining data to model species distributions. Ecology 100: e02710. 10.1002/ecy.2710

Frost DR, Grant T, Faivovich JN, Bain RH, Haas A, Haddad CLFB, D. Sá RO, Channing A, Wilkinson M, Donnellan SC, Raxworthy CJ, Campbell JA, Blotto BL, Moler P, Drewes RC, Nussbaum RA, Lynch JD, Green DM, and Wheeler WC (2006) The Amphibian Tree of Life. Bulletin of the American Museum of Natural History 297: 1–291. 10.1206/0003-0090(2006)297[0001:TATOL]2.0.CO;2.hdl:2246/5781

Garcia VOS, Ivy C, and Fu J (2017) Syntopic frogs reveal different patterns of interaction with the landscape: A comparative landscape genetic study of Pelophylax nigromaculatus and Fejervarya limnocharis from central China. Ecology and evolution 7: 9294–9306. 10.1002/ece3.3459

Gerick AA, Munshaw RG, Palen WJ, Combes SA, and O’regan SM (2014) Thermal physiology and species distribution models reveal climate vulnerability of temperate amphibians. Journal of Biogeography 41: 713–723. 10.1111/jbi.12261

Hijmans RJ (2021) raster: Geographic Data Analysis and Modeling. R package version 3.4-13: https://CRAN.R-project.org/package=raster. https://CRAN.R-project.org/package=raster

Hijmans RJ, Phillips S, Leathwick J, and Elith J (2020) dismo: Species Distribution Modeling. R package version 1.3-3: https://CRAN.R-project.org/package=dismo.

Hofman S, Pabijan M, Osikowski A, Litvinchuk SN, and Szymura JM (2016) Phylogenetic relationships among four new complete mitogenome sequences of Pelophylax (Amphibia: Anura) from the Balkans and Cyprus. Mitochondrial DNA 27: 3434–3437. 10.3109/19401736.2015.1025266

Huelsenbeck JP, and Ronquist F (2001) MRBAYES: Bayesian inference of phylogenetic trees. Bioinformatics 17: 754–755.

Iucn Ssc Amphibian Specialist Group (2019a) Rana coreana. The IUCN Red List of Threatened Species e.T89108544A110101367. 10.2305/IUCN.UK.2019-1.RLTS.T89108544A110101367.en.

Iucn Ssc Amphibian Specialist Group (2019b) Rana huanrensis. The IUCN Red List of Threatened Species 2019 e.T58619A63855773. 10.2305/IUCN.UK.2019-1.RLTS.T58619A63855773.en.

Iucn Ssc Amphibian Specialist Group (2021) Pelophylax chosenicus. The IUCN Red List of Threatened Species e.T58577A110101963. 10.2305/IUCN.UK.2021-3.RLTS.T58577A110101963.en

Jeong TJ, Jun J, Han S, Kim HT, Oh K, and Kwak M (2013) DNA barcode reference data for the Korean herpetofauna and their applications. Molecular Ecology Resources 13: 1019–1032. 10.1111/1755-0998.12055

Jiang J, and Zhou K (2005) Phylogenetic relationships among Chinese ranids inferred from sequence data set of 12S and 16S rDNA. Herpetological Journal 15: 1–8. 10.1142/9789812703576

Jiang L, Zhao L, Liu Y, Leng Z, Zhao L, and Ruan Q (2017) The complete mitochondrial genome sequence of the Dark-spotted frog Pelophylax nigromaculatus (Amphibia, Anura, Ranidae). Mitochondrial DNA Part A: DNA Mapping, Sequencing, and Analysis 28: 236–237. 10.3109/19401736.2015.1115857

Kashiwagi K, Shinkai T, Kajii E, and Kashiwagi A (2005) The effects of reactive oxygen species on amphibian aging. Comparative Biochemistry and Physiology Part C: Toxicology & Pharmacology 140: 197–205. 10.1016/j.cca.2005.02.001

Kass JM, Muscarella R, Galante PJ, Bohl CL, Pinilla-Buitrago GE, Boria RA, Soley-Guardia M, and Anderson RP (2021) ENMeval 2.0: Redesigned for customizable and reproducible modeling of species’ niches and distributions. Methods in Ecology and Evolution 12: 1602–1608. 10.1111/2041-210X.13628

Kearse M, Moir R, Wilson A, Stones-Havas S, Cheung M, Sturrock S, Buxton S, Cooper A, Markowitz S, and Duran C (2012) Geneious Basic: an integrated and extendable desktop software platform for the organization and analysis of sequence data. Bioinformatics 28: 1647–1649. 10.1093/bioinformatics/bts199

Kim H-T, Kim H, Jeon G, and Kim D (2019) Arrow guide of amphibians and reptiles. Econature, Seoul, Republic of Korea.

Kim H, Adhikari P, Chang M, and Seo C (2021) Potential distribution of amphibians with different habitat characteristics in response to climate change in South Korea. Animals 11: 2185. 10.3390/ani11082185

Kobayashi T, Tateishi R, Alsaaideh B, Sharma RC, Wakaizumi T, Miyamoto D, Bai X, Long BD, Gegentana G, and Maitiniyazi A (2017) Production of global land cover data - GLCNMO2013. Journal of Geography and Geology 9: 1–15. 10.5539/jgg.v9n3p1

Komaki S, Igawa T, Lin SM, Tojo K, Min MS, and Sumida M (2015) Robust molecular phylogeny and palaeodistribution modelling resolve a complex evolutionary history: glacial cycling drove recurrent mtDNA introgression among Pelophylax frogs in East Asia. Journal of Biogeography 42: 2159–2171. 10.1111/jbi.12584

Lanfear R, Frandsen PB, Wright AM, Senfeld T, and Calcott B (2017) PartitionFinder 2: new methods for selecting partitioned models of evolution for molecular and morphological phylogenetic analyses. Molecular Biology and Evolution 34: 772–773. 10.1093/molbev/msw260

Lee JK, Chung O-S, Park J-Y, Kim H-J, Hur W-H, Kim S-H, and Kim J-H (2018) Effects of the Saemangeum Reclamation Project on migratory shorebird staging in the Saemangeum and Geum Estuaries, South Korea. Bird Conservation International 28: 238–250. 10.1017/S0959270916000605

Lee S-D, and Miller-Rushing AJ (2014) Degradation, urbanization, and restoration: a review of the challenges and future of conservation on the Korean Peninsula. Biological Conservation 176: 262–276. 10.1016/j.biocon.2014.05.010

Liu K, Wang F, Chen W, Tu L, Min M-S, Bi K, and Fu J (2010) Rampant historical mitochondrial genome introgression between two species of green pond frogs, Pelophylax nigromaculatus and P. plancyi. BMC evolutionary biology 10: 201. 10.1186/1471-2148-10-201

Manne LL, and Pimm SL (2001) Beyond eight forms of rarity: which species are threatened and which will be next? Animal Conservation 4: 221–229. 10.1017/S1367943001001263

Min M-S, Park SK, Che J, Park DS, and Lee H (2008) Genetic diversity among local populations of the gold-spotted pond frog, Rana plancyi chosenica (Amphibia: Ranidae), assessed by mitochondrial cytochrome b gene and control region sequences. Animal Systematics, Evolution and Diversity 24: 25–32. 10.5635/kjsz.2008.24.1.025

Nakanishi K, Honma A, Furukawa M, Takakura K-I, Fujii N, Morii K, Terasawa Y, and Nishida T (2020) Habitat partitioning of two closely related pond frogs, Pelophylax nigromaculatus and Pelophylax porosus brevipodus, during their breeding season. Evolutionary Ecology 34: 855–866. 10.1007/s10682-020-10061-1

Nibr (2019) Red Data Book of Republic of Korea. Volume 2: Amphibians and Reptiles. National Institute of Biological Resources, Ministry of Environment, Incheon, Republic of Korea.

Nishioka M (1972) Nucleo-cytoplasmic hybrids between Rana brevipoda and Rana plancyi chosenica. Scientific report of the Laboratory for Amphibian Biology of Hiroshima University 1: 259–275. 10.15027/287

Nishioka M, and Okumoto H (1983) Reproductive capacity and progeny of amphidiploids between Rana nigromaculata and Rana plancyi chosenica. Scientific report of the Laboratory for Amphibian Biology of Hiroshima University 6: 141–181. 10.15027/352

Okada Y (1926) A study on the distribution of tailess batrachians of Japan. Annotationes Zoologicae Japonenses 11: 137–144.

Okada Y (1928) Frogs in Korea. Journal of the Chosen Biological Society 6: 15–46.

Okada Y (1931) The tailless bactracians of the Japanese Empire. Journal of the Imperial Agricultural Experiment Station, Nishigahara, Tokyo 29: 1–215.

Otto CR, Snodgrass JW, Forester DC, Mitchell JC, and Miller RW (2007) Climatic variation and the distribution of an amphibian polyploid complex. Journal of Animal Ecology 76: 1053–1061. 10.1111/j.1365-2656.2007.01300.x

Park C, Kwon K, Yoo N, Lee J, Kang D, Park J, Yoo J, Kim K, and Yoon J (2021a) Post-release monitoring after reintroduction of captive-reared Korean endangered frog, Pelophylax chosenicus. Proceedings of the National Institute of Ecology of the Republic of Korea 2: 114–119. 10.22920/PNIE.2021.2.2.114

Park D-S, Park S-R, and Sung H-C (2009) Colonization and extinction patterns of a metapopulation of Gold-spotted Pond Frogs, Rana plancyi chosenica. Journal of Ecology and Environment 32: 103–107. 10.5141/JEFB.2009.32.2.103

Park I-K, Park D, and Borzée A (2021b) Defining conservation requirements for the Suweon Treefrog (Dryophytes suweonensis) using species distribution models. Diversity 13: 69. 10.3390/d13020069

Petitot M, Manceau N, Geniez P, and Besnard A (2014) Optimizing occupancy surveys by maximizing detection probability: application to amphibian monitoring in the Mediterranean region. Ecology and evolution 4: 3538–3549. 10.1002/ece3.1207

Pyron RA (2014) Biogeographic analysis reveals ancient continental vicariance and recent oceanic dispersal in amphibians. Systematic Biology 63: 779–797. 10.1093/sysbio/syu042

R Core Team (2021) R: A language and environment for statistical computing. R Foundation for Statistical Computing, Vienna, Austria.

Ra N-Y, Bae S, Min S-H, Son H, Baek S-J, and Geum J-W (2019) Recovery process of the Endangered Gold-Spotted Pond Frog(Pelophylax chosenicus) in the Du-ung Wetland. The Korean Research Society of Herpetologists 18: https://www.earticle.net/Article/A361215.

Ra N-Y, Sung H-C, and Park D (2008a) Characteristics of Rana plancyi chosenica habitats and potential factors involved in the decline of the species. The Korean Research Society of Herpetologists 8: 14.

Ra N-Y, Sung H-C, and Park D (2016) Current status of the gold-spotted pond frog, Rana plancyi chosenica, and its habitat distributions in Korea. The Korean Research Society of Herpetologists 13–14. https://www.earticle.net/Article/A281134

Ra NY, and Park D (2011) Saving the Gold-spotted pond frog in South Korea. FrogLog 98: 15.

Ra NY, Park D, Cheong S, Kim NS, and Sung HC (2010) Habitat associations of the endangered Gold-spotted pond frog (Rana chosenica). Zoological Science 27: 396–401. 10.2108/zsj.27.396

Ra NY, Sung HC, Cheong S, Lee JH, Eom J, and Park D (2008b) Habitat use and home range of the endangered Gold-spotted pond frog (Rana chosenica). Zoological science 25: 894–903. 10.2108/zsj.25.894

Rambaut A, Drummond AJ, Xie D, Baele G, and Suchard MA (2018) Posterior summarization in Bayesian phylogenetics using Tracer 1.7. Systematic Biology 67: 901–904. 10.1093/sysbio/syy032

Richardson DM, and Whittaker RJ (2010) Conservation biogeography–foundations, concepts and challenges. Diversity and Distributions 16: 313–320. 10.1111/j.1472-4642.2010.00660.x

Ryu SH, and Hwang UW (2011) Complete mitochondrial genome of the Seoul frog Rana chosenica (Amphibia, Ranidae): Comparison of R. chosenica and R. plancyi. Mitochondrial DNA 22: 53–54. 10.3109/19401736.2011.603313

Seo M, Lee B, Park C, Oh H, Kim H, Lee K, Kang C, Gil H, and Park J (2014) Threatened wildlife at a glance. National Institute of Biological Resources, Incheon, Republic of Korea.

Shannon FA (1956) The Reptiles and Amphibians of Korea. Herpetologica 12: 22–49.

Shim JH (2003) Study on the In Extu and restoration strategy planning for the protected wildlife Anura (Rana plancyi chosenica Okada) in Korea. Korean Journal of Nature Conservation 1: 63–74. 10.30960/kjnc.2003.1.4.63

Shin Y, Jang Y, Allain SJR, and Borzée A (2020) Catalogue of herpetological specimens of the Ewha Womans University Natural History Museum (EWNHM), Republic of Korea. ZooKeys 965: 103–139. 10.3897/zookeys.965.52976.

Shipley BR, Bach R, Do Y, Strathearn H, Mcguire JL, and Dilkina B (2022) megaSDM: integrating dispersal and time-step analyses into species distribution models. Ecography in press. 10.1111/ecog.05450

Sievers F, Wilm A, Dineen D, Gibson TJ, Karplus K, Li W, Lopez R, Mcwilliam H, Remmert M, Söding J, Thompson JD, and Higgins DG (2011) Fast, scalable generation of high-quality protein multiple sequence alignments using Clustal Omega. Molecular Systems Biology 7: 539. 10.1038/msb.2011.75

Small C, and Cohen JE (2004) Continental Physiography, Climate, and the Global Distribution of Human Population. Current Anthropology 45: 269–277.

Soberón J, and Peterson AT (2005) Interpretation of models of fundamental ecological niches and species’ distributional areas. Biodiversity Informatics 2: 1–10. 10.17161/bi.v2i0.4

Son E, and Borzée A (2020) Pelophylax chosenicus (Gold-spotted Pond Frog), Geographic distribution note. Herpetological Review 51: 134.

Sumida M, Kondo Y, Kanamori Y, and Nishioka M (2002) Inter-and intraspecific evolutionary relationships of the rice frog Rana limnocharis and the allied species R. cancrivora inferred from crossing experiments and mitochondrial DNA sequences of the 12S and 16S rRNA genes. Molecular Phylogenetics and Evolution 25: 293–305. 10.1016/S1055-7903(02)00243-9

Sung HC, Cha SM, Cheong SW, Park DS, and Park SR (2007) Monitoring local populations and breeding migration patterns of the Gold-spotted pond frog, Rana chosenica. Journal of Ecology and Environment 30: 121–126. 10.5141/JEFB.2007.30.2.121

Tokumoto Y, Ramamonjisoa N, Zheng XJ, and Natuhara Y (2019) 16S rDNA sequences of 8 frog species and rhod sequences of two Pelophylax spp. in Aichi prefecture. Japan. Bulletin of Nagoya Biodiversity Center 6: 15–22.

Tuanmu M-N, and Jetz W (2014) A global 1-km consensus land-cover product for biodiversity and ecosystem modeling. Global Ecology and Biogeography 23: 1031–1045. 10.1111/geb.12182

Vignali S, Barras A, Arlettaz R, and Braunisch V (2020) SDMtune: An R package to tune and evaluate species distribution models. Ecology and Evolution 10: 11488–11506. 10.1002/ece3.6786

Wallace AR (1876) The geographical distribution of animals. Harvard University Press, Cambridge, USA.

Wang S, Fan L, Liu C, Li J, Gao X, Zhu W, and Li Y (2017) The origin of invasion of an alien frog species in Tibet, China. Current Zoology 63: 615–621. 10.1093/cz/zow117

Wang S, Liu C, Zhu W, Gao X, and Li Y (2016) Tracing the Origin of the Black-spotted Frog, Pelophylax nigromaculatus, in the Xinjiang Uyghur Autonomous Region. Asian Herpetological Research 7: 69–74. 10.16373/j.cnki.ahr.150071

Wang Z, Zeng J, Meng W, Lohman DJ, and Pierce NE (2021) Out of sight, out of mind: public and research interest in insects is negatively correlated with their conservation status. Insect Conservation and Diversity 14: 700–708. 10.1111/icad.12499

Warren DL, and Seifert SN (2011) Ecological niche modeling in Maxent: the importance of model complexity and the performance of model selection criteria. Ecological Applications 21: 335–342. 10.1890/10-1171.1

Yang DS, and Koo BH (2016) A study on the improvement plan for a habitat of “Gold-spotted pond frog (Pelophylax chosenicus)” in danger of regional extinction in the urban area - case on the abandoned railroad site on Su-in line. Journal of the Korean Society of Environmental Restoration Technology 19: 95–107. 10.13087/kosert.2016.19.2.95

Yang SY, Kim JB, Min MS, Suh JH, and Kang YJ (2000) Monograph of Korean Amphibia. Academy Book, Seoul.

Yoo N, Do MS, Nam HK, Choi G, Son SJ, and Yoo JC (2019) Habitat characteristics of anuran species inhabiting rice fields of western mid-south Korea - in the case of Daeho reclamation agricultural land by farming practices. Korean Journal of Ecology and Environment 52: 366–377. 10.11614/KSL.2019.52.4.366

Yoon IB, Kim JI, and Yang SY (1998) Study on the food habits of Rana nigromaculata Hallowell and Rana plancyi chosenica Okada (Salientia; Ranidae) in Korea. Korean Journal of Environmental Biology 16: 69–76.

Yuan Z-Y, Zhou W-W, Chen X, Poyarkov Jr NA, Chen H-M, Jang-Liaw N-H, Chou W-H, Matzke NJ, Iizuka K, and Min M-S (2016) Spatiotemporal diversification of the true frogs (genus Rana): a historical framework for a widely studied group of model organisms. Systematic biology 65: 824–842. 10.1093/sysbio/syw055

Zhang Z, Mammola S, Liang Z, Capinha C, Wei Q, Wu Y, Zhou J, and Wang C (2020) Future climate change will severely reduce habitat suitability of the Critically Endangered Chinese giant salamander. Freshwater Biology 65: 971–980. 10.1111/fwb.13483

