## Supplementary File 1 for "Korean endemic species no more: on the occurrence of *Pelophylax chosenicus* in China"

Supplementary File 1. Dataset on presence and absence points for *Pelophylax chosenicus* used in this study. The dataset includes all landscape types, but also a binary encoded occurrence in rice paddies, which was used independently for the analyses. Binary encoded such as 0 = absence and 1 = presence.

| Longitude | Latitude | Binary encoded presence | Land feature | Binary encoded presence in the forest land feature | Binary encoded presence in the agriculture land feature | Binary encoded presence in the shrub land feature | Binary encoded presence in the grassland land feature | Binary encoded presence in the urban land feature | Binary encoded presence in the water land feature | Binary encoded for presence in rice paddies |
| --- | --- | --- | --- | --- | --- | --- | --- | --- | --- | --- |
| 126.8827 | 36.30347 | 1 | Broadleaf deciduous forest | 1 | 0 | 0 | 0 | 0 | 0 | 0 |
| 127.3117 | 36.85113 | 1 | Broadleaf deciduous forest | 1 | 0 | 0 | 0 | 0 | 0 | 0 |
| 127.3087 | 36.86886 | 1 | Broadleaf deciduous forest | 1 | 0 | 0 | 0 | 0 | 0 | 0 |
| 127.4047 | 36.97209 | 1 | Broadleaf deciduous forest | 1 | 0 | 0 | 0 | 0 | 0 | 0 |
| 127.6696 | 37.04482 | 1 | Broadleaf deciduous forest | 1 | 0 | 0 | 0 | 0 | 0 | 0 |
| 127.3265 | 37.66782 | 1 | Broadleaf deciduous forest | 1 | 0 | 0 | 0 | 0 | 0 | 0 |
| 126.1857 | 36.77429 | 1 | Broadleaf deciduous forest | 1 | 0 | 0 | 0 | 0 | 0 | 0 |
| 126.6904 | 37.95774 | 1 | Broadleaf deciduous forest | 1 | 0 | 0 | 0 | 0 | 0 | 0 |
| 126.879 | 36.39289 | 1 | Broadleaf deciduous forest | 1 | 0 | 0 | 0 | 0 | 0 | 0 |
| 126.9685 | 36.85628 | 1 | Broadleaf deciduous forest | 1 | 0 | 0 | 0 | 0 | 0 | 0 |
| 127.044 | 38.19683 | 1 | Broadleaf deciduous forest | 1 | 0 | 0 | 0 | 0 | 0 | 0 |
| 127.2034 | 36.45207 | 1 | Broadleaf deciduous forest | 1 | 0 | 0 | 0 | 0 | 0 | 0 |
| 127.4153 | 36.78956 | 1 | Broadleaf deciduous forest | 1 | 0 | 0 | 0 | 0 | 0 | 0 |
| 127.5907 | 36.68132 | 1 | Broadleaf deciduous forest | 1 | 0 | 0 | 0 | 0 | 0 | 0 |
| 127.9463 | 37.20683 | 1 | Broadleaf deciduous forest | 1 | 0 | 0 | 0 | 0 | 0 | 0 |
| 126.4858 | 33.26216 | 0 | Broadleaf deciduous forest | 1 | 0 | 0 | 0 | 0 | 0 | 0 |
| 127.9853 | 35.66556 | 0 | Broadleaf deciduous forest | 1 | 0 | 0 | 0 | 0 | 0 | 0 |
| 127.8595 | 35.6714 | 0 | Broadleaf deciduous forest | 1 | 0 | 0 | 0 | 0 | 0 | 0 |
| 128.0841 | 35.67283 | 0 | Broadleaf deciduous forest | 1 | 0 | 0 | 0 | 0 | 0 | 0 |
| 127.8118 | 35.75323 | 0 | Broadleaf deciduous forest | 1 | 0 | 0 | 0 | 0 | 0 | 0 |
| 128.1175 | 35.76649 | 0 | Broadleaf deciduous forest | 1 | 0 | 0 | 0 | 0 | 0 | 0 |
| 127.8843 | 35.77311 | 0 | Broadleaf deciduous forest | 1 | 0 | 0 | 0 | 0 | 0 | 0 |
| 128.0875 | 35.77506 | 0 | Broadleaf deciduous forest | 1 | 0 | 0 | 0 | 0 | 0 | 0 |
| 127.9786 | 35.79641 | 0 | Broadleaf deciduous forest | 1 | 0 | 0 | 0 | 0 | 0 | 0 |
| 127.9674 | 35.82441 | 0 | Broadleaf deciduous forest | 1 | 0 | 0 | 0 | 0 | 0 | 0 |
| 128.908 | 35.86742 | 0 | Broadleaf deciduous forest | 1 | 0 | 0 | 0 | 0 | 0 | 0 |
| 127.005 | 36.27188 | 0 | Broadleaf deciduous forest | 1 | 0 | 0 | 0 | 0 | 0 | 0 |
| 128.0152 | 36.28471 | 0 | Broadleaf deciduous forest | 1 | 0 | 0 | 0 | 0 | 0 | 0 |
| 128.1457 | 36.31013 | 0 | Broadleaf deciduous forest | 1 | 0 | 0 | 0 | 0 | 0 | 0 |
| 128.357 | 36.35872 | 0 | Broadleaf deciduous forest | 1 | 0 | 0 | 0 | 0 | 0 | 0 |
| 128.0177 | 36.44755 | 0 | Broadleaf deciduous forest | 1 | 0 | 0 | 0 | 0 | 0 | 0 |
| 128.2566 | 36.49282 | 0 | Broadleaf deciduous forest | 1 | 0 | 0 | 0 | 0 | 0 | 0 |
| 128.2387 | 36.50491 | 0 | Broadleaf deciduous forest | 1 | 0 | 0 | 0 | 0 | 0 | 0 |
| 128.0907 | 36.53069 | 0 | Broadleaf deciduous forest | 1 | 0 | 0 | 0 | 0 | 0 | 0 |
| 127.806 | 36.59452 | 0 | Broadleaf deciduous forest | 1 | 0 | 0 | 0 | 0 | 0 | 0 |
| 127.1136 | 36.66611 | 0 | Broadleaf deciduous forest | 1 | 0 | 0 | 0 | 0 | 0 | 0 |
| 127.5962 | 36.70207 | 0 | Broadleaf deciduous forest | 1 | 0 | 0 | 0 | 0 | 0 | 0 |
| 126.6203 | 36.71695 | 0 | Broadleaf deciduous forest | 1 | 0 | 0 | 0 | 0 | 0 | 0 |
| 126.6264 | 36.77089 | 0 | Broadleaf deciduous forest | 1 | 0 | 0 | 0 | 0 | 0 | 0 |
| 127.8947 | 36.80252 | 0 | Broadleaf deciduous forest | 1 | 0 | 0 | 0 | 0 | 0 | 0 |
| 127.496 | 36.81213 | 0 | Broadleaf deciduous forest | 1 | 0 | 0 | 0 | 0 | 0 | 0 |
| 127.4535 | 36.81699 | 0 | Broadleaf deciduous forest | 1 | 0 | 0 | 0 | 0 | 0 | 0 |
| 127.9301 | 36.82812 | 0 | Broadleaf deciduous forest | 1 | 0 | 0 | 0 | 0 | 0 | 0 |
| 127.3127 | 36.85736 | 0 | Broadleaf deciduous forest | 1 | 0 | 0 | 0 | 0 | 0 | 0 |
| 128.9761 | 36.8678 | 0 | Broadleaf deciduous forest | 1 | 0 | 0 | 0 | 0 | 0 | 0 |
| 127.2943 | 36.89878 | 0 | Broadleaf deciduous forest | 1 | 0 | 0 | 0 | 0 | 0 | 0 |
| 128.2436 | 36.90596 | 0 | Broadleaf deciduous forest | 1 | 0 | 0 | 0 | 0 | 0 | 0 |
| 128.226 | 36.91023 | 0 | Broadleaf deciduous forest | 1 | 0 | 0 | 0 | 0 | 0 | 0 |
| 127.8814 | 36.91719 | 0 | Broadleaf deciduous forest | 1 | 0 | 0 | 0 | 0 | 0 | 0 |
| 127.7303 | 36.92664 | 0 | Broadleaf deciduous forest | 1 | 0 | 0 | 0 | 0 | 0 | 0 |
| 128.1375 | 36.9288 | 0 | Broadleaf deciduous forest | 1 | 0 | 0 | 0 | 0 | 0 | 0 |
| 127.6279 | 37.01501 | 0 | Broadleaf deciduous forest | 1 | 0 | 0 | 0 | 0 | 0 | 0 |
| 127.3809 | 37.02099 | 0 | Broadleaf deciduous forest | 1 | 0 | 0 | 0 | 0 | 0 | 0 |
| 127.6877 | 37.09052 | 0 | Broadleaf deciduous forest | 1 | 0 | 0 | 0 | 0 | 0 | 0 |
| 127.7509 | 37.11781 | 0 | Broadleaf deciduous forest | 1 | 0 | 0 | 0 | 0 | 0 | 0 |
| 128.3206 | 37.11894 | 0 | Broadleaf deciduous forest | 1 | 0 | 0 | 0 | 0 | 0 | 0 |
| 128.7211 | 37.13583 | 0 | Broadleaf deciduous forest | 1 | 0 | 0 | 0 | 0 | 0 | 0 |
| 127.727 | 37.14423 | 0 | Broadleaf deciduous forest | 1 | 0 | 0 | 0 | 0 | 0 | 0 |
| 127.1742 | 37.15511 | 0 | Broadleaf deciduous forest | 1 | 0 | 0 | 0 | 0 | 0 | 0 |
| 127.8187 | 37.20046 | 0 | Broadleaf deciduous forest | 1 | 0 | 0 | 0 | 0 | 0 | 0 |
| 128.6652 | 37.22688 | 0 | Broadleaf deciduous forest | 1 | 0 | 0 | 0 | 0 | 0 | 0 |
| 127.4062 | 37.23195 | 0 | Broadleaf deciduous forest | 1 | 0 | 0 | 0 | 0 | 0 | 0 |
| 128.4293 | 37.2412 | 0 | Broadleaf deciduous forest | 1 | 0 | 0 | 0 | 0 | 0 | 0 |
| 128.5965 | 37.24894 | 0 | Broadleaf deciduous forest | 1 | 0 | 0 | 0 | 0 | 0 | 0 |
| 128.6164 | 37.25863 | 0 | Broadleaf deciduous forest | 1 | 0 | 0 | 0 | 0 | 0 | 0 |
| 128.5689 | 37.26547 | 0 | Broadleaf deciduous forest | 1 | 0 | 0 | 0 | 0 | 0 | 0 |
| 128.4722 | 37.26631 | 0 | Broadleaf deciduous forest | 1 | 0 | 0 | 0 | 0 | 0 | 0 |
| 127.9448 | 37.26822 | 0 | Broadleaf deciduous forest | 1 | 0 | 0 | 0 | 0 | 0 | 0 |
| 128.6675 | 37.26888 | 0 | Broadleaf deciduous forest | 1 | 0 | 0 | 0 | 0 | 0 | 0 |
| 127.7943 | 37.3041 | 0 | Broadleaf deciduous forest | 1 | 0 | 0 | 0 | 0 | 0 | 0 |
| 128.4219 | 37.32489 | 0 | Broadleaf deciduous forest | 1 | 0 | 0 | 0 | 0 | 0 | 0 |
| 128.5351 | 37.33264 | 0 | Broadleaf deciduous forest | 1 | 0 | 0 | 0 | 0 | 0 | 0 |
| 128.6088 | 37.33494 | 0 | Broadleaf deciduous forest | 1 | 0 | 0 | 0 | 0 | 0 | 0 |
| 128.7661 | 37.34626 | 0 | Broadleaf deciduous forest | 1 | 0 | 0 | 0 | 0 | 0 | 0 |
| 128.529 | 37.35767 | 0 | Broadleaf deciduous forest | 1 | 0 | 0 | 0 | 0 | 0 | 0 |
| 128.2102 | 37.36501 | 0 | Broadleaf deciduous forest | 1 | 0 | 0 | 0 | 0 | 0 | 0 |
| 127.73 | 37.36757 | 0 | Broadleaf deciduous forest | 1 | 0 | 0 | 0 | 0 | 0 | 0 |
| 128.39 | 37.41608 | 0 | Broadleaf deciduous forest | 1 | 0 | 0 | 0 | 0 | 0 | 0 |
| 128.3878 | 37.42851 | 0 | Broadleaf deciduous forest | 1 | 0 | 0 | 0 | 0 | 0 | 0 |
| 128.3006 | 37.45045 | 0 | Broadleaf deciduous forest | 1 | 0 | 0 | 0 | 0 | 0 | 0 |
| 128.2947 | 37.53052 | 0 | Broadleaf deciduous forest | 1 | 0 | 0 | 0 | 0 | 0 | 0 |
| 128.2785 | 37.53284 | 0 | Broadleaf deciduous forest | 1 | 0 | 0 | 0 | 0 | 0 | 0 |
| 128.3842 | 37.5471 | 0 | Broadleaf deciduous forest | 1 | 0 | 0 | 0 | 0 | 0 | 0 |
| 128.8752 | 37.60738 | 0 | Broadleaf deciduous forest | 1 | 0 | 0 | 0 | 0 | 0 | 0 |
| 127.5932 | 37.64033 | 0 | Broadleaf deciduous forest | 1 | 0 | 0 | 0 | 0 | 0 | 0 |
| 126.9563 | 37.65354 | 0 | Broadleaf deciduous forest | 1 | 0 | 0 | 0 | 0 | 0 | 0 |
| 127.5031 | 37.68778 | 0 | Broadleaf deciduous forest | 1 | 0 | 0 | 0 | 0 | 0 | 0 |
| 126.7087 | 37.87789 | 0 | Broadleaf deciduous forest | 1 | 0 | 0 | 0 | 0 | 0 | 0 |
| 127.5596 | 37.90486 | 0 | Broadleaf deciduous forest | 1 | 0 | 0 | 0 | 0 | 0 | 0 |
| 124.6422 | 37.93359 | 0 | Broadleaf deciduous forest | 1 | 0 | 0 | 0 | 0 | 0 | 0 |
| 127.0425 | 37.93905 | 0 | Broadleaf deciduous forest | 1 | 0 | 0 | 0 | 0 | 0 | 0 |
| 127.5953 | 37.97528 | 0 | Broadleaf deciduous forest | 1 | 0 | 0 | 0 | 0 | 0 | 0 |
| 127.6019 | 37.9775 | 0 | Broadleaf deciduous forest | 1 | 0 | 0 | 0 | 0 | 0 | 0 |
| 127.6053 | 37.98925 | 0 | Broadleaf deciduous forest | 1 | 0 | 0 | 0 | 0 | 0 | 0 |
| 127.0509 | 37.99593 | 0 | Broadleaf deciduous forest | 1 | 0 | 0 | 0 | 0 | 0 | 0 |
| 127.5816 | 37.99953 | 0 | Broadleaf deciduous forest | 1 | 0 | 0 | 0 | 0 | 0 | 0 |
| 127.0215 | 38.00736 | 0 | Broadleaf deciduous forest | 1 | 0 | 0 | 0 | 0 | 0 | 0 |
| 127.18 | 38.03396 | 0 | Broadleaf deciduous forest | 1 | 0 | 0 | 0 | 0 | 0 | 0 |
| 127.1026 | 38.1258 | 0 | Broadleaf deciduous forest | 1 | 0 | 0 | 0 | 0 | 0 | 0 |
| 127.1293 | 38.13345 | 0 | Broadleaf deciduous forest | 1 | 0 | 0 | 0 | 0 | 0 | 0 |
| 127.1784 | 38.14355 | 0 | Broadleaf deciduous forest | 1 | 0 | 0 | 0 | 0 | 0 | 0 |
| 127.1456 | 38.17082 | 0 | Broadleaf deciduous forest | 1 | 0 | 0 | 0 | 0 | 0 | 0 |
| 127.4725 | 38.20472 | 0 | Broadleaf deciduous forest | 1 | 0 | 0 | 0 | 0 | 0 | 0 |
| 127.5767 | 38.31384 | 0 | Broadleaf deciduous forest | 1 | 0 | 0 | 0 | 0 | 0 | 0 |
| 128.3407 | 38.41266 | 0 | Broadleaf deciduous forest | 1 | 0 | 0 | 0 | 0 | 0 | 0 |
| 128.3872 | 38.42944 | 0 | Broadleaf deciduous forest | 1 | 0 | 0 | 0 | 0 | 0 | 0 |
| 128.2687 | 38.49063 | 0 | Broadleaf deciduous forest | 1 | 0 | 0 | 0 | 0 | 0 | 0 |
| 126.3864 | 41.97143 | 0 | Broadleaf deciduous forest | 1 | 0 | 0 | 0 | 0 | 0 | 0 |
| 130.2487 | 42.23715 | 0 | Broadleaf deciduous forest | 1 | 0 | 0 | 0 | 0 | 0 | 0 |
| 130.2245 | 42.25347 | 0 | Broadleaf deciduous forest | 1 | 0 | 0 | 0 | 0 | 0 | 0 |
| 130.2435 | 42.28709 | 0 | Broadleaf deciduous forest | 1 | 0 | 0 | 0 | 0 | 0 | 0 |
| 130.2078 | 42.29204 | 0 | Broadleaf deciduous forest | 1 | 0 | 0 | 0 | 0 | 0 | 0 |
| 130.2211 | 42.29834 | 0 | Broadleaf deciduous forest | 1 | 0 | 0 | 0 | 0 | 0 | 0 |
| 130.3708 | 42.29899 | 0 | Broadleaf deciduous forest | 1 | 0 | 0 | 0 | 0 | 0 | 0 |
| 130.3133 | 42.30049 | 0 | Broadleaf deciduous forest | 1 | 0 | 0 | 0 | 0 | 0 | 0 |
| 130.228 | 42.31233 | 0 | Broadleaf deciduous forest | 1 | 0 | 0 | 0 | 0 | 0 | 0 |
| 130.218 | 42.33008 | 0 | Broadleaf deciduous forest | 1 | 0 | 0 | 0 | 0 | 0 | 0 |
| 130.2429 | 42.33925 | 0 | Broadleaf deciduous forest | 1 | 0 | 0 | 0 | 0 | 0 | 0 |
| 130.4325 | 42.37006 | 0 | Broadleaf deciduous forest | 1 | 0 | 0 | 0 | 0 | 0 | 0 |
| 130.4124 | 42.379 | 0 | Broadleaf deciduous forest | 1 | 0 | 0 | 0 | 0 | 0 | 0 |
| 130.2571 | 42.38224 | 0 | Broadleaf deciduous forest | 1 | 0 | 0 | 0 | 0 | 0 | 0 |
| 130.4548 | 42.38254 | 0 | Broadleaf deciduous forest | 1 | 0 | 0 | 0 | 0 | 0 | 0 |
| 130.2406 | 42.3845 | 0 | Broadleaf deciduous forest | 1 | 0 | 0 | 0 | 0 | 0 | 0 |
| 130.498 | 42.39299 | 0 | Broadleaf deciduous forest | 1 | 0 | 0 | 0 | 0 | 0 | 0 |
| 130.291 | 42.39624 | 0 | Broadleaf deciduous forest | 1 | 0 | 0 | 0 | 0 | 0 | 0 |
| 128.1264 | 42.47998 | 0 | Broadleaf deciduous forest | 1 | 0 | 0 | 0 | 0 | 0 | 0 |
| 127.4917 | 35.24778 | 0 | Broadleaf evergreen forest | 1 | 0 | 0 | 0 | 0 | 0 | 0 |
| 128.1641 | 35.70705 | 0 | Broadleaf evergreen forest | 1 | 0 | 0 | 0 | 0 | 0 | 0 |
| 128.1843 | 35.75487 | 0 | Broadleaf evergreen forest | 1 | 0 | 0 | 0 | 0 | 0 | 0 |
| 127.9974 | 35.7747 | 0 | Broadleaf evergreen forest | 1 | 0 | 0 | 0 | 0 | 0 | 0 |
| 128.0379 | 36.40297 | 0 | Broadleaf evergreen forest | 1 | 0 | 0 | 0 | 0 | 0 | 0 |
| 127.2831 | 36.87276 | 0 | Broadleaf evergreen forest | 1 | 0 | 0 | 0 | 0 | 0 | 0 |
| 128.6448 | 36.8912 | 0 | Broadleaf evergreen forest | 1 | 0 | 0 | 0 | 0 | 0 | 0 |
| 128.9434 | 36.91627 | 0 | Broadleaf evergreen forest | 1 | 0 | 0 | 0 | 0 | 0 | 0 |
| 127.1108 | 37.13967 | 0 | Broadleaf evergreen forest | 1 | 0 | 0 | 0 | 0 | 0 | 0 |
| 126.7998 | 37.9847 | 0 | Broadleaf evergreen forest | 1 | 0 | 0 | 0 | 0 | 0 | 0 |
| 127.4653 | 38.19139 | 0 | Broadleaf evergreen forest | 1 | 0 | 0 | 0 | 0 | 0 | 0 |
| 125.2726 | 38.84808 | 0 | Broadleaf evergreen forest | 1 | 0 | 0 | 0 | 0 | 0 | 0 |
| 130.257 | 42.28976 | 0 | Broadleaf evergreen forest | 1 | 0 | 0 | 0 | 0 | 0 | 0 |
| 130.5219 | 42.39029 | 0 | Broadleaf evergreen forest | 1 | 0 | 0 | 0 | 0 | 0 | 0 |
| 126.8956 | 35.84019 | 1 | Cropland | 0 | 1 | 0 | 0 | 0 | 0 | 0 |
| 126.8003 | 35.88137 | 1 | Cropland | 0 | 1 | 0 | 0 | 0 | 0 | 0 |
| 127.0673 | 35.96983 | 1 | Cropland | 0 | 1 | 0 | 0 | 0 | 0 | 0 |
| 127.0138 | 36.26531 | 1 | Cropland | 0 | 1 | 0 | 0 | 0 | 0 | 0 |
| 127.1837 | 36.31141 | 1 | Cropland | 0 | 1 | 0 | 0 | 0 | 0 | 0 |
| 126.806 | 36.90589 | 1 | Cropland | 0 | 1 | 0 | 0 | 0 | 0 | 0 |
| 126.9791 | 36.97719 | 1 | Cropland | 0 | 1 | 0 | 0 | 0 | 0 | 0 |
| 126.5996 | 37.60648 | 1 | Cropland | 0 | 1 | 0 | 0 | 0 | 0 | 0 |
| 126.6103 | 37.7891 | 1 | Cropland | 0 | 1 | 0 | 0 | 0 | 0 | 0 |
| 126.7676 | 37.7959 | 1 | Cropland | 0 | 1 | 0 | 0 | 0 | 0 | 0 |
| 127.0847 | 38.06315 | 1 | Cropland | 0 | 1 | 0 | 0 | 0 | 0 | 0 |
| 125.2948 | 38.83659 | 1 | Cropland | 0 | 1 | 0 | 0 | 0 | 0 | 0 |
| 125.2195 | 38.92011 | 1 | Cropland | 0 | 1 | 0 | 0 | 0 | 0 | 0 |
| 125.6883 | 39.06454 | 1 | Cropland | 0 | 1 | 0 | 0 | 0 | 0 | 0 |
| 124.861 | 39.743 | 1 | Cropland | 0 | 1 | 0 | 0 | 0 | 0 | 0 |
| 124.923 | 39.7094 | 1 | Cropland | 0 | 1 | 0 | 0 | 0 | 0 | 0 |
| 125.2406 | 38.86312 | 1 | Cropland | 0 | 1 | 0 | 0 | 0 | 0 | 0 |
| 125.2692 | 38.96191 | 1 | Cropland | 0 | 1 | 0 | 0 | 0 | 0 | 0 |
| 125.2788 | 38.8651 | 1 | Cropland | 0 | 1 | 0 | 0 | 0 | 0 | 0 |
| 125.3767 | 39.47147 | 1 | Cropland | 0 | 1 | 0 | 0 | 0 | 0 | 0 |
| 125.4323 | 39.49264 | 1 | Cropland | 0 | 1 | 0 | 0 | 0 | 0 | 0 |
| 125.45 | 39.5452 | 1 | Cropland | 0 | 1 | 0 | 0 | 0 | 0 | 0 |
| 125.5637 | 39.49301 | 1 | Cropland | 0 | 1 | 0 | 0 | 0 | 0 | 0 |
| 125.5958 | 39.20426 | 1 | Cropland | 0 | 1 | 0 | 0 | 0 | 0 | 0 |
| 125.615 | 38.4454 | 1 | Cropland | 0 | 1 | 0 | 0 | 0 | 0 | 0 |
| 125.7457 | 39.11752 | 1 | Cropland | 0 | 1 | 0 | 0 | 0 | 0 | 0 |
| 126.4141 | 36.63627 | 1 | Cropland | 0 | 1 | 0 | 0 | 0 | 0 | 0 |
| 126.4765 | 36.76844 | 1 | Cropland | 0 | 1 | 0 | 0 | 0 | 0 | 0 |
| 126.5131 | 36.63135 | 1 | Cropland | 0 | 1 | 0 | 0 | 0 | 0 | 0 |
| 126.5252 | 36.76177 | 1 | Cropland | 0 | 1 | 0 | 0 | 0 | 0 | 0 |
| 126.5289 | 36.65143 | 1 | Cropland | 0 | 1 | 0 | 0 | 0 | 0 | 0 |
| 126.5452 | 36.67784 | 1 | Cropland | 0 | 1 | 0 | 0 | 0 | 0 | 0 |
| 126.5791 | 36.3333 | 1 | Cropland | 0 | 1 | 0 | 0 | 0 | 0 | 0 |
| 126.5813 | 36.2498 | 1 | Cropland | 0 | 1 | 0 | 0 | 0 | 0 | 0 |
| 126.612 | 36.9118 | 1 | Cropland | 0 | 1 | 0 | 0 | 0 | 0 | 0 |
| 126.6451 | 37.6221 | 1 | Cropland | 0 | 1 | 0 | 0 | 0 | 0 | 0 |
| 126.6656 | 37.66279 | 1 | Cropland | 0 | 1 | 0 | 0 | 0 | 0 | 0 |
| 126.672 | 37.9534 | 1 | Cropland | 0 | 1 | 0 | 0 | 0 | 0 | 0 |
| 126.6882 | 36.62565 | 1 | Cropland | 0 | 1 | 0 | 0 | 0 | 0 | 0 |
| 126.7325 | 35.62163 | 1 | Cropland | 0 | 1 | 0 | 0 | 0 | 0 | 0 |
| 126.7908 | 37.54887 | 1 | Cropland | 0 | 1 | 0 | 0 | 0 | 0 | 0 |
| 126.8438 | 35.91251 | 1 | Cropland | 0 | 1 | 0 | 0 | 0 | 0 | 0 |
| 126.8684 | 36.8155 | 1 | Cropland | 0 | 1 | 0 | 0 | 0 | 0 | 0 |
| 126.8698 | 37.05841 | 1 | Cropland | 0 | 1 | 0 | 0 | 0 | 0 | 0 |
| 126.8711 | 36.97192 | 1 | Cropland | 0 | 1 | 0 | 0 | 0 | 0 | 0 |
| 126.878 | 36.9954 | 1 | Cropland | 0 | 1 | 0 | 0 | 0 | 0 | 0 |
| 126.8786 | 36.83721 | 1 | Cropland | 0 | 1 | 0 | 0 | 0 | 0 | 0 |
| 126.8956 | 36.95393 | 1 | Cropland | 0 | 1 | 0 | 0 | 0 | 0 | 0 |
| 126.919 | 36.31757 | 1 | Cropland | 0 | 1 | 0 | 0 | 0 | 0 | 0 |
| 126.9243 | 36.94103 | 1 | Cropland | 0 | 1 | 0 | 0 | 0 | 0 | 0 |
| 126.9318 | 36.31882 | 1 | Cropland | 0 | 1 | 0 | 0 | 0 | 0 | 0 |
| 126.938 | 36.8049 | 1 | Cropland | 0 | 1 | 0 | 0 | 0 | 0 | 0 |
| 126.9457 | 36.98268 | 1 | Cropland | 0 | 1 | 0 | 0 | 0 | 0 | 0 |
| 126.9672 | 36.80647 | 1 | Cropland | 0 | 1 | 0 | 0 | 0 | 0 | 0 |
| 126.9679 | 35.85279 | 1 | Cropland | 0 | 1 | 0 | 0 | 0 | 0 | 0 |
| 127.0013 | 35.89609 | 1 | Cropland | 0 | 1 | 0 | 0 | 0 | 0 | 0 |
| 127.053 | 36.0864 | 1 | Cropland | 0 | 1 | 0 | 0 | 0 | 0 | 0 |
| 127.056 | 36.0954 | 1 | Cropland | 0 | 1 | 0 | 0 | 0 | 0 | 0 |
| 127.0629 | 35.97767 | 1 | Cropland | 0 | 1 | 0 | 0 | 0 | 0 | 0 |
| 127.1349 | 36.23129 | 1 | Cropland | 0 | 1 | 0 | 0 | 0 | 0 | 0 |
| 127.1554 | 35.94961 | 1 | Cropland | 0 | 1 | 0 | 0 | 0 | 0 | 0 |
| 127.2128 | 37.14774 | 1 | Cropland | 0 | 1 | 0 | 0 | 0 | 0 | 0 |
| 127.2305 | 36.49944 | 1 | Cropland | 0 | 1 | 0 | 0 | 0 | 0 | 0 |
| 127.2664 | 36.54985 | 1 | Cropland | 0 | 1 | 0 | 0 | 0 | 0 | 0 |
| 127.3421 | 36.58559 | 1 | Cropland | 0 | 1 | 0 | 0 | 0 | 0 | 0 |
| 127.3497 | 36.59865 | 1 | Cropland | 0 | 1 | 0 | 0 | 0 | 0 | 0 |
| 127.6262 | 37.02333 | 1 | Cropland | 0 | 1 | 0 | 0 | 0 | 0 | 0 |
| 128.8936 | 35.19444 | 0 | Cropland | 0 | 1 | 0 | 0 | 0 | 0 | 0 |
| 127.9969 | 35.66611 | 0 | Cropland | 0 | 1 | 0 | 0 | 0 | 0 | 0 |
| 126.8841 | 35.85464 | 0 | Cropland | 0 | 1 | 0 | 0 | 0 | 0 | 0 |
| 126.8658 | 35.88536 | 0 | Cropland | 0 | 1 | 0 | 0 | 0 | 0 | 0 |
| 127.0083 | 35.88611 | 0 | Cropland | 0 | 1 | 0 | 0 | 0 | 0 | 0 |
| 127.0156 | 35.88778 | 0 | Cropland | 0 | 1 | 0 | 0 | 0 | 0 | 0 |
| 127.0392 | 35.91028 | 0 | Cropland | 0 | 1 | 0 | 0 | 0 | 0 | 0 |
| 127.0622 | 35.91745 | 0 | Cropland | 0 | 1 | 0 | 0 | 0 | 0 | 0 |
| 126.8132 | 35.92876 | 0 | Cropland | 0 | 1 | 0 | 0 | 0 | 0 | 0 |
| 127.0047 | 35.93111 | 0 | Cropland | 0 | 1 | 0 | 0 | 0 | 0 | 0 |
| 126.9056 | 35.96083 | 0 | Cropland | 0 | 1 | 0 | 0 | 0 | 0 | 0 |
| 126.9173 | 35.97219 | 0 | Cropland | 0 | 1 | 0 | 0 | 0 | 0 | 0 |
| 126.9458 | 35.99417 | 0 | Cropland | 0 | 1 | 0 | 0 | 0 | 0 | 0 |
| 126.9556 | 35.99692 | 0 | Cropland | 0 | 1 | 0 | 0 | 0 | 0 | 0 |
| 126.7425 | 36.05472 | 0 | Cropland | 0 | 1 | 0 | 0 | 0 | 0 | 0 |
| 128.3682 | 36.15487 | 0 | Cropland | 0 | 1 | 0 | 0 | 0 | 0 | 0 |
| 128.227 | 36.16778 | 0 | Cropland | 0 | 1 | 0 | 0 | 0 | 0 | 0 |
| 128.2278 | 36.17973 | 0 | Cropland | 0 | 1 | 0 | 0 | 0 | 0 | 0 |
| 128.2306 | 36.19466 | 0 | Cropland | 0 | 1 | 0 | 0 | 0 | 0 | 0 |
| 128.2141 | 36.20115 | 0 | Cropland | 0 | 1 | 0 | 0 | 0 | 0 | 0 |
| 128.3337 | 36.21746 | 0 | Cropland | 0 | 1 | 0 | 0 | 0 | 0 | 0 |
| 128.2377 | 36.24744 | 0 | Cropland | 0 | 1 | 0 | 0 | 0 | 0 | 0 |
| 128.2082 | 36.2581 | 0 | Cropland | 0 | 1 | 0 | 0 | 0 | 0 | 0 |
| 126.8422 | 36.25877 | 0 | Cropland | 0 | 1 | 0 | 0 | 0 | 0 | 0 |
| 126.8782 | 36.28371 | 0 | Cropland | 0 | 1 | 0 | 0 | 0 | 0 | 0 |
| 128.0843 | 36.28864 | 0 | Cropland | 0 | 1 | 0 | 0 | 0 | 0 | 0 |
| 128.4632 | 36.30154 | 0 | Cropland | 0 | 1 | 0 | 0 | 0 | 0 | 0 |
| 126.9497 | 36.33667 | 0 | Cropland | 0 | 1 | 0 | 0 | 0 | 0 | 0 |
| 126.941 | 36.36845 | 0 | Cropland | 0 | 1 | 0 | 0 | 0 | 0 | 0 |
| 128.4384 | 36.37533 | 0 | Cropland | 0 | 1 | 0 | 0 | 0 | 0 | 0 |
| 128.4211 | 36.37597 | 0 | Cropland | 0 | 1 | 0 | 0 | 0 | 0 | 0 |
| 128.3934 | 36.38911 | 0 | Cropland | 0 | 1 | 0 | 0 | 0 | 0 | 0 |
| 128.4004 | 36.39598 | 0 | Cropland | 0 | 1 | 0 | 0 | 0 | 0 | 0 |
| 128.1655 | 36.3981 | 0 | Cropland | 0 | 1 | 0 | 0 | 0 | 0 | 0 |
| 128.3943 | 36.40319 | 0 | Cropland | 0 | 1 | 0 | 0 | 0 | 0 | 0 |
| 128.3647 | 36.40407 | 0 | Cropland | 0 | 1 | 0 | 0 | 0 | 0 | 0 |
| 128.3804 | 36.40686 | 0 | Cropland | 0 | 1 | 0 | 0 | 0 | 0 | 0 |
| 128.1839 | 36.43121 | 0 | Cropland | 0 | 1 | 0 | 0 | 0 | 0 | 0 |
| 128.3587 | 36.44283 | 0 | Cropland | 0 | 1 | 0 | 0 | 0 | 0 | 0 |
| 128.2423 | 36.47938 | 0 | Cropland | 0 | 1 | 0 | 0 | 0 | 0 | 0 |
| 128.2996 | 36.48534 | 0 | Cropland | 0 | 1 | 0 | 0 | 0 | 0 | 0 |
| 128.2872 | 36.49897 | 0 | Cropland | 0 | 1 | 0 | 0 | 0 | 0 | 0 |
| 128.1322 | 36.50694 | 0 | Cropland | 0 | 1 | 0 | 0 | 0 | 0 | 0 |
| 128.2079 | 36.58448 | 0 | Cropland | 0 | 1 | 0 | 0 | 0 | 0 | 0 |
| 128.2867 | 36.59547 | 0 | Cropland | 0 | 1 | 0 | 0 | 0 | 0 | 0 |
| 128.2352 | 36.59986 | 0 | Cropland | 0 | 1 | 0 | 0 | 0 | 0 | 0 |
| 126.7358 | 36.68611 | 0 | Cropland | 0 | 1 | 0 | 0 | 0 | 0 | 0 |
| 126.6878 | 36.69971 | 0 | Cropland | 0 | 1 | 0 | 0 | 0 | 0 | 0 |
| 126.7332 | 36.70024 | 0 | Cropland | 0 | 1 | 0 | 0 | 0 | 0 | 0 |
| 127.4647 | 36.71637 | 0 | Cropland | 0 | 1 | 0 | 0 | 0 | 0 | 0 |
| 127.1303 | 36.71861 | 0 | Cropland | 0 | 1 | 0 | 0 | 0 | 0 | 0 |
| 127.4637 | 36.72138 | 0 | Cropland | 0 | 1 | 0 | 0 | 0 | 0 | 0 |
| 127.4738 | 36.72694 | 0 | Cropland | 0 | 1 | 0 | 0 | 0 | 0 | 0 |
| 127.4869 | 36.73199 | 0 | Cropland | 0 | 1 | 0 | 0 | 0 | 0 | 0 |
| 127.0081 | 36.80833 | 0 | Cropland | 0 | 1 | 0 | 0 | 0 | 0 | 0 |
| 126.9869 | 36.81076 | 0 | Cropland | 0 | 1 | 0 | 0 | 0 | 0 | 0 |
| 126.8907 | 36.81174 | 0 | Cropland | 0 | 1 | 0 | 0 | 0 | 0 | 0 |
| 127.4587 | 36.8514 | 0 | Cropland | 0 | 1 | 0 | 0 | 0 | 0 | 0 |
| 127.4256 | 36.86342 | 0 | Cropland | 0 | 1 | 0 | 0 | 0 | 0 | 0 |
| 127.5031 | 36.86628 | 0 | Cropland | 0 | 1 | 0 | 0 | 0 | 0 | 0 |
| 127.4954 | 36.86861 | 0 | Cropland | 0 | 1 | 0 | 0 | 0 | 0 | 0 |
| 127.8617 | 36.87526 | 0 | Cropland | 0 | 1 | 0 | 0 | 0 | 0 | 0 |
| 127.854 | 36.87987 | 0 | Cropland | 0 | 1 | 0 | 0 | 0 | 0 | 0 |
| 127.472 | 36.88051 | 0 | Cropland | 0 | 1 | 0 | 0 | 0 | 0 | 0 |
| 127.4562 | 36.89671 | 0 | Cropland | 0 | 1 | 0 | 0 | 0 | 0 | 0 |
| 127.2047 | 36.90225 | 0 | Cropland | 0 | 1 | 0 | 0 | 0 | 0 | 0 |
| 127.7758 | 36.90503 | 0 | Cropland | 0 | 1 | 0 | 0 | 0 | 0 | 0 |
| 127.7665 | 36.90801 | 0 | Cropland | 0 | 1 | 0 | 0 | 0 | 0 | 0 |
| 127.7722 | 36.90878 | 0 | Cropland | 0 | 1 | 0 | 0 | 0 | 0 | 0 |
| 127.6951 | 36.91126 | 0 | Cropland | 0 | 1 | 0 | 0 | 0 | 0 | 0 |
| 127.7779 | 36.91146 | 0 | Cropland | 0 | 1 | 0 | 0 | 0 | 0 | 0 |
| 127.7615 | 36.91356 | 0 | Cropland | 0 | 1 | 0 | 0 | 0 | 0 | 0 |
| 127.5376 | 36.91746 | 0 | Cropland | 0 | 1 | 0 | 0 | 0 | 0 | 0 |
| 127.7665 | 36.91796 | 0 | Cropland | 0 | 1 | 0 | 0 | 0 | 0 | 0 |
| 127.7577 | 36.91848 | 0 | Cropland | 0 | 1 | 0 | 0 | 0 | 0 | 0 |
| 127.6995 | 36.92086 | 0 | Cropland | 0 | 1 | 0 | 0 | 0 | 0 | 0 |
| 126.7936 | 36.92694 | 0 | Cropland | 0 | 1 | 0 | 0 | 0 | 0 | 0 |
| 127.0751 | 36.92849 | 0 | Cropland | 0 | 1 | 0 | 0 | 0 | 0 | 0 |
| 126.9947 | 36.92916 | 0 | Cropland | 0 | 1 | 0 | 0 | 0 | 0 | 0 |
| 127.7639 | 36.93057 | 0 | Cropland | 0 | 1 | 0 | 0 | 0 | 0 | 0 |
| 128.1767 | 36.93528 | 0 | Cropland | 0 | 1 | 0 | 0 | 0 | 0 | 0 |
| 127.4363 | 36.94121 | 0 | Cropland | 0 | 1 | 0 | 0 | 0 | 0 | 0 |
| 127.4593 | 36.9546 | 0 | Cropland | 0 | 1 | 0 | 0 | 0 | 0 | 0 |
| 127.4739 | 36.95618 | 0 | Cropland | 0 | 1 | 0 | 0 | 0 | 0 | 0 |
| 127.7867 | 36.95992 | 0 | Cropland | 0 | 1 | 0 | 0 | 0 | 0 | 0 |
| 127.8372 | 36.96282 | 0 | Cropland | 0 | 1 | 0 | 0 | 0 | 0 | 0 |
| 127.1014 | 36.96306 | 0 | Cropland | 0 | 1 | 0 | 0 | 0 | 0 | 0 |
| 126.9576 | 36.96314 | 0 | Cropland | 0 | 1 | 0 | 0 | 0 | 0 | 0 |
| 127.7893 | 36.96484 | 0 | Cropland | 0 | 1 | 0 | 0 | 0 | 0 | 0 |
| 127.1911 | 36.96917 | 0 | Cropland | 0 | 1 | 0 | 0 | 0 | 0 | 0 |
| 127.0841 | 36.97008 | 0 | Cropland | 0 | 1 | 0 | 0 | 0 | 0 | 0 |
| 127.7686 | 36.97066 | 0 | Cropland | 0 | 1 | 0 | 0 | 0 | 0 | 0 |
| 127.7742 | 36.97089 | 0 | Cropland | 0 | 1 | 0 | 0 | 0 | 0 | 0 |
| 127.4947 | 36.9744 | 0 | Cropland | 0 | 1 | 0 | 0 | 0 | 0 | 0 |
| 127.7879 | 36.97463 | 0 | Cropland | 0 | 1 | 0 | 0 | 0 | 0 | 0 |
| 127.7819 | 36.97489 | 0 | Cropland | 0 | 1 | 0 | 0 | 0 | 0 | 0 |
| 127.5012 | 36.98273 | 0 | Cropland | 0 | 1 | 0 | 0 | 0 | 0 | 0 |
| 126.985 | 36.984 | 0 | Cropland | 0 | 1 | 0 | 0 | 0 | 0 | 0 |
| 127.482 | 36.98869 | 0 | Cropland | 0 | 1 | 0 | 0 | 0 | 0 | 0 |
| 127.4763 | 36.98886 | 0 | Cropland | 0 | 1 | 0 | 0 | 0 | 0 | 0 |
| 127.0097 | 36.9925 | 0 | Cropland | 0 | 1 | 0 | 0 | 0 | 0 | 0 |
| 127.0217 | 36.99358 | 0 | Cropland | 0 | 1 | 0 | 0 | 0 | 0 | 0 |
| 127.7626 | 36.99524 | 0 | Cropland | 0 | 1 | 0 | 0 | 0 | 0 | 0 |
| 127.4823 | 36.99884 | 0 | Cropland | 0 | 1 | 0 | 0 | 0 | 0 | 0 |
| 126.9834 | 37.00042 | 0 | Cropland | 0 | 1 | 0 | 0 | 0 | 0 | 0 |
| 127.7215 | 37.00388 | 0 | Cropland | 0 | 1 | 0 | 0 | 0 | 0 | 0 |
| 127.3117 | 37.00778 | 0 | Cropland | 0 | 1 | 0 | 0 | 0 | 0 | 0 |
| 126.8631 | 37.00889 | 0 | Cropland | 0 | 1 | 0 | 0 | 0 | 0 | 0 |
| 127.4875 | 37.01158 | 0 | Cropland | 0 | 1 | 0 | 0 | 0 | 0 | 0 |
| 127.3071 | 37.01382 | 0 | Cropland | 0 | 1 | 0 | 0 | 0 | 0 | 0 |
| 126.8822 | 37.0219 | 0 | Cropland | 0 | 1 | 0 | 0 | 0 | 0 | 0 |
| 127.9266 | 37.02614 | 0 | Cropland | 0 | 1 | 0 | 0 | 0 | 0 | 0 |
| 127.5149 | 37.0274 | 0 | Cropland | 0 | 1 | 0 | 0 | 0 | 0 | 0 |
| 126.8845 | 37.03007 | 0 | Cropland | 0 | 1 | 0 | 0 | 0 | 0 | 0 |
| 127.5414 | 37.04214 | 0 | Cropland | 0 | 1 | 0 | 0 | 0 | 0 | 0 |
| 127.5266 | 37.05123 | 0 | Cropland | 0 | 1 | 0 | 0 | 0 | 0 | 0 |
| 127.541 | 37.05232 | 0 | Cropland | 0 | 1 | 0 | 0 | 0 | 0 | 0 |
| 127.9153 | 37.07624 | 0 | Cropland | 0 | 1 | 0 | 0 | 0 | 0 | 0 |
| 127.5204 | 37.07678 | 0 | Cropland | 0 | 1 | 0 | 0 | 0 | 0 | 0 |
| 127.5409 | 37.09659 | 0 | Cropland | 0 | 1 | 0 | 0 | 0 | 0 | 0 |
| 127.6349 | 37.09771 | 0 | Cropland | 0 | 1 | 0 | 0 | 0 | 0 | 0 |
| 127.567 | 37.10317 | 0 | Cropland | 0 | 1 | 0 | 0 | 0 | 0 | 0 |
| 127.4927 | 37.10404 | 0 | Cropland | 0 | 1 | 0 | 0 | 0 | 0 | 0 |
| 127.5449 | 37.1212 | 0 | Cropland | 0 | 1 | 0 | 0 | 0 | 0 | 0 |
| 127.625 | 37.1239 | 0 | Cropland | 0 | 1 | 0 | 0 | 0 | 0 | 0 |
| 127.5276 | 37.1308 | 0 | Cropland | 0 | 1 | 0 | 0 | 0 | 0 | 0 |
| 127.6175 | 37.14482 | 0 | Cropland | 0 | 1 | 0 | 0 | 0 | 0 | 0 |
| 127.6248 | 37.14576 | 0 | Cropland | 0 | 1 | 0 | 0 | 0 | 0 | 0 |
| 126.7057 | 37.15061 | 0 | Cropland | 0 | 1 | 0 | 0 | 0 | 0 | 0 |
| 127.4817 | 37.15913 | 0 | Cropland | 0 | 1 | 0 | 0 | 0 | 0 | 0 |
| 127.4896 | 37.15922 | 0 | Cropland | 0 | 1 | 0 | 0 | 0 | 0 | 0 |
| 128.2056 | 37.16273 | 0 | Cropland | 0 | 1 | 0 | 0 | 0 | 0 | 0 |
| 127.6589 | 37.16361 | 0 | Cropland | 0 | 1 | 0 | 0 | 0 | 0 | 0 |
| 127.4987 | 37.16734 | 0 | Cropland | 0 | 1 | 0 | 0 | 0 | 0 | 0 |
| 127.6561 | 37.17029 | 0 | Cropland | 0 | 1 | 0 | 0 | 0 | 0 | 0 |
| 127.6717 | 37.18037 | 0 | Cropland | 0 | 1 | 0 | 0 | 0 | 0 | 0 |
| 127.6698 | 37.19239 | 0 | Cropland | 0 | 1 | 0 | 0 | 0 | 0 | 0 |
| 127.4618 | 37.19916 | 0 | Cropland | 0 | 1 | 0 | 0 | 0 | 0 | 0 |
| 127.5457 | 37.21714 | 0 | Cropland | 0 | 1 | 0 | 0 | 0 | 0 | 0 |
| 127.5282 | 37.21763 | 0 | Cropland | 0 | 1 | 0 | 0 | 0 | 0 | 0 |
| 127.4428 | 37.2427 | 0 | Cropland | 0 | 1 | 0 | 0 | 0 | 0 | 0 |
| 127.5159 | 37.24711 | 0 | Cropland | 0 | 1 | 0 | 0 | 0 | 0 | 0 |
| 127.8028 | 37.27917 | 0 | Cropland | 0 | 1 | 0 | 0 | 0 | 0 | 0 |
| 127.8147 | 37.29128 | 0 | Cropland | 0 | 1 | 0 | 0 | 0 | 0 | 0 |
| 127.5662 | 37.29604 | 0 | Cropland | 0 | 1 | 0 | 0 | 0 | 0 | 0 |
| 127.8152 | 37.30241 | 0 | Cropland | 0 | 1 | 0 | 0 | 0 | 0 | 0 |
| 127.5619 | 37.30561 | 0 | Cropland | 0 | 1 | 0 | 0 | 0 | 0 | 0 |
| 127.8153 | 37.30859 | 0 | Cropland | 0 | 1 | 0 | 0 | 0 | 0 | 0 |
| 127.5651 | 37.31015 | 0 | Cropland | 0 | 1 | 0 | 0 | 0 | 0 | 0 |
| 127.5678 | 37.31464 | 0 | Cropland | 0 | 1 | 0 | 0 | 0 | 0 | 0 |
| 127.6166 | 37.31721 | 0 | Cropland | 0 | 1 | 0 | 0 | 0 | 0 | 0 |
| 127.5456 | 37.33696 | 0 | Cropland | 0 | 1 | 0 | 0 | 0 | 0 | 0 |
| 127.5606 | 37.33972 | 0 | Cropland | 0 | 1 | 0 | 0 | 0 | 0 | 0 |
| 127.6858 | 37.37512 | 0 | Cropland | 0 | 1 | 0 | 0 | 0 | 0 | 0 |
| 126.8036 | 37.41778 | 0 | Cropland | 0 | 1 | 0 | 0 | 0 | 0 | 0 |
| 126.408 | 37.45059 | 0 | Cropland | 0 | 1 | 0 | 0 | 0 | 0 | 0 |
| 126.3683 | 37.46021 | 0 | Cropland | 0 | 1 | 0 | 0 | 0 | 0 | 0 |
| 126.7955 | 37.57068 | 0 | Cropland | 0 | 1 | 0 | 0 | 0 | 0 | 0 |
| 126.79 | 37.57171 | 0 | Cropland | 0 | 1 | 0 | 0 | 0 | 0 | 0 |
| 126.7864 | 37.57639 | 0 | Cropland | 0 | 1 | 0 | 0 | 0 | 0 | 0 |
| 126.6053 | 37.59618 | 0 | Cropland | 0 | 1 | 0 | 0 | 0 | 0 | 0 |
| 126.6112 | 37.60028 | 0 | Cropland | 0 | 1 | 0 | 0 | 0 | 0 | 0 |
| 126.5705 | 37.60403 | 0 | Cropland | 0 | 1 | 0 | 0 | 0 | 0 | 0 |
| 126.5639 | 37.60916 | 0 | Cropland | 0 | 1 | 0 | 0 | 0 | 0 | 0 |
| 126.5846 | 37.61121 | 0 | Cropland | 0 | 1 | 0 | 0 | 0 | 0 | 0 |
| 126.5661 | 37.6148 | 0 | Cropland | 0 | 1 | 0 | 0 | 0 | 0 | 0 |
| 126.5824 | 37.62534 | 0 | Cropland | 0 | 1 | 0 | 0 | 0 | 0 | 0 |
| 126.7264 | 37.6266 | 0 | Cropland | 0 | 1 | 0 | 0 | 0 | 0 | 0 |
| 126.5713 | 37.6365 | 0 | Cropland | 0 | 1 | 0 | 0 | 0 | 0 | 0 |
| 126.6407 | 37.65909 | 0 | Cropland | 0 | 1 | 0 | 0 | 0 | 0 | 0 |
| 126.6439 | 37.66353 | 0 | Cropland | 0 | 1 | 0 | 0 | 0 | 0 | 0 |
| 126.6297 | 37.66827 | 0 | Cropland | 0 | 1 | 0 | 0 | 0 | 0 | 0 |
| 126.6354 | 37.6695 | 0 | Cropland | 0 | 1 | 0 | 0 | 0 | 0 | 0 |
| 126.6248 | 37.67103 | 0 | Cropland | 0 | 1 | 0 | 0 | 0 | 0 | 0 |
| 126.6626 | 37.67269 | 0 | Cropland | 0 | 1 | 0 | 0 | 0 | 0 | 0 |
| 126.6523 | 37.67598 | 0 | Cropland | 0 | 1 | 0 | 0 | 0 | 0 | 0 |
| 126.6375 | 37.67778 | 0 | Cropland | 0 | 1 | 0 | 0 | 0 | 0 | 0 |
| 126.6483 | 37.68109 | 0 | Cropland | 0 | 1 | 0 | 0 | 0 | 0 | 0 |
| 126.6295 | 37.68444 | 0 | Cropland | 0 | 1 | 0 | 0 | 0 | 0 | 0 |
| 126.6223 | 37.68771 | 0 | Cropland | 0 | 1 | 0 | 0 | 0 | 0 | 0 |
| 126.6956 | 37.68825 | 0 | Cropland | 0 | 1 | 0 | 0 | 0 | 0 | 0 |
| 126.686 | 37.69722 | 0 | Cropland | 0 | 1 | 0 | 0 | 0 | 0 | 0 |
| 126.6144 | 37.69882 | 0 | Cropland | 0 | 1 | 0 | 0 | 0 | 0 | 0 |
| 126.5942 | 37.70514 | 0 | Cropland | 0 | 1 | 0 | 0 | 0 | 0 | 0 |
| 126.6073 | 37.71531 | 0 | Cropland | 0 | 1 | 0 | 0 | 0 | 0 | 0 |
| 126.7333 | 37.73835 | 0 | Cropland | 0 | 1 | 0 | 0 | 0 | 0 | 0 |
| 126.5814 | 37.73965 | 0 | Cropland | 0 | 1 | 0 | 0 | 0 | 0 | 0 |
| 126.7196 | 37.74029 | 0 | Cropland | 0 | 1 | 0 | 0 | 0 | 0 | 0 |
| 126.6936 | 37.74333 | 0 | Cropland | 0 | 1 | 0 | 0 | 0 | 0 | 0 |
| 126.5811 | 37.74639 | 0 | Cropland | 0 | 1 | 0 | 0 | 0 | 0 | 0 |
| 126.7081 | 37.74645 | 0 | Cropland | 0 | 1 | 0 | 0 | 0 | 0 | 0 |
| 126.5753 | 37.74784 | 0 | Cropland | 0 | 1 | 0 | 0 | 0 | 0 | 0 |
| 126.6104 | 37.74815 | 0 | Cropland | 0 | 1 | 0 | 0 | 0 | 0 | 0 |
| 126.6186 | 37.74917 | 0 | Cropland | 0 | 1 | 0 | 0 | 0 | 0 | 0 |
| 126.6946 | 37.75449 | 0 | Cropland | 0 | 1 | 0 | 0 | 0 | 0 | 0 |
| 126.7105 | 37.76356 | 0 | Cropland | 0 | 1 | 0 | 0 | 0 | 0 | 0 |
| 126.7744 | 37.77676 | 0 | Cropland | 0 | 1 | 0 | 0 | 0 | 0 | 0 |
| 126.2667 | 37.79373 | 0 | Cropland | 0 | 1 | 0 | 0 | 0 | 0 | 0 |
| 126.6128 | 37.79884 | 0 | Cropland | 0 | 1 | 0 | 0 | 0 | 0 | 0 |
| 126.7903 | 37.79945 | 0 | Cropland | 0 | 1 | 0 | 0 | 0 | 0 | 0 |
| 126.7946 | 37.80569 | 0 | Cropland | 0 | 1 | 0 | 0 | 0 | 0 | 0 |
| 126.7425 | 37.81275 | 0 | Cropland | 0 | 1 | 0 | 0 | 0 | 0 | 0 |
| 126.7436 | 37.82171 | 0 | Cropland | 0 | 1 | 0 | 0 | 0 | 0 | 0 |
| 126.7075 | 37.82444 | 0 | Cropland | 0 | 1 | 0 | 0 | 0 | 0 | 0 |
| 126.7425 | 37.83313 | 0 | Cropland | 0 | 1 | 0 | 0 | 0 | 0 | 0 |
| 126.6131 | 37.83493 | 0 | Cropland | 0 | 1 | 0 | 0 | 0 | 0 | 0 |
| 126.7225 | 37.83521 | 0 | Cropland | 0 | 1 | 0 | 0 | 0 | 0 | 0 |
| 126.7532 | 37.8434 | 0 | Cropland | 0 | 1 | 0 | 0 | 0 | 0 | 0 |
| 126.7739 | 37.85158 | 0 | Cropland | 0 | 1 | 0 | 0 | 0 | 0 | 0 |
| 126.7442 | 37.88696 | 0 | Cropland | 0 | 1 | 0 | 0 | 0 | 0 | 0 |
| 126.7633 | 37.89222 | 0 | Cropland | 0 | 1 | 0 | 0 | 0 | 0 | 0 |
| 126.7648 | 37.89858 | 0 | Cropland | 0 | 1 | 0 | 0 | 0 | 0 | 0 |
| 126.7669 | 37.93651 | 0 | Cropland | 0 | 1 | 0 | 0 | 0 | 0 | 0 |
| 126.3656 | 37.94107 | 0 | Cropland | 0 | 1 | 0 | 0 | 0 | 0 | 0 |
| 124.717 | 37.9534 | 0 | Cropland | 0 | 1 | 0 | 0 | 0 | 0 | 0 |
| 126.6328 | 37.966 | 0 | Cropland | 0 | 1 | 0 | 0 | 0 | 0 | 0 |
| 126.6574 | 37.99915 | 0 | Cropland | 0 | 1 | 0 | 0 | 0 | 0 | 0 |
| 127.0286 | 38.02444 | 0 | Cropland | 0 | 1 | 0 | 0 | 0 | 0 | 0 |
| 127.0598 | 38.03437 | 0 | Cropland | 0 | 1 | 0 | 0 | 0 | 0 | 0 |
| 127.0711 | 38.0375 | 0 | Cropland | 0 | 1 | 0 | 0 | 0 | 0 | 0 |
| 127.0303 | 38.04167 | 0 | Cropland | 0 | 1 | 0 | 0 | 0 | 0 | 0 |
| 127.0654 | 38.04317 | 0 | Cropland | 0 | 1 | 0 | 0 | 0 | 0 | 0 |
| 127.0133 | 38.0467 | 0 | Cropland | 0 | 1 | 0 | 0 | 0 | 0 | 0 |
| 127.0776 | 38.06301 | 0 | Cropland | 0 | 1 | 0 | 0 | 0 | 0 | 0 |
| 127.0726 | 38.06845 | 0 | Cropland | 0 | 1 | 0 | 0 | 0 | 0 | 0 |
| 127.074 | 38.13483 | 0 | Cropland | 0 | 1 | 0 | 0 | 0 | 0 | 0 |
| 127.0894 | 38.14123 | 0 | Cropland | 0 | 1 | 0 | 0 | 0 | 0 | 0 |
| 127.2789 | 38.18155 | 0 | Cropland | 0 | 1 | 0 | 0 | 0 | 0 | 0 |
| 127.1123 | 38.18544 | 0 | Cropland | 0 | 1 | 0 | 0 | 0 | 0 | 0 |
| 127.2674 | 38.18589 | 0 | Cropland | 0 | 1 | 0 | 0 | 0 | 0 | 0 |
| 127.2649 | 38.19267 | 0 | Cropland | 0 | 1 | 0 | 0 | 0 | 0 | 0 |
| 127.2547 | 38.20003 | 0 | Cropland | 0 | 1 | 0 | 0 | 0 | 0 | 0 |
| 127.2396 | 38.20013 | 0 | Cropland | 0 | 1 | 0 | 0 | 0 | 0 | 0 |
| 127.2476 | 38.20059 | 0 | Cropland | 0 | 1 | 0 | 0 | 0 | 0 | 0 |
| 127.2319 | 38.21772 | 0 | Cropland | 0 | 1 | 0 | 0 | 0 | 0 | 0 |
| 127.2432 | 38.22216 | 0 | Cropland | 0 | 1 | 0 | 0 | 0 | 0 | 0 |
| 127.1739 | 38.25711 | 0 | Cropland | 0 | 1 | 0 | 0 | 0 | 0 | 0 |
| 127.2263 | 38.3579 | 0 | Cropland | 0 | 1 | 0 | 0 | 0 | 0 | 0 |
| 125.214 | 38.80432 | 0 | Cropland | 0 | 1 | 0 | 0 | 0 | 0 | 0 |
| 125.2572 | 38.81136 | 0 | Cropland | 0 | 1 | 0 | 0 | 0 | 0 | 0 |
| 125.3213 | 38.82774 | 0 | Cropland | 0 | 1 | 0 | 0 | 0 | 0 | 0 |
| 125.3823 | 38.83077 | 0 | Cropland | 0 | 1 | 0 | 0 | 0 | 0 | 0 |
| 125.3007 | 38.83453 | 0 | Cropland | 0 | 1 | 0 | 0 | 0 | 0 | 0 |
| 125.2583 | 38.88863 | 0 | Cropland | 0 | 1 | 0 | 0 | 0 | 0 | 0 |
| 125.2321 | 38.94614 | 0 | Cropland | 0 | 1 | 0 | 0 | 0 | 0 | 0 |
| 125.2099 | 38.97502 | 0 | Cropland | 0 | 1 | 0 | 0 | 0 | 0 | 0 |
| 125.2169 | 38.99347 | 0 | Cropland | 0 | 1 | 0 | 0 | 0 | 0 | 0 |
| 125.2891 | 38.99969 | 0 | Cropland | 0 | 1 | 0 | 0 | 0 | 0 | 0 |
| 121.9571 | 39.19975 | 0 | Cropland | 0 | 1 | 0 | 0 | 0 | 0 | 0 |
| 122.2793 | 39.48047 | 0 | Cropland | 0 | 1 | 0 | 0 | 0 | 0 | 0 |
| 122.6991 | 39.61137 | 0 | Cropland | 0 | 1 | 0 | 0 | 0 | 0 | 0 |
| 124.2649 | 39.88214 | 0 | Cropland | 0 | 1 | 0 | 0 | 0 | 0 | 0 |
| 124.3613 | 39.89522 | 0 | Cropland | 0 | 1 | 0 | 0 | 0 | 0 | 0 |
| 124.3572 | 39.9017 | 0 | Cropland | 0 | 1 | 0 | 0 | 0 | 0 | 0 |
| 124.2774 | 39.90321 | 0 | Cropland | 0 | 1 | 0 | 0 | 0 | 0 | 0 |
| 124.2028 | 39.9243 | 0 | Cropland | 0 | 1 | 0 | 0 | 0 | 0 | 0 |
| 124.3634 | 39.93805 | 0 | Cropland | 0 | 1 | 0 | 0 | 0 | 0 | 0 |
| 124.3333 | 39.96819 | 0 | Cropland | 0 | 1 | 0 | 0 | 0 | 0 | 0 |
| 124.2944 | 39.98778 | 0 | Cropland | 0 | 1 | 0 | 0 | 0 | 0 | 0 |
| 125.6829 | 40.80969 | 0 | Cropland | 0 | 1 | 0 | 0 | 0 | 0 | 0 |
| 123.2651 | 41.87329 | 0 | Cropland | 0 | 1 | 0 | 0 | 0 | 0 | 0 |
| 123.3117 | 41.90775 | 0 | Cropland | 0 | 1 | 0 | 0 | 0 | 0 | 0 |
| 123.4917 | 41.90926 | 0 | Cropland | 0 | 1 | 0 | 0 | 0 | 0 | 0 |
| 123.3041 | 41.91205 | 0 | Cropland | 0 | 1 | 0 | 0 | 0 | 0 | 0 |
| 123.4667 | 41.9346 | 0 | Cropland | 0 | 1 | 0 | 0 | 0 | 0 | 0 |
| 123.4271 | 41.9694 | 0 | Cropland | 0 | 1 | 0 | 0 | 0 | 0 | 0 |
| 123.2129 | 41.99225 | 0 | Cropland | 0 | 1 | 0 | 0 | 0 | 0 | 0 |
| 130.3908 | 42.33883 | 0 | Cropland | 0 | 1 | 0 | 0 | 0 | 0 | 0 |
| 130.3578 | 42.35056 | 0 | Cropland | 0 | 1 | 0 | 0 | 0 | 0 | 0 |
| 127.044 | 36.31247 | 1 | Cropland / Other vegetation mosaic | 0 | 1 | 0 | 0 | 0 | 0 | 0 |
| 126.8551 | 36.83974 | 1 | Cropland / Other vegetation mosaic | 0 | 1 | 0 | 0 | 0 | 0 | 0 |
| 127.643 | 37.07308 | 1 | Cropland / Other vegetation mosaic | 0 | 1 | 0 | 0 | 0 | 0 | 0 |
| 126.7307 | 37.1466 | 1 | Cropland / Other vegetation mosaic | 0 | 1 | 0 | 0 | 0 | 0 | 0 |
| 126.5184 | 37.62179 | 1 | Cropland / Other vegetation mosaic | 0 | 1 | 0 | 0 | 0 | 0 | 0 |
| 126.246 | 37.63292 | 1 | Cropland / Other vegetation mosaic | 0 | 1 | 0 | 0 | 0 | 0 | 0 |
| 126.7422 | 37.90653 | 1 | Cropland / Other vegetation mosaic | 0 | 1 | 0 | 0 | 0 | 0 | 0 |
| 127.1501 | 38.01774 | 1 | Cropland / Other vegetation mosaic | 0 | 1 | 0 | 0 | 0 | 0 | 0 |
| 126.5372 | 36.62244 | 1 | Cropland / Other vegetation mosaic | 0 | 1 | 0 | 0 | 0 | 0 | 0 |
| 126.5516 | 36.75896 | 1 | Cropland / Other vegetation mosaic | 0 | 1 | 0 | 0 | 0 | 0 | 0 |
| 126.9064 | 36.26364 | 1 | Cropland / Other vegetation mosaic | 0 | 1 | 0 | 0 | 0 | 0 | 0 |
| 126.6175 | 33.4488 | 0 | Cropland / Other vegetation mosaic | 0 | 1 | 0 | 0 | 0 | 0 | 0 |
| 126.7467 | 33.45957 | 0 | Cropland / Other vegetation mosaic | 0 | 1 | 0 | 0 | 0 | 0 | 0 |
| 126.7495 | 33.46513 | 0 | Cropland / Other vegetation mosaic | 0 | 1 | 0 | 0 | 0 | 0 | 0 |
| 128.7858 | 35.30001 | 0 | Cropland / Other vegetation mosaic | 0 | 1 | 0 | 0 | 0 | 0 | 0 |
| 129.2878 | 35.34395 | 0 | Cropland / Other vegetation mosaic | 0 | 1 | 0 | 0 | 0 | 0 | 0 |
| 129.3219 | 35.35861 | 0 | Cropland / Other vegetation mosaic | 0 | 1 | 0 | 0 | 0 | 0 | 0 |
| 127.811 | 35.64296 | 0 | Cropland / Other vegetation mosaic | 0 | 1 | 0 | 0 | 0 | 0 | 0 |
| 127.8266 | 35.66364 | 0 | Cropland / Other vegetation mosaic | 0 | 1 | 0 | 0 | 0 | 0 | 0 |
| 126.7767 | 35.92278 | 0 | Cropland / Other vegetation mosaic | 0 | 1 | 0 | 0 | 0 | 0 | 0 |
| 126.9058 | 35.95332 | 0 | Cropland / Other vegetation mosaic | 0 | 1 | 0 | 0 | 0 | 0 | 0 |
| 128.319 | 36.22246 | 0 | Cropland / Other vegetation mosaic | 0 | 1 | 0 | 0 | 0 | 0 | 0 |
| 128.3566 | 36.28904 | 0 | Cropland / Other vegetation mosaic | 0 | 1 | 0 | 0 | 0 | 0 | 0 |
| 127.0159 | 36.32133 | 0 | Cropland / Other vegetation mosaic | 0 | 1 | 0 | 0 | 0 | 0 | 0 |
| 126.4079 | 36.43654 | 0 | Cropland / Other vegetation mosaic | 0 | 1 | 0 | 0 | 0 | 0 | 0 |
| 126.4178 | 36.4375 | 0 | Cropland / Other vegetation mosaic | 0 | 1 | 0 | 0 | 0 | 0 | 0 |
| 126.4153 | 36.44194 | 0 | Cropland / Other vegetation mosaic | 0 | 1 | 0 | 0 | 0 | 0 | 0 |
| 128.2017 | 36.58032 | 0 | Cropland / Other vegetation mosaic | 0 | 1 | 0 | 0 | 0 | 0 | 0 |
| 127.8247 | 36.82819 | 0 | Cropland / Other vegetation mosaic | 0 | 1 | 0 | 0 | 0 | 0 | 0 |
| 127.4809 | 36.84099 | 0 | Cropland / Other vegetation mosaic | 0 | 1 | 0 | 0 | 0 | 0 | 0 |
| 126.9046 | 36.85713 | 0 | Cropland / Other vegetation mosaic | 0 | 1 | 0 | 0 | 0 | 0 | 0 |
| 127.5062 | 36.86237 | 0 | Cropland / Other vegetation mosaic | 0 | 1 | 0 | 0 | 0 | 0 | 0 |
| 126.9417 | 36.89695 | 0 | Cropland / Other vegetation mosaic | 0 | 1 | 0 | 0 | 0 | 0 | 0 |
| 127.7769 | 36.94456 | 0 | Cropland / Other vegetation mosaic | 0 | 1 | 0 | 0 | 0 | 0 | 0 |
| 127.4557 | 36.94857 | 0 | Cropland / Other vegetation mosaic | 0 | 1 | 0 | 0 | 0 | 0 | 0 |
| 127.7822 | 36.95389 | 0 | Cropland / Other vegetation mosaic | 0 | 1 | 0 | 0 | 0 | 0 | 0 |
| 127.7714 | 36.96396 | 0 | Cropland / Other vegetation mosaic | 0 | 1 | 0 | 0 | 0 | 0 | 0 |
| 127.7769 | 36.96611 | 0 | Cropland / Other vegetation mosaic | 0 | 1 | 0 | 0 | 0 | 0 | 0 |
| 127.8076 | 36.98184 | 0 | Cropland / Other vegetation mosaic | 0 | 1 | 0 | 0 | 0 | 0 | 0 |
| 127.5399 | 37.03553 | 0 | Cropland / Other vegetation mosaic | 0 | 1 | 0 | 0 | 0 | 0 | 0 |
| 127.4343 | 37.08379 | 0 | Cropland / Other vegetation mosaic | 0 | 1 | 0 | 0 | 0 | 0 | 0 |
| 127.1719 | 37.11583 | 0 | Cropland / Other vegetation mosaic | 0 | 1 | 0 | 0 | 0 | 0 | 0 |
| 126.865 | 37.2493 | 0 | Cropland / Other vegetation mosaic | 0 | 1 | 0 | 0 | 0 | 0 | 0 |
| 127.809 | 37.28124 | 0 | Cropland / Other vegetation mosaic | 0 | 1 | 0 | 0 | 0 | 0 | 0 |
| 127.5571 | 37.28339 | 0 | Cropland / Other vegetation mosaic | 0 | 1 | 0 | 0 | 0 | 0 | 0 |
| 127.8272 | 37.31583 | 0 | Cropland / Other vegetation mosaic | 0 | 1 | 0 | 0 | 0 | 0 | 0 |
| 126.3916 | 37.44514 | 0 | Cropland / Other vegetation mosaic | 0 | 1 | 0 | 0 | 0 | 0 | 0 |
| 126.393 | 37.44973 | 0 | Cropland / Other vegetation mosaic | 0 | 1 | 0 | 0 | 0 | 0 | 0 |
| 126.5012 | 37.51642 | 0 | Cropland / Other vegetation mosaic | 0 | 1 | 0 | 0 | 0 | 0 | 0 |
| 126.5802 | 37.60594 | 0 | Cropland / Other vegetation mosaic | 0 | 1 | 0 | 0 | 0 | 0 | 0 |
| 126.5645 | 37.62001 | 0 | Cropland / Other vegetation mosaic | 0 | 1 | 0 | 0 | 0 | 0 | 0 |
| 126.5994 | 37.6249 | 0 | Cropland / Other vegetation mosaic | 0 | 1 | 0 | 0 | 0 | 0 | 0 |
| 126.5225 | 37.63972 | 0 | Cropland / Other vegetation mosaic | 0 | 1 | 0 | 0 | 0 | 0 | 0 |
| 126.484 | 37.66302 | 0 | Cropland / Other vegetation mosaic | 0 | 1 | 0 | 0 | 0 | 0 | 0 |
| 126.3978 | 37.69632 | 0 | Cropland / Other vegetation mosaic | 0 | 1 | 0 | 0 | 0 | 0 | 0 |
| 126.3112 | 37.70625 | 0 | Cropland / Other vegetation mosaic | 0 | 1 | 0 | 0 | 0 | 0 | 0 |
| 126.3053 | 37.70893 | 0 | Cropland / Other vegetation mosaic | 0 | 1 | 0 | 0 | 0 | 0 | 0 |
| 126.3169 | 37.7106 | 0 | Cropland / Other vegetation mosaic | 0 | 1 | 0 | 0 | 0 | 0 | 0 |
| 126.3074 | 37.72151 | 0 | Cropland / Other vegetation mosaic | 0 | 1 | 0 | 0 | 0 | 0 | 0 |
| 126.3147 | 37.72827 | 0 | Cropland / Other vegetation mosaic | 0 | 1 | 0 | 0 | 0 | 0 | 0 |
| 126.3639 | 37.73131 | 0 | Cropland / Other vegetation mosaic | 0 | 1 | 0 | 0 | 0 | 0 | 0 |
| 126.3847 | 37.735 | 0 | Cropland / Other vegetation mosaic | 0 | 1 | 0 | 0 | 0 | 0 | 0 |
| 126.3978 | 37.75333 | 0 | Cropland / Other vegetation mosaic | 0 | 1 | 0 | 0 | 0 | 0 | 0 |
| 126.5504 | 37.76 | 0 | Cropland / Other vegetation mosaic | 0 | 1 | 0 | 0 | 0 | 0 | 0 |
| 126.4319 | 37.76694 | 0 | Cropland / Other vegetation mosaic | 0 | 1 | 0 | 0 | 0 | 0 | 0 |
| 126.8111 | 37.79616 | 0 | Cropland / Other vegetation mosaic | 0 | 1 | 0 | 0 | 0 | 0 | 0 |
| 126.4081 | 37.79777 | 0 | Cropland / Other vegetation mosaic | 0 | 1 | 0 | 0 | 0 | 0 | 0 |
| 126.7556 | 37.88583 | 0 | Cropland / Other vegetation mosaic | 0 | 1 | 0 | 0 | 0 | 0 | 0 |
| 126.771 | 37.89809 | 0 | Cropland / Other vegetation mosaic | 0 | 1 | 0 | 0 | 0 | 0 | 0 |
| 124.6831 | 37.94889 | 0 | Cropland / Other vegetation mosaic | 0 | 1 | 0 | 0 | 0 | 0 | 0 |
| 124.6328 | 37.95083 | 0 | Cropland / Other vegetation mosaic | 0 | 1 | 0 | 0 | 0 | 0 | 0 |
| 127.0419 | 38.02905 | 0 | Cropland / Other vegetation mosaic | 0 | 1 | 0 | 0 | 0 | 0 | 0 |
| 127.0153 | 38.03766 | 0 | Cropland / Other vegetation mosaic | 0 | 1 | 0 | 0 | 0 | 0 | 0 |
| 127.0358 | 38.10663 | 0 | Cropland / Other vegetation mosaic | 0 | 1 | 0 | 0 | 0 | 0 | 0 |
| 127.2635 | 38.17051 | 0 | Cropland / Other vegetation mosaic | 0 | 1 | 0 | 0 | 0 | 0 | 0 |
| 127.244 | 38.22799 | 0 | Cropland / Other vegetation mosaic | 0 | 1 | 0 | 0 | 0 | 0 | 0 |
| 127.2533 | 38.23727 | 0 | Cropland / Other vegetation mosaic | 0 | 1 | 0 | 0 | 0 | 0 | 0 |
| 127.2355 | 38.24151 | 0 | Cropland / Other vegetation mosaic | 0 | 1 | 0 | 0 | 0 | 0 | 0 |
| 127.2549 | 38.25303 | 0 | Cropland / Other vegetation mosaic | 0 | 1 | 0 | 0 | 0 | 0 | 0 |
| 127.282 | 38.25913 | 0 | Cropland / Other vegetation mosaic | 0 | 1 | 0 | 0 | 0 | 0 | 0 |
| 127.18 | 38.26102 | 0 | Cropland / Other vegetation mosaic | 0 | 1 | 0 | 0 | 0 | 0 | 0 |
| 127.5293 | 38.39356 | 0 | Cropland / Other vegetation mosaic | 0 | 1 | 0 | 0 | 0 | 0 | 0 |
| 130.3908 | 42.30389 | 0 | Cropland / Other vegetation mosaic | 0 | 1 | 0 | 0 | 0 | 0 | 0 |
| 130.3667 | 42.33528 | 0 | Cropland / Other vegetation mosaic | 0 | 1 | 0 | 0 | 0 | 0 | 0 |
| 130.5864 | 42.34583 | 0 | Cropland / Other vegetation mosaic | 0 | 1 | 0 | 0 | 0 | 0 | 0 |
| 130.5621 | 42.36698 | 0 | Cropland / Other vegetation mosaic | 0 | 1 | 0 | 0 | 0 | 0 | 0 |
| 130.5215 | 42.36794 | 0 | Cropland / Other vegetation mosaic | 0 | 1 | 0 | 0 | 0 | 0 | 0 |
| 126.7164 | 37.65857 | 1 | Herbaceous vegetation | 0 | 0 | 0 | 1 | 0 | 0 | 0 |
| 126.7658 | 37.842 | 1 | Herbaceous vegetation | 0 | 0 | 0 | 1 | 0 | 0 | 0 |
| 126.2258 | 36.87961 | 1 | Herbaceous vegetation | 0 | 0 | 0 | 1 | 0 | 0 | 0 |
| 128.2284 | 36.18512 | 0 | Herbaceous vegetation | 0 | 0 | 0 | 1 | 0 | 0 | 0 |
| 127.8099 | 36.89626 | 0 | Herbaceous vegetation | 0 | 0 | 0 | 1 | 0 | 0 | 0 |
| 126.8533 | 37.2725 | 0 | Herbaceous vegetation | 0 | 0 | 0 | 1 | 0 | 0 | 0 |
| 126.5062 | 37.52601 | 0 | Herbaceous vegetation | 0 | 0 | 0 | 1 | 0 | 0 | 0 |
| 126.666 | 37.69212 | 0 | Herbaceous vegetation | 0 | 0 | 0 | 1 | 0 | 0 | 0 |
| 126.5586 | 37.73174 | 0 | Herbaceous vegetation | 0 | 0 | 0 | 1 | 0 | 0 | 0 |
| 126.6895 | 37.76167 | 0 | Herbaceous vegetation | 0 | 0 | 0 | 1 | 0 | 0 | 0 |
| 126.7741 | 37.81113 | 0 | Herbaceous vegetation | 0 | 0 | 0 | 1 | 0 | 0 | 0 |
| 126.6544 | 37.82536 | 0 | Herbaceous vegetation | 0 | 0 | 0 | 1 | 0 | 0 | 0 |
| 126.3708 | 37.95551 | 0 | Herbaceous vegetation | 0 | 0 | 0 | 1 | 0 | 0 | 0 |
| 125.3433 | 38.9286 | 0 | Herbaceous vegetation | 0 | 0 | 0 | 1 | 0 | 0 | 0 |
| 130.2806 | 42.24503 | 0 | Herbaceous vegetation | 0 | 0 | 0 | 1 | 0 | 0 | 0 |
| 127.1809 | 38.04144 | 1 | Mixed forest | 1 | 0 | 0 | 0 | 0 | 0 | 0 |
| 126.5996 | 36.72264 | 1 | Mixed forest | 1 | 0 | 0 | 0 | 0 | 0 | 0 |
| 126.7195 | 37.55029 | 1 | Mixed forest | 1 | 0 | 0 | 0 | 0 | 0 | 0 |
| 126.8711 | 36.3818 | 1 | Mixed forest | 1 | 0 | 0 | 0 | 0 | 0 | 0 |
| 127.1911 | 38.12016 | 1 | Mixed forest | 1 | 0 | 0 | 0 | 0 | 0 | 0 |
| 127.2829 | 36.51571 | 1 | Mixed forest | 1 | 0 | 0 | 0 | 0 | 0 | 0 |
| 127.303 | 36.4282 | 1 | Mixed forest | 1 | 0 | 0 | 0 | 0 | 0 | 0 |
| 127.385 | 37.1861 | 1 | Mixed forest | 1 | 0 | 0 | 0 | 0 | 0 | 0 |
| 127.5741 | 36.61807 | 1 | Mixed forest | 1 | 0 | 0 | 0 | 0 | 0 | 0 |
| 129.2116 | 35.26937 | 0 | Mixed forest | 1 | 0 | 0 | 0 | 0 | 0 | 0 |
| 128.8568 | 35.36907 | 0 | Mixed forest | 1 | 0 | 0 | 0 | 0 | 0 | 0 |
| 128.1538 | 35.65 | 0 | Mixed forest | 1 | 0 | 0 | 0 | 0 | 0 | 0 |
| 127.8713 | 35.76239 | 0 | Mixed forest | 1 | 0 | 0 | 0 | 0 | 0 | 0 |
| 127.802 | 35.76637 | 0 | Mixed forest | 1 | 0 | 0 | 0 | 0 | 0 | 0 |
| 128.0318 | 35.79904 | 0 | Mixed forest | 1 | 0 | 0 | 0 | 0 | 0 | 0 |
| 128.9247 | 35.83899 | 0 | Mixed forest | 1 | 0 | 0 | 0 | 0 | 0 | 0 |
| 127.9141 | 35.9505 | 0 | Mixed forest | 1 | 0 | 0 | 0 | 0 | 0 | 0 |
| 127.2756 | 36.0575 | 0 | Mixed forest | 1 | 0 | 0 | 0 | 0 | 0 | 0 |
| 126.7217 | 36.17662 | 0 | Mixed forest | 1 | 0 | 0 | 0 | 0 | 0 | 0 |
| 128.3534 | 36.33581 | 0 | Mixed forest | 1 | 0 | 0 | 0 | 0 | 0 | 0 |
| 127.1874 | 36.33969 | 0 | Mixed forest | 1 | 0 | 0 | 0 | 0 | 0 | 0 |
| 128.2505 | 36.34389 | 0 | Mixed forest | 1 | 0 | 0 | 0 | 0 | 0 | 0 |
| 128.1755 | 36.35064 | 0 | Mixed forest | 1 | 0 | 0 | 0 | 0 | 0 | 0 |
| 128.0618 | 36.43699 | 0 | Mixed forest | 1 | 0 | 0 | 0 | 0 | 0 | 0 |
| 128.0851 | 36.52896 | 0 | Mixed forest | 1 | 0 | 0 | 0 | 0 | 0 | 0 |
| 128.0925 | 36.82404 | 0 | Mixed forest | 1 | 0 | 0 | 0 | 0 | 0 | 0 |
| 127.9683 | 36.82615 | 0 | Mixed forest | 1 | 0 | 0 | 0 | 0 | 0 | 0 |
| 128.9694 | 36.83726 | 0 | Mixed forest | 1 | 0 | 0 | 0 | 0 | 0 | 0 |
| 127.8264 | 36.92003 | 0 | Mixed forest | 1 | 0 | 0 | 0 | 0 | 0 | 0 |
| 127.6659 | 36.94743 | 0 | Mixed forest | 1 | 0 | 0 | 0 | 0 | 0 | 0 |
| 127.4078 | 36.98717 | 0 | Mixed forest | 1 | 0 | 0 | 0 | 0 | 0 | 0 |
| 127.4208 | 37.00366 | 0 | Mixed forest | 1 | 0 | 0 | 0 | 0 | 0 | 0 |
| 128.552 | 37.21397 | 0 | Mixed forest | 1 | 0 | 0 | 0 | 0 | 0 | 0 |
| 127.8843 | 37.24433 | 0 | Mixed forest | 1 | 0 | 0 | 0 | 0 | 0 | 0 |
| 128.503 | 37.25875 | 0 | Mixed forest | 1 | 0 | 0 | 0 | 0 | 0 | 0 |
| 128.468 | 37.26264 | 0 | Mixed forest | 1 | 0 | 0 | 0 | 0 | 0 | 0 |
| 127.9712 | 37.28374 | 0 | Mixed forest | 1 | 0 | 0 | 0 | 0 | 0 | 0 |
| 128.8464 | 37.66583 | 0 | Mixed forest | 1 | 0 | 0 | 0 | 0 | 0 | 0 |
| 127.5259 | 37.67411 | 0 | Mixed forest | 1 | 0 | 0 | 0 | 0 | 0 | 0 |
| 126.3819 | 37.70472 | 0 | Mixed forest | 1 | 0 | 0 | 0 | 0 | 0 | 0 |
| 127.4799 | 37.72349 | 0 | Mixed forest | 1 | 0 | 0 | 0 | 0 | 0 | 0 |
| 124.6893 | 37.83264 | 0 | Mixed forest | 1 | 0 | 0 | 0 | 0 | 0 | 0 |
| 127.6196 | 37.90357 | 0 | Mixed forest | 1 | 0 | 0 | 0 | 0 | 0 | 0 |
| 124.6432 | 37.92642 | 0 | Mixed forest | 1 | 0 | 0 | 0 | 0 | 0 | 0 |
| 127.6083 | 37.93403 | 0 | Mixed forest | 1 | 0 | 0 | 0 | 0 | 0 | 0 |
| 127.635 | 37.9766 | 0 | Mixed forest | 1 | 0 | 0 | 0 | 0 | 0 | 0 |
| 127.1089 | 37.98119 | 0 | Mixed forest | 1 | 0 | 0 | 0 | 0 | 0 | 0 |
| 127.1812 | 38.0957 | 0 | Mixed forest | 1 | 0 | 0 | 0 | 0 | 0 | 0 |
| 127.1886 | 38.12254 | 0 | Mixed forest | 1 | 0 | 0 | 0 | 0 | 0 | 0 |
| 126.9737 | 37.45766 | 1 | Needleleaf deciduous forest | 1 | 0 | 0 | 0 | 0 | 0 | 0 |
| 127.0843 | 38.184 | 1 | Needleleaf deciduous forest | 1 | 0 | 0 | 0 | 0 | 0 | 0 |
| 126.5718 | 36.70259 | 1 | Needleleaf deciduous forest | 1 | 0 | 0 | 0 | 0 | 0 | 0 |
| 127.4111 | 36.80262 | 1 | Needleleaf deciduous forest | 1 | 0 | 0 | 0 | 0 | 0 | 0 |
| 127.5842 | 36.68244 | 0 | Needleleaf deciduous forest | 1 | 0 | 0 | 0 | 0 | 0 | 0 |
| 128.8131 | 36.99832 | 0 | Needleleaf deciduous forest | 1 | 0 | 0 | 0 | 0 | 0 | 0 |
| 125.9518 | 37.08121 | 0 | Needleleaf deciduous forest | 1 | 0 | 0 | 0 | 0 | 0 | 0 |
| 128.7026 | 37.3991 | 0 | Needleleaf deciduous forest | 1 | 0 | 0 | 0 | 0 | 0 | 0 |
| 127.6129 | 37.98348 | 0 | Needleleaf deciduous forest | 1 | 0 | 0 | 0 | 0 | 0 | 0 |
| 123.5861 | 41.83972 | 0 | Needleleaf deciduous forest | 1 | 0 | 0 | 0 | 0 | 0 | 0 |
| 130.5508 | 42.34637 | 0 | Needleleaf deciduous forest | 1 | 0 | 0 | 0 | 0 | 0 | 0 |
| 130.2078 | 42.3488 | 0 | Needleleaf deciduous forest | 1 | 0 | 0 | 0 | 0 | 0 | 0 |
| 127.1423 | 38.07996 | 1 | Needleleaf evergreen forest | 1 | 0 | 0 | 0 | 0 | 0 | 0 |
| 126.2559 | 36.83056 | 1 | Needleleaf evergreen forest | 1 | 0 | 0 | 0 | 0 | 0 | 0 |
| 126.5277 | 36.39892 | 1 | Needleleaf evergreen forest | 1 | 0 | 0 | 0 | 0 | 0 | 0 |
| 126.8009 | 36.35416 | 1 | Needleleaf evergreen forest | 1 | 0 | 0 | 0 | 0 | 0 | 0 |
| 128.087 | 35.5631 | 1 | Needleleaf evergreen forest | 1 | 0 | 0 | 0 | 0 | 0 | 0 |
| 128.072 | 35.6089 | 0 | Needleleaf evergreen forest | 1 | 0 | 0 | 0 | 0 | 0 | 0 |
| 128.1502 | 35.69586 | 0 | Needleleaf evergreen forest | 1 | 0 | 0 | 0 | 0 | 0 | 0 |
| 127.8639 | 35.85916 | 0 | Needleleaf evergreen forest | 1 | 0 | 0 | 0 | 0 | 0 | 0 |
| 128.2954 | 36.26231 | 0 | Needleleaf evergreen forest | 1 | 0 | 0 | 0 | 0 | 0 | 0 |
| 126.6465 | 36.80577 | 0 | Needleleaf evergreen forest | 1 | 0 | 0 | 0 | 0 | 0 | 0 |
| 128.7119 | 37.35972 | 0 | Needleleaf evergreen forest | 1 | 0 | 0 | 0 | 0 | 0 | 0 |
| 126.4656 | 37.62222 | 0 | Needleleaf evergreen forest | 1 | 0 | 0 | 0 | 0 | 0 | 0 |
| 127.0963 | 37.64581 | 0 | Needleleaf evergreen forest | 1 | 0 | 0 | 0 | 0 | 0 | 0 |
| 126.3049 | 37.78113 | 0 | Needleleaf evergreen forest | 1 | 0 | 0 | 0 | 0 | 0 | 0 |
| 128.408 | 38.50338 | 0 | Needleleaf evergreen forest | 1 | 0 | 0 | 0 | 0 | 0 | 0 |
| 127.1388 | 36.35927 | 1 | Open canopy forest | 1 | 0 | 0 | 0 | 0 | 0 | 0 |
| 127.0572 | 36.38565 | 1 | Open canopy forest | 1 | 0 | 0 | 0 | 0 | 0 | 0 |
| 127.4021 | 36.81985 | 1 | Open canopy forest | 1 | 0 | 0 | 0 | 0 | 0 | 0 |
| 127.3892 | 37.05043 | 1 | Open canopy forest | 1 | 0 | 0 | 0 | 0 | 0 | 0 |
| 126.7867 | 37.86831 | 1 | Open canopy forest | 1 | 0 | 0 | 0 | 0 | 0 | 0 |
| 126.5312 | 36.31683 | 1 | Open canopy forest | 1 | 0 | 0 | 0 | 0 | 0 | 0 |
| 126.777 | 37.81658 | 1 | Open canopy forest | 1 | 0 | 0 | 0 | 0 | 0 | 0 |
| 126.8268 | 37.82385 | 1 | Open canopy forest | 1 | 0 | 0 | 0 | 0 | 0 | 0 |
| 126.924 | 36.2179 | 1 | Open canopy forest | 1 | 0 | 0 | 0 | 0 | 0 | 0 |
| 126.9364 | 36.24751 | 1 | Open canopy forest | 1 | 0 | 0 | 0 | 0 | 0 | 0 |
| 127.2219 | 36.46994 | 1 | Open canopy forest | 1 | 0 | 0 | 0 | 0 | 0 | 0 |
| 127.2347 | 36.51818 | 1 | Open canopy forest | 1 | 0 | 0 | 0 | 0 | 0 | 0 |
| 126.9198 | 35.22317 | 0 | Open canopy forest | 1 | 0 | 0 | 0 | 0 | 0 | 0 |
| 129.2125 | 35.22931 | 0 | Open canopy forest | 1 | 0 | 0 | 0 | 0 | 0 | 0 |
| 128.4251 | 36.30497 | 0 | Open canopy forest | 1 | 0 | 0 | 0 | 0 | 0 | 0 |
| 126.4227 | 36.36846 | 0 | Open canopy forest | 1 | 0 | 0 | 0 | 0 | 0 | 0 |
| 126.406 | 36.36981 | 0 | Open canopy forest | 1 | 0 | 0 | 0 | 0 | 0 | 0 |
| 128.1138 | 36.38159 | 0 | Open canopy forest | 1 | 0 | 0 | 0 | 0 | 0 | 0 |
| 126.4212 | 36.40484 | 0 | Open canopy forest | 1 | 0 | 0 | 0 | 0 | 0 | 0 |
| 126.3984 | 36.42697 | 0 | Open canopy forest | 1 | 0 | 0 | 0 | 0 | 0 | 0 |
| 126.6342 | 36.60534 | 0 | Open canopy forest | 1 | 0 | 0 | 0 | 0 | 0 | 0 |
| 126.9959 | 36.80784 | 0 | Open canopy forest | 1 | 0 | 0 | 0 | 0 | 0 | 0 |
| 127.4744 | 36.84456 | 0 | Open canopy forest | 1 | 0 | 0 | 0 | 0 | 0 | 0 |
| 127.4176 | 36.85694 | 0 | Open canopy forest | 1 | 0 | 0 | 0 | 0 | 0 | 0 |
| 126.9524 | 36.8928 | 0 | Open canopy forest | 1 | 0 | 0 | 0 | 0 | 0 | 0 |
| 127.7952 | 36.90846 | 0 | Open canopy forest | 1 | 0 | 0 | 0 | 0 | 0 | 0 |
| 127.7473 | 36.91963 | 0 | Open canopy forest | 1 | 0 | 0 | 0 | 0 | 0 | 0 |
| 127.181 | 36.92384 | 0 | Open canopy forest | 1 | 0 | 0 | 0 | 0 | 0 | 0 |
| 127.8821 | 36.96655 | 0 | Open canopy forest | 1 | 0 | 0 | 0 | 0 | 0 | 0 |
| 127.5089 | 36.98548 | 0 | Open canopy forest | 1 | 0 | 0 | 0 | 0 | 0 | 0 |
| 127.7568 | 36.99547 | 0 | Open canopy forest | 1 | 0 | 0 | 0 | 0 | 0 | 0 |
| 127.7965 | 37.03615 | 0 | Open canopy forest | 1 | 0 | 0 | 0 | 0 | 0 | 0 |
| 127.5341 | 37.06277 | 0 | Open canopy forest | 1 | 0 | 0 | 0 | 0 | 0 | 0 |
| 128.4239 | 37.22833 | 0 | Open canopy forest | 1 | 0 | 0 | 0 | 0 | 0 | 0 |
| 128.4228 | 37.23778 | 0 | Open canopy forest | 1 | 0 | 0 | 0 | 0 | 0 | 0 |
| 128.4333 | 37.2521 | 0 | Open canopy forest | 1 | 0 | 0 | 0 | 0 | 0 | 0 |
| 128.9703 | 37.25972 | 0 | Open canopy forest | 1 | 0 | 0 | 0 | 0 | 0 | 0 |
| 127.7828 | 37.26149 | 0 | Open canopy forest | 1 | 0 | 0 | 0 | 0 | 0 | 0 |
| 127.7987 | 37.26531 | 0 | Open canopy forest | 1 | 0 | 0 | 0 | 0 | 0 | 0 |
| 128.7536 | 37.34889 | 0 | Open canopy forest | 1 | 0 | 0 | 0 | 0 | 0 | 0 |
| 128.7914 | 37.35361 | 0 | Open canopy forest | 1 | 0 | 0 | 0 | 0 | 0 | 0 |
| 128.7721 | 37.35623 | 0 | Open canopy forest | 1 | 0 | 0 | 0 | 0 | 0 | 0 |
| 126.8081 | 37.40222 | 0 | Open canopy forest | 1 | 0 | 0 | 0 | 0 | 0 | 0 |
| 128.297 | 37.42682 | 0 | Open canopy forest | 1 | 0 | 0 | 0 | 0 | 0 | 0 |
| 128.4025 | 37.42694 | 0 | Open canopy forest | 1 | 0 | 0 | 0 | 0 | 0 | 0 |
| 126.5386 | 37.50669 | 0 | Open canopy forest | 1 | 0 | 0 | 0 | 0 | 0 | 0 |
| 126.4369 | 37.63028 | 0 | Open canopy forest | 1 | 0 | 0 | 0 | 0 | 0 | 0 |
| 126.3262 | 37.70221 | 0 | Open canopy forest | 1 | 0 | 0 | 0 | 0 | 0 | 0 |
| 126.7481 | 37.73382 | 0 | Open canopy forest | 1 | 0 | 0 | 0 | 0 | 0 | 0 |
| 126.5257 | 37.76282 | 0 | Open canopy forest | 1 | 0 | 0 | 0 | 0 | 0 | 0 |
| 127.4639 | 37.77859 | 0 | Open canopy forest | 1 | 0 | 0 | 0 | 0 | 0 | 0 |
| 127.5834 | 37.81692 | 0 | Open canopy forest | 1 | 0 | 0 | 0 | 0 | 0 | 0 |
| 124.6922 | 37.93194 | 0 | Open canopy forest | 1 | 0 | 0 | 0 | 0 | 0 | 0 |
| 124.7023 | 37.93317 | 0 | Open canopy forest | 1 | 0 | 0 | 0 | 0 | 0 | 0 |
| 126.7523 | 37.97129 | 0 | Open canopy forest | 1 | 0 | 0 | 0 | 0 | 0 | 0 |
| 127.0467 | 38.00037 | 0 | Open canopy forest | 1 | 0 | 0 | 0 | 0 | 0 | 0 |
| 127.1539 | 38.01552 | 0 | Open canopy forest | 1 | 0 | 0 | 0 | 0 | 0 | 0 |
| 127.029 | 38.03303 | 0 | Open canopy forest | 1 | 0 | 0 | 0 | 0 | 0 | 0 |
| 127.0826 | 38.0478 | 0 | Open canopy forest | 1 | 0 | 0 | 0 | 0 | 0 | 0 |
| 127.0199 | 38.09222 | 0 | Open canopy forest | 1 | 0 | 0 | 0 | 0 | 0 | 0 |
| 127.0488 | 38.1076 | 0 | Open canopy forest | 1 | 0 | 0 | 0 | 0 | 0 | 0 |
| 127.0414 | 38.11927 | 0 | Open canopy forest | 1 | 0 | 0 | 0 | 0 | 0 | 0 |
| 127.0622 | 38.13072 | 0 | Open canopy forest | 1 | 0 | 0 | 0 | 0 | 0 | 0 |
| 127.2782 | 38.1702 | 0 | Open canopy forest | 1 | 0 | 0 | 0 | 0 | 0 | 0 |
| 127.2677 | 38.32005 | 0 | Open canopy forest | 1 | 0 | 0 | 0 | 0 | 0 | 0 |
| 127.128 | 38.3215 | 0 | Open canopy forest | 1 | 0 | 0 | 0 | 0 | 0 | 0 |
| 125.3533 | 38.83667 | 0 | Open canopy forest | 1 | 0 | 0 | 0 | 0 | 0 | 0 |
| 125.3534 | 38.90713 | 0 | Open canopy forest | 1 | 0 | 0 | 0 | 0 | 0 | 0 |
| 124.2451 | 39.87194 | 0 | Open canopy forest | 1 | 0 | 0 | 0 | 0 | 0 | 0 |
| 130.405 | 42.33016 | 0 | Open canopy forest | 1 | 0 | 0 | 0 | 0 | 0 | 0 |
| 130.4872 | 42.352 | 0 | Open canopy forest | 1 | 0 | 0 | 0 | 0 | 0 | 0 |
| 130.3557 | 42.36448 | 0 | Open canopy forest | 1 | 0 | 0 | 0 | 0 | 0 | 0 |
| 127.0658 | 35.80614 | 1 | Paddy field | 0 | 1 | 0 | 0 | 0 | 0 | 1 |
| 126.8991 | 36.32033 | 1 | Paddy field | 0 | 1 | 0 | 0 | 0 | 0 | 1 |
| 126.9274 | 36.33966 | 1 | Paddy field | 0 | 1 | 0 | 0 | 0 | 0 | 1 |
| 126.9131 | 36.36726 | 1 | Paddy field | 0 | 1 | 0 | 0 | 0 | 0 | 1 |
| 126.9581 | 36.36758 | 1 | Paddy field | 0 | 1 | 0 | 0 | 0 | 0 | 1 |
| 126.9571 | 36.38873 | 1 | Paddy field | 0 | 1 | 0 | 0 | 0 | 0 | 1 |
| 126.8173 | 36.81695 | 1 | Paddy field | 0 | 1 | 0 | 0 | 0 | 0 | 1 |
| 127.7579 | 37.16133 | 1 | Paddy field | 0 | 1 | 0 | 0 | 0 | 0 | 1 |
| 126.6712 | 37.85352 | 1 | Paddy field | 0 | 1 | 0 | 0 | 0 | 0 | 1 |
| 127.0288 | 38.1089 | 1 | Paddy field | 0 | 1 | 0 | 0 | 0 | 0 | 1 |
| 124.2353 | 39.95572 | 1 | Paddy field | 0 | 1 | 0 | 0 | 0 | 0 | 1 |
| 124.452 | 39.901 | 1 | Paddy field | 0 | 1 | 0 | 0 | 0 | 0 | 1 |
| 125.221 | 38.8367 | 1 | Paddy field | 0 | 1 | 0 | 0 | 0 | 0 | 1 |
| 125.2297 | 38.85943 | 1 | Paddy field | 0 | 1 | 0 | 0 | 0 | 0 | 1 |
| 125.513 | 39.0741 | 1 | Paddy field | 0 | 1 | 0 | 0 | 0 | 0 | 1 |
| 125.513 | 39.5027 | 1 | Paddy field | 0 | 1 | 0 | 0 | 0 | 0 | 1 |
| 125.528 | 39.4522 | 1 | Paddy field | 0 | 1 | 0 | 0 | 0 | 0 | 1 |
| 125.605 | 39.19611 | 1 | Paddy field | 0 | 1 | 0 | 0 | 0 | 0 | 1 |
| 125.6777 | 39.11628 | 1 | Paddy field | 0 | 1 | 0 | 0 | 0 | 0 | 1 |
| 125.762 | 39.1522 | 1 | Paddy field | 0 | 1 | 0 | 0 | 0 | 0 | 1 |
| 126.316 | 36.6654 | 1 | Paddy field | 0 | 1 | 0 | 0 | 0 | 0 | 1 |
| 126.402 | 36.65504 | 1 | Paddy field | 0 | 1 | 0 | 0 | 0 | 0 | 1 |
| 126.4029 | 36.64047 | 1 | Paddy field | 0 | 1 | 0 | 0 | 0 | 0 | 1 |
| 126.4031 | 36.72806 | 1 | Paddy field | 0 | 1 | 0 | 0 | 0 | 0 | 1 |
| 126.426 | 36.69654 | 1 | Paddy field | 0 | 1 | 0 | 0 | 0 | 0 | 1 |
| 126.4377 | 36.77574 | 1 | Paddy field | 0 | 1 | 0 | 0 | 0 | 0 | 1 |
| 126.5323 | 36.63423 | 1 | Paddy field | 0 | 1 | 0 | 0 | 0 | 0 | 1 |
| 126.538 | 36.30268 | 1 | Paddy field | 0 | 1 | 0 | 0 | 0 | 0 | 1 |
| 126.559 | 36.39821 | 1 | Paddy field | 0 | 1 | 0 | 0 | 0 | 0 | 1 |
| 126.5621 | 36.77019 | 1 | Paddy field | 0 | 1 | 0 | 0 | 0 | 0 | 1 |
| 126.5739 | 36.34283 | 1 | Paddy field | 0 | 1 | 0 | 0 | 0 | 0 | 1 |
| 126.5897 | 36.63189 | 1 | Paddy field | 0 | 1 | 0 | 0 | 0 | 0 | 1 |
| 126.59 | 36.60223 | 1 | Paddy field | 0 | 1 | 0 | 0 | 0 | 0 | 1 |
| 126.5933 | 36.79476 | 1 | Paddy field | 0 | 1 | 0 | 0 | 0 | 0 | 1 |
| 126.7109 | 36.43696 | 1 | Paddy field | 0 | 1 | 0 | 0 | 0 | 0 | 1 |
| 126.7336 | 36.18311 | 1 | Paddy field | 0 | 1 | 0 | 0 | 0 | 0 | 1 |
| 126.8034 | 37.41084 | 1 | Paddy field | 0 | 1 | 0 | 0 | 0 | 0 | 1 |
| 126.8037 | 36.82759 | 1 | Paddy field | 0 | 1 | 0 | 0 | 0 | 0 | 1 |
| 126.8041 | 36.23931 | 1 | Paddy field | 0 | 1 | 0 | 0 | 0 | 0 | 1 |
| 126.8129 | 36.31947 | 1 | Paddy field | 0 | 1 | 0 | 0 | 0 | 0 | 1 |
| 126.8257 | 35.99329 | 1 | Paddy field | 0 | 1 | 0 | 0 | 0 | 0 | 1 |
| 126.8521 | 36.32932 | 1 | Paddy field | 0 | 1 | 0 | 0 | 0 | 0 | 1 |
| 126.862 | 35.9574 | 1 | Paddy field | 0 | 1 | 0 | 0 | 0 | 0 | 1 |
| 126.863 | 35.9222 | 1 | Paddy field | 0 | 1 | 0 | 0 | 0 | 0 | 1 |
| 126.8871 | 36.81949 | 1 | Paddy field | 0 | 1 | 0 | 0 | 0 | 0 | 1 |
| 126.8873 | 36.2128 | 1 | Paddy field | 0 | 1 | 0 | 0 | 0 | 0 | 1 |
| 126.9044 | 36.34584 | 1 | Paddy field | 0 | 1 | 0 | 0 | 0 | 0 | 1 |
| 126.908 | 36.6456 | 1 | Paddy field | 0 | 1 | 0 | 0 | 0 | 0 | 1 |
| 126.9129 | 36.21158 | 1 | Paddy field | 0 | 1 | 0 | 0 | 0 | 0 | 1 |
| 126.9193 | 36.96861 | 1 | Paddy field | 0 | 1 | 0 | 0 | 0 | 0 | 1 |
| 126.9344 | 35.97661 | 1 | Paddy field | 0 | 1 | 0 | 0 | 0 | 0 | 1 |
| 126.955 | 36.8111 | 1 | Paddy field | 0 | 1 | 0 | 0 | 0 | 0 | 1 |
| 126.9619 | 36.37362 | 1 | Paddy field | 0 | 1 | 0 | 0 | 0 | 0 | 1 |
| 126.964 | 36.1794 | 1 | Paddy field | 0 | 1 | 0 | 0 | 0 | 0 | 1 |
| 126.9694 | 36.32087 | 1 | Paddy field | 0 | 1 | 0 | 0 | 0 | 0 | 1 |
| 126.9696 | 36.82117 | 1 | Paddy field | 0 | 1 | 0 | 0 | 0 | 0 | 1 |
| 126.9925 | 36.88842 | 1 | Paddy field | 0 | 1 | 0 | 0 | 0 | 0 | 1 |
| 127.0444 | 35.94273 | 1 | Paddy field | 0 | 1 | 0 | 0 | 0 | 0 | 1 |
| 127.0572 | 36.03933 | 1 | Paddy field | 0 | 1 | 0 | 0 | 0 | 0 | 1 |
| 127.1976 | 36.01437 | 1 | Paddy field | 0 | 1 | 0 | 0 | 0 | 0 | 1 |
| 127.2075 | 36.53923 | 1 | Paddy field | 0 | 1 | 0 | 0 | 0 | 0 | 1 |
| 127.2426 | 36.58209 | 1 | Paddy field | 0 | 1 | 0 | 0 | 0 | 0 | 1 |
| 127.2522 | 36.53162 | 1 | Paddy field | 0 | 1 | 0 | 0 | 0 | 0 | 1 |
| 127.263 | 36.54067 | 1 | Paddy field | 0 | 1 | 0 | 0 | 0 | 0 | 1 |
| 127.2754 | 36.45004 | 1 | Paddy field | 0 | 1 | 0 | 0 | 0 | 0 | 1 |
| 127.276 | 36.5676 | 1 | Paddy field | 0 | 1 | 0 | 0 | 0 | 0 | 1 |
| 127.303 | 37.0662 | 1 | Paddy field | 0 | 1 | 0 | 0 | 0 | 0 | 1 |
| 127.3102 | 36.48295 | 1 | Paddy field | 0 | 1 | 0 | 0 | 0 | 0 | 1 |
| 127.3143 | 36.41053 | 1 | Paddy field | 0 | 1 | 0 | 0 | 0 | 0 | 1 |
| 127.3217 | 36.54598 | 1 | Paddy field | 0 | 1 | 0 | 0 | 0 | 0 | 1 |
| 127.322 | 36.4021 | 1 | Paddy field | 0 | 1 | 0 | 0 | 0 | 0 | 1 |
| 127.3281 | 36.50045 | 1 | Paddy field | 0 | 1 | 0 | 0 | 0 | 0 | 1 |
| 127.339 | 36.5423 | 1 | Paddy field | 0 | 1 | 0 | 0 | 0 | 0 | 1 |
| 127.3445 | 36.49212 | 1 | Paddy field | 0 | 1 | 0 | 0 | 0 | 0 | 1 |
| 127.3612 | 36.56839 | 1 | Paddy field | 0 | 1 | 0 | 0 | 0 | 0 | 1 |
| 127.3714 | 36.51063 | 1 | Paddy field | 0 | 1 | 0 | 0 | 0 | 0 | 1 |
| 127.4039 | 36.61882 | 1 | Paddy field | 0 | 1 | 0 | 0 | 0 | 0 | 1 |
| 127.5093 | 36.77853 | 1 | Paddy field | 0 | 1 | 0 | 0 | 0 | 0 | 1 |
| 128.4908 | 35.7376 | 1 | Paddy field | 0 | 1 | 0 | 0 | 0 | 0 | 1 |
| 126.6445 | 33.53625 | 0 | Paddy field | 0 | 1 | 0 | 0 | 0 | 0 | 1 |
| 127.9208 | 35.63977 | 0 | Paddy field | 0 | 1 | 0 | 0 | 0 | 0 | 1 |
| 127.9502 | 35.64069 | 0 | Paddy field | 0 | 1 | 0 | 0 | 0 | 0 | 1 |
| 128.1273 | 35.6557 | 0 | Paddy field | 0 | 1 | 0 | 0 | 0 | 0 | 1 |
| 128.0206 | 35.67944 | 0 | Paddy field | 0 | 1 | 0 | 0 | 0 | 0 | 1 |
| 127.9118 | 35.7896 | 0 | Paddy field | 0 | 1 | 0 | 0 | 0 | 0 | 1 |
| 126.9281 | 35.93056 | 0 | Paddy field | 0 | 1 | 0 | 0 | 0 | 0 | 1 |
| 126.9108 | 35.96361 | 0 | Paddy field | 0 | 1 | 0 | 0 | 0 | 0 | 1 |
| 126.8997 | 35.99028 | 0 | Paddy field | 0 | 1 | 0 | 0 | 0 | 0 | 1 |
| 127.1017 | 35.99028 | 0 | Paddy field | 0 | 1 | 0 | 0 | 0 | 0 | 1 |
| 126.7353 | 36.02972 | 0 | Paddy field | 0 | 1 | 0 | 0 | 0 | 0 | 1 |
| 126.7403 | 36.03278 | 0 | Paddy field | 0 | 1 | 0 | 0 | 0 | 0 | 1 |
| 126.9811 | 36.09444 | 0 | Paddy field | 0 | 1 | 0 | 0 | 0 | 0 | 1 |
| 128.3563 | 36.14812 | 0 | Paddy field | 0 | 1 | 0 | 0 | 0 | 0 | 1 |
| 128.203 | 36.17915 | 0 | Paddy field | 0 | 1 | 0 | 0 | 0 | 0 | 1 |
| 128.3503 | 36.18155 | 0 | Paddy field | 0 | 1 | 0 | 0 | 0 | 0 | 1 |
| 126.8802 | 36.21538 | 0 | Paddy field | 0 | 1 | 0 | 0 | 0 | 0 | 1 |
| 126.5522 | 36.21832 | 0 | Paddy field | 0 | 1 | 0 | 0 | 0 | 0 | 1 |
| 127.1496 | 36.21932 | 0 | Paddy field | 0 | 1 | 0 | 0 | 0 | 0 | 1 |
| 126.9317 | 36.22138 | 0 | Paddy field | 0 | 1 | 0 | 0 | 0 | 0 | 1 |
| 128.2409 | 36.22645 | 0 | Paddy field | 0 | 1 | 0 | 0 | 0 | 0 | 1 |
| 128.2971 | 36.22818 | 0 | Paddy field | 0 | 1 | 0 | 0 | 0 | 0 | 1 |
| 127.1575 | 36.22917 | 0 | Paddy field | 0 | 1 | 0 | 0 | 0 | 0 | 1 |
| 126.9504 | 36.23124 | 0 | Paddy field | 0 | 1 | 0 | 0 | 0 | 0 | 1 |
| 128.3419 | 36.24272 | 0 | Paddy field | 0 | 1 | 0 | 0 | 0 | 0 | 1 |
| 128.2434 | 36.24358 | 0 | Paddy field | 0 | 1 | 0 | 0 | 0 | 0 | 1 |
| 128.1006 | 36.29136 | 0 | Paddy field | 0 | 1 | 0 | 0 | 0 | 0 | 1 |
| 127.1023 | 36.31652 | 0 | Paddy field | 0 | 1 | 0 | 0 | 0 | 0 | 1 |
| 128.5418 | 36.31669 | 0 | Paddy field | 0 | 1 | 0 | 0 | 0 | 0 | 1 |
| 128.1189 | 36.32671 | 0 | Paddy field | 0 | 1 | 0 | 0 | 0 | 0 | 1 |
| 128.5554 | 36.33226 | 0 | Paddy field | 0 | 1 | 0 | 0 | 0 | 0 | 1 |
| 127.1456 | 36.33278 | 0 | Paddy field | 0 | 1 | 0 | 0 | 0 | 0 | 1 |
| 127.1289 | 36.33513 | 0 | Paddy field | 0 | 1 | 0 | 0 | 0 | 0 | 1 |
| 126.9227 | 36.33924 | 0 | Paddy field | 0 | 1 | 0 | 0 | 0 | 0 | 1 |
| 128.2567 | 36.33944 | 0 | Paddy field | 0 | 1 | 0 | 0 | 0 | 0 | 1 |
| 126.9931 | 36.34165 | 0 | Paddy field | 0 | 1 | 0 | 0 | 0 | 0 | 1 |
| 128.1341 | 36.35103 | 0 | Paddy field | 0 | 1 | 0 | 0 | 0 | 0 | 1 |
| 128.4422 | 36.3613 | 0 | Paddy field | 0 | 1 | 0 | 0 | 0 | 0 | 1 |
| 128.401 | 36.36909 | 0 | Paddy field | 0 | 1 | 0 | 0 | 0 | 0 | 1 |
| 128.3866 | 36.38577 | 0 | Paddy field | 0 | 1 | 0 | 0 | 0 | 0 | 1 |
| 128.2908 | 36.40627 | 0 | Paddy field | 0 | 1 | 0 | 0 | 0 | 0 | 1 |
| 128.3596 | 36.41099 | 0 | Paddy field | 0 | 1 | 0 | 0 | 0 | 0 | 1 |
| 128.3317 | 36.41742 | 0 | Paddy field | 0 | 1 | 0 | 0 | 0 | 0 | 1 |
| 128.3556 | 36.4186 | 0 | Paddy field | 0 | 1 | 0 | 0 | 0 | 0 | 1 |
| 126.4142 | 36.42622 | 0 | Paddy field | 0 | 1 | 0 | 0 | 0 | 0 | 1 |
| 126.4079 | 36.45347 | 0 | Paddy field | 0 | 1 | 0 | 0 | 0 | 0 | 1 |
| 128.1849 | 36.46186 | 0 | Paddy field | 0 | 1 | 0 | 0 | 0 | 0 | 1 |
| 128.2162 | 36.47049 | 0 | Paddy field | 0 | 1 | 0 | 0 | 0 | 0 | 1 |
| 128.1836 | 36.47194 | 0 | Paddy field | 0 | 1 | 0 | 0 | 0 | 0 | 1 |
| 128.1714 | 36.47358 | 0 | Paddy field | 0 | 1 | 0 | 0 | 0 | 0 | 1 |
| 127.7376 | 36.48518 | 0 | Paddy field | 0 | 1 | 0 | 0 | 0 | 0 | 1 |
| 128.287 | 36.48993 | 0 | Paddy field | 0 | 1 | 0 | 0 | 0 | 0 | 1 |
| 128.2924 | 36.49545 | 0 | Paddy field | 0 | 1 | 0 | 0 | 0 | 0 | 1 |
| 126.7777 | 36.55517 | 0 | Paddy field | 0 | 1 | 0 | 0 | 0 | 0 | 1 |
| 126.7711 | 36.62295 | 0 | Paddy field | 0 | 1 | 0 | 0 | 0 | 0 | 1 |
| 126.7343 | 36.64076 | 0 | Paddy field | 0 | 1 | 0 | 0 | 0 | 0 | 1 |
| 127.5134 | 36.64577 | 0 | Paddy field | 0 | 1 | 0 | 0 | 0 | 0 | 1 |
| 126.798 | 36.65489 | 0 | Paddy field | 0 | 1 | 0 | 0 | 0 | 0 | 1 |
| 126.6633 | 36.68022 | 0 | Paddy field | 0 | 1 | 0 | 0 | 0 | 0 | 1 |
| 127.4641 | 36.69094 | 0 | Paddy field | 0 | 1 | 0 | 0 | 0 | 0 | 1 |
| 126.5543 | 36.72088 | 0 | Paddy field | 0 | 1 | 0 | 0 | 0 | 0 | 1 |
| 127.4107 | 36.72799 | 0 | Paddy field | 0 | 1 | 0 | 0 | 0 | 0 | 1 |
| 127.4738 | 36.74163 | 0 | Paddy field | 0 | 1 | 0 | 0 | 0 | 0 | 1 |
| 126.7036 | 36.74885 | 0 | Paddy field | 0 | 1 | 0 | 0 | 0 | 0 | 1 |
| 126.7847 | 36.75111 | 0 | Paddy field | 0 | 1 | 0 | 0 | 0 | 0 | 1 |
| 126.8573 | 36.76025 | 0 | Paddy field | 0 | 1 | 0 | 0 | 0 | 0 | 1 |
| 127.1036 | 36.76694 | 0 | Paddy field | 0 | 1 | 0 | 0 | 0 | 0 | 1 |
| 126.8555 | 36.77472 | 0 | Paddy field | 0 | 1 | 0 | 0 | 0 | 0 | 1 |
| 126.8333 | 36.80444 | 0 | Paddy field | 0 | 1 | 0 | 0 | 0 | 0 | 1 |
| 126.9619 | 36.80883 | 0 | Paddy field | 0 | 1 | 0 | 0 | 0 | 0 | 1 |
| 126.8757 | 36.81982 | 0 | Paddy field | 0 | 1 | 0 | 0 | 0 | 0 | 1 |
| 126.9166 | 36.82535 | 0 | Paddy field | 0 | 1 | 0 | 0 | 0 | 0 | 1 |
| 127.4897 | 36.83848 | 0 | Paddy field | 0 | 1 | 0 | 0 | 0 | 0 | 1 |
| 126.8014 | 36.84083 | 0 | Paddy field | 0 | 1 | 0 | 0 | 0 | 0 | 1 |
| 127.5021 | 36.8473 | 0 | Paddy field | 0 | 1 | 0 | 0 | 0 | 0 | 1 |
| 126.9444 | 36.85752 | 0 | Paddy field | 0 | 1 | 0 | 0 | 0 | 0 | 1 |
| 127.4958 | 36.8602 | 0 | Paddy field | 0 | 1 | 0 | 0 | 0 | 0 | 1 |
| 128.8098 | 36.86796 | 0 | Paddy field | 0 | 1 | 0 | 0 | 0 | 0 | 1 |
| 127.8674 | 36.86928 | 0 | Paddy field | 0 | 1 | 0 | 0 | 0 | 0 | 1 |
| 127.4868 | 36.87525 | 0 | Paddy field | 0 | 1 | 0 | 0 | 0 | 0 | 1 |
| 127.9967 | 36.8814 | 0 | Paddy field | 0 | 1 | 0 | 0 | 0 | 0 | 1 |
| 127.8411 | 36.88334 | 0 | Paddy field | 0 | 1 | 0 | 0 | 0 | 0 | 1 |
| 127.847 | 36.88593 | 0 | Paddy field | 0 | 1 | 0 | 0 | 0 | 0 | 1 |
| 127.4635 | 36.8865 | 0 | Paddy field | 0 | 1 | 0 | 0 | 0 | 0 | 1 |
| 126.9456 | 36.88833 | 0 | Paddy field | 0 | 1 | 0 | 0 | 0 | 0 | 1 |
| 127.8219 | 36.89223 | 0 | Paddy field | 0 | 1 | 0 | 0 | 0 | 0 | 1 |
| 128.8832 | 36.8942 | 0 | Paddy field | 0 | 1 | 0 | 0 | 0 | 0 | 1 |
| 126.935 | 36.89944 | 0 | Paddy field | 0 | 1 | 0 | 0 | 0 | 0 | 1 |
| 126.9531 | 36.90722 | 0 | Paddy field | 0 | 1 | 0 | 0 | 0 | 0 | 1 |
| 126.9867 | 36.90861 | 0 | Paddy field | 0 | 1 | 0 | 0 | 0 | 0 | 1 |
| 128.1095 | 36.91571 | 0 | Paddy field | 0 | 1 | 0 | 0 | 0 | 0 | 1 |
| 127.5922 | 36.92947 | 0 | Paddy field | 0 | 1 | 0 | 0 | 0 | 0 | 1 |
| 127.4381 | 36.9517 | 0 | Paddy field | 0 | 1 | 0 | 0 | 0 | 0 | 1 |
| 127.4811 | 36.96174 | 0 | Paddy field | 0 | 1 | 0 | 0 | 0 | 0 | 1 |
| 127.7514 | 36.9725 | 0 | Paddy field | 0 | 1 | 0 | 0 | 0 | 0 | 1 |
| 127.4783 | 36.97696 | 0 | Paddy field | 0 | 1 | 0 | 0 | 0 | 0 | 1 |
| 127.4798 | 36.98147 | 0 | Paddy field | 0 | 1 | 0 | 0 | 0 | 0 | 1 |
| 127.0345 | 36.99019 | 0 | Paddy field | 0 | 1 | 0 | 0 | 0 | 0 | 1 |
| 127.2844 | 36.99222 | 0 | Paddy field | 0 | 1 | 0 | 0 | 0 | 0 | 1 |
| 127.4824 | 36.99379 | 0 | Paddy field | 0 | 1 | 0 | 0 | 0 | 0 | 1 |
| 126.9043 | 36.99774 | 0 | Paddy field | 0 | 1 | 0 | 0 | 0 | 0 | 1 |
| 127.8421 | 36.99904 | 0 | Paddy field | 0 | 1 | 0 | 0 | 0 | 0 | 1 |
| 126.9054 | 37.00341 | 0 | Paddy field | 0 | 1 | 0 | 0 | 0 | 0 | 1 |
| 127.5392 | 37.02981 | 0 | Paddy field | 0 | 1 | 0 | 0 | 0 | 0 | 1 |
| 127.153 | 37.03188 | 0 | Paddy field | 0 | 1 | 0 | 0 | 0 | 0 | 1 |
| 127.7519 | 37.05651 | 0 | Paddy field | 0 | 1 | 0 | 0 | 0 | 0 | 1 |
| 127.5332 | 37.05674 | 0 | Paddy field | 0 | 1 | 0 | 0 | 0 | 0 | 1 |
| 127.9093 | 37.06326 | 0 | Paddy field | 0 | 1 | 0 | 0 | 0 | 0 | 1 |
| 126.8186 | 37.06638 | 0 | Paddy field | 0 | 1 | 0 | 0 | 0 | 0 | 1 |
| 127.5396 | 37.07401 | 0 | Paddy field | 0 | 1 | 0 | 0 | 0 | 0 | 1 |
| 128.2778 | 37.07462 | 0 | Paddy field | 0 | 1 | 0 | 0 | 0 | 0 | 1 |
| 127.5386 | 37.08454 | 0 | Paddy field | 0 | 1 | 0 | 0 | 0 | 0 | 1 |
| 127.5313 | 37.09379 | 0 | Paddy field | 0 | 1 | 0 | 0 | 0 | 0 | 1 |
| 127.6241 | 37.09719 | 0 | Paddy field | 0 | 1 | 0 | 0 | 0 | 0 | 1 |
| 127.5799 | 37.09778 | 0 | Paddy field | 0 | 1 | 0 | 0 | 0 | 0 | 1 |
| 127.5743 | 37.09842 | 0 | Paddy field | 0 | 1 | 0 | 0 | 0 | 0 | 1 |
| 127.1421 | 37.09909 | 0 | Paddy field | 0 | 1 | 0 | 0 | 0 | 0 | 1 |
| 127.1493 | 37.10019 | 0 | Paddy field | 0 | 1 | 0 | 0 | 0 | 0 | 1 |
| 127.5249 | 37.10425 | 0 | Paddy field | 0 | 1 | 0 | 0 | 0 | 0 | 1 |
| 127.6562 | 37.10832 | 0 | Paddy field | 0 | 1 | 0 | 0 | 0 | 0 | 1 |
| 127.0132 | 37.10838 | 0 | Paddy field | 0 | 1 | 0 | 0 | 0 | 0 | 1 |
| 127.0081 | 37.11028 | 0 | Paddy field | 0 | 1 | 0 | 0 | 0 | 0 | 1 |
| 128.147 | 37.13436 | 0 | Paddy field | 0 | 1 | 0 | 0 | 0 | 0 | 1 |
| 127.6155 | 37.13872 | 0 | Paddy field | 0 | 1 | 0 | 0 | 0 | 0 | 1 |
| 127.6252 | 37.1395 | 0 | Paddy field | 0 | 1 | 0 | 0 | 0 | 0 | 1 |
| 126.8523 | 37.14553 | 0 | Paddy field | 0 | 1 | 0 | 0 | 0 | 0 | 1 |
| 127.6515 | 37.15663 | 0 | Paddy field | 0 | 1 | 0 | 0 | 0 | 0 | 1 |
| 128.2048 | 37.15687 | 0 | Paddy field | 0 | 1 | 0 | 0 | 0 | 0 | 1 |
| 127.4537 | 37.16091 | 0 | Paddy field | 0 | 1 | 0 | 0 | 0 | 0 | 1 |
| 127.7436 | 37.18639 | 0 | Paddy field | 0 | 1 | 0 | 0 | 0 | 0 | 1 |
| 127.6876 | 37.19129 | 0 | Paddy field | 0 | 1 | 0 | 0 | 0 | 0 | 1 |
| 127.7413 | 37.20001 | 0 | Paddy field | 0 | 1 | 0 | 0 | 0 | 0 | 1 |
| 127.7216 | 37.20455 | 0 | Paddy field | 0 | 1 | 0 | 0 | 0 | 0 | 1 |
| 127.4428 | 37.2184 | 0 | Paddy field | 0 | 1 | 0 | 0 | 0 | 0 | 1 |
| 126.8065 | 37.22093 | 0 | Paddy field | 0 | 1 | 0 | 0 | 0 | 0 | 1 |
| 127.7535 | 37.22873 | 0 | Paddy field | 0 | 1 | 0 | 0 | 0 | 0 | 1 |
| 127.7712 | 37.24852 | 0 | Paddy field | 0 | 1 | 0 | 0 | 0 | 0 | 1 |
| 127.6146 | 37.25192 | 0 | Paddy field | 0 | 1 | 0 | 0 | 0 | 0 | 1 |
| 127.7991 | 37.27061 | 0 | Paddy field | 0 | 1 | 0 | 0 | 0 | 0 | 1 |
| 127.7935 | 37.27483 | 0 | Paddy field | 0 | 1 | 0 | 0 | 0 | 0 | 1 |
| 127.8222 | 37.27863 | 0 | Paddy field | 0 | 1 | 0 | 0 | 0 | 0 | 1 |
| 127.5661 | 37.28942 | 0 | Paddy field | 0 | 1 | 0 | 0 | 0 | 0 | 1 |
| 127.5582 | 37.30131 | 0 | Paddy field | 0 | 1 | 0 | 0 | 0 | 0 | 1 |
| 127.5637 | 37.32789 | 0 | Paddy field | 0 | 1 | 0 | 0 | 0 | 0 | 1 |
| 127.8883 | 37.32826 | 0 | Paddy field | 0 | 1 | 0 | 0 | 0 | 0 | 1 |
| 127.548 | 37.34187 | 0 | Paddy field | 0 | 1 | 0 | 0 | 0 | 0 | 1 |
| 128.498 | 37.34392 | 0 | Paddy field | 0 | 1 | 0 | 0 | 0 | 0 | 1 |
| 127.8104 | 37.36714 | 0 | Paddy field | 0 | 1 | 0 | 0 | 0 | 0 | 1 |
| 127.6522 | 37.36925 | 0 | Paddy field | 0 | 1 | 0 | 0 | 0 | 0 | 1 |
| 126.8047 | 37.4075 | 0 | Paddy field | 0 | 1 | 0 | 0 | 0 | 0 | 1 |
| 126.4012 | 37.44999 | 0 | Paddy field | 0 | 1 | 0 | 0 | 0 | 0 | 1 |
| 129.0982 | 37.58177 | 0 | Paddy field | 0 | 1 | 0 | 0 | 0 | 0 | 1 |
| 127.1089 | 37.5847 | 0 | Paddy field | 0 | 1 | 0 | 0 | 0 | 0 | 1 |
| 126.5976 | 37.59147 | 0 | Paddy field | 0 | 1 | 0 | 0 | 0 | 0 | 1 |
| 126.6048 | 37.60471 | 0 | Paddy field | 0 | 1 | 0 | 0 | 0 | 0 | 1 |
| 126.5114 | 37.63833 | 0 | Paddy field | 0 | 1 | 0 | 0 | 0 | 0 | 1 |
| 126.4478 | 37.63981 | 0 | Paddy field | 0 | 1 | 0 | 0 | 0 | 0 | 1 |
| 126.4297 | 37.64111 | 0 | Paddy field | 0 | 1 | 0 | 0 | 0 | 0 | 1 |
| 126.4209 | 37.64408 | 0 | Paddy field | 0 | 1 | 0 | 0 | 0 | 0 | 1 |
| 126.2978 | 37.71512 | 0 | Paddy field | 0 | 1 | 0 | 0 | 0 | 0 | 1 |
| 126.3659 | 37.74004 | 0 | Paddy field | 0 | 1 | 0 | 0 | 0 | 0 | 1 |
| 126.6256 | 37.74195 | 0 | Paddy field | 0 | 1 | 0 | 0 | 0 | 0 | 1 |
| 126.3675 | 37.74695 | 0 | Paddy field | 0 | 1 | 0 | 0 | 0 | 0 | 1 |
| 126.7547 | 37.74891 | 0 | Paddy field | 0 | 1 | 0 | 0 | 0 | 0 | 1 |
| 126.3789 | 37.75299 | 0 | Paddy field | 0 | 1 | 0 | 0 | 0 | 0 | 1 |
| 126.7412 | 37.78537 | 0 | Paddy field | 0 | 1 | 0 | 0 | 0 | 0 | 1 |
| 126.7469 | 37.78881 | 0 | Paddy field | 0 | 1 | 0 | 0 | 0 | 0 | 1 |
| 126.82 | 37.79333 | 0 | Paddy field | 0 | 1 | 0 | 0 | 0 | 0 | 1 |
| 126.2742 | 37.80472 | 0 | Paddy field | 0 | 1 | 0 | 0 | 0 | 0 | 1 |
| 126.7128 | 37.81778 | 0 | Paddy field | 0 | 1 | 0 | 0 | 0 | 0 | 1 |
| 126.6164 | 37.88916 | 0 | Paddy field | 0 | 1 | 0 | 0 | 0 | 0 | 1 |
| 126.7548 | 37.89267 | 0 | Paddy field | 0 | 1 | 0 | 0 | 0 | 0 | 1 |
| 126.7582 | 37.89649 | 0 | Paddy field | 0 | 1 | 0 | 0 | 0 | 0 | 1 |
| 127.0047 | 38.03786 | 0 | Paddy field | 0 | 1 | 0 | 0 | 0 | 0 | 1 |
| 127.141 | 38.05431 | 0 | Paddy field | 0 | 1 | 0 | 0 | 0 | 0 | 1 |
| 127.0485 | 38.11639 | 0 | Paddy field | 0 | 1 | 0 | 0 | 0 | 0 | 1 |
| 127.1018 | 38.15839 | 0 | Paddy field | 0 | 1 | 0 | 0 | 0 | 0 | 1 |
| 127.2407 | 38.18949 | 0 | Paddy field | 0 | 1 | 0 | 0 | 0 | 0 | 1 |
| 127.2311 | 38.19831 | 0 | Paddy field | 0 | 1 | 0 | 0 | 0 | 0 | 1 |
| 127.2486 | 38.23191 | 0 | Paddy field | 0 | 1 | 0 | 0 | 0 | 0 | 1 |
| 127.2532 | 38.24481 | 0 | Paddy field | 0 | 1 | 0 | 0 | 0 | 0 | 1 |
| 127.2637 | 38.25454 | 0 | Paddy field | 0 | 1 | 0 | 0 | 0 | 0 | 1 |
| 127.3532 | 38.26328 | 0 | Paddy field | 0 | 1 | 0 | 0 | 0 | 0 | 1 |
| 128.1433 | 38.28889 | 0 | Paddy field | 0 | 1 | 0 | 0 | 0 | 0 | 1 |
| 125.3193 | 38.84485 | 0 | Paddy field | 0 | 1 | 0 | 0 | 0 | 0 | 1 |
| 125.2142 | 38.89851 | 0 | Paddy field | 0 | 1 | 0 | 0 | 0 | 0 | 1 |
| 125.3763 | 38.96651 | 0 | Paddy field | 0 | 1 | 0 | 0 | 0 | 0 | 1 |
| 125.2182 | 38.98765 | 0 | Paddy field | 0 | 1 | 0 | 0 | 0 | 0 | 1 |
| 125.7069 | 39.0778 | 0 | Paddy field | 0 | 1 | 0 | 0 | 0 | 0 | 1 |
| 125.6829 | 39.12811 | 0 | Paddy field | 0 | 1 | 0 | 0 | 0 | 0 | 1 |
| 125.6343 | 39.17161 | 0 | Paddy field | 0 | 1 | 0 | 0 | 0 | 0 | 1 |
| 124.2111 | 39.80164 | 0 | Paddy field | 0 | 1 | 0 | 0 | 0 | 0 | 1 |
| 124.2563 | 39.84208 | 0 | Paddy field | 0 | 1 | 0 | 0 | 0 | 0 | 1 |
| 123.5775 | 39.87672 | 0 | Paddy field | 0 | 1 | 0 | 0 | 0 | 0 | 1 |
| 124.3741 | 39.88905 | 0 | Paddy field | 0 | 1 | 0 | 0 | 0 | 0 | 1 |
| 124.2373 | 39.92901 | 0 | Paddy field | 0 | 1 | 0 | 0 | 0 | 0 | 1 |
| 124.3956 | 39.96665 | 0 | Paddy field | 0 | 1 | 0 | 0 | 0 | 0 | 1 |
| 130.5794 | 42.355 | 0 | Paddy field | 0 | 1 | 0 | 0 | 0 | 0 | 1 |
| 126.5504 | 36.63942 | 1 | Shrub | 0 | 0 | 1 | 0 | 0 | 0 | 0 |
| 126.6515 | 37.5682 | 1 | Shrub | 0 | 0 | 1 | 0 | 0 | 0 | 0 |
| 126.989 | 36.8733 | 1 | Shrub | 0 | 0 | 1 | 0 | 0 | 0 | 0 |
| 127.0169 | 38.07928 | 1 | Shrub | 0 | 0 | 1 | 0 | 0 | 0 | 0 |
| 127.3178 | 36.51285 | 1 | Shrub | 0 | 0 | 1 | 0 | 0 | 0 | 0 |
| 128.3771 | 36.17787 | 0 | Shrub | 0 | 0 | 1 | 0 | 0 | 0 | 0 |
| 126.4149 | 36.41405 | 0 | Shrub | 0 | 0 | 1 | 0 | 0 | 0 | 0 |
| 127.8004 | 36.90055 | 0 | Shrub | 0 | 0 | 1 | 0 | 0 | 0 | 0 |
| 127.4406 | 36.92221 | 0 | Shrub | 0 | 0 | 1 | 0 | 0 | 0 | 0 |
| 127.8308 | 36.96987 | 0 | Shrub | 0 | 0 | 1 | 0 | 0 | 0 | 0 |
| 127.8861 | 37.00395 | 0 | Shrub | 0 | 0 | 1 | 0 | 0 | 0 | 0 |
| 127.4364 | 37.00864 | 0 | Shrub | 0 | 0 | 1 | 0 | 0 | 0 | 0 |
| 126.7884 | 37.09275 | 0 | Shrub | 0 | 0 | 1 | 0 | 0 | 0 | 0 |
| 128.4211 | 37.22284 | 0 | Shrub | 0 | 0 | 1 | 0 | 0 | 0 | 0 |
| 128.4603 | 37.25837 | 0 | Shrub | 0 | 0 | 1 | 0 | 0 | 0 | 0 |
| 126.5225 | 37.64778 | 0 | Shrub | 0 | 0 | 1 | 0 | 0 | 0 | 0 |
| 126.2958 | 37.70556 | 0 | Shrub | 0 | 0 | 1 | 0 | 0 | 0 | 0 |
| 126.5319 | 37.75008 | 0 | Shrub | 0 | 0 | 1 | 0 | 0 | 0 | 0 |
| 128.3289 | 37.94537 | 0 | Shrub | 0 | 0 | 1 | 0 | 0 | 0 | 0 |
| 124.6433 | 37.94917 | 0 | Shrub | 0 | 0 | 1 | 0 | 0 | 0 | 0 |
| 124.6497 | 37.96869 | 0 | Shrub | 0 | 0 | 1 | 0 | 0 | 0 | 0 |
| 127.0603 | 38.04583 | 0 | Shrub | 0 | 0 | 1 | 0 | 0 | 0 | 0 |
| 127.5101 | 38.05662 | 0 | Shrub | 0 | 0 | 1 | 0 | 0 | 0 | 0 |
| 127.0655 | 38.05681 | 0 | Shrub | 0 | 0 | 1 | 0 | 0 | 0 | 0 |
| 127.0181 | 38.06556 | 0 | Shrub | 0 | 0 | 1 | 0 | 0 | 0 | 0 |
| 127.0174 | 38.08095 | 0 | Shrub | 0 | 0 | 1 | 0 | 0 | 0 | 0 |
| 127.1132 | 38.19302 | 0 | Shrub | 0 | 0 | 1 | 0 | 0 | 0 | 0 |
| 127.2237 | 38.24432 | 0 | Shrub | 0 | 0 | 1 | 0 | 0 | 0 | 0 |
| 124.3341 | 39.89104 | 0 | Shrub | 0 | 0 | 1 | 0 | 0 | 0 | 0 |
| 130.5138 | 42.33822 | 0 | Shrub | 0 | 0 | 1 | 0 | 0 | 0 | 0 |
| 126.5877 | 37.00169 | 1 | Sparse vegetation | 0 | 0 | 0 | 0 | 0 | 0 | 0 |
| 126.6602 | 37.18018 | 0 | Sparse vegetation | 0 | 0 | 0 | 0 | 0 | 0 | 0 |
| 126.5434 | 37.51882 | 0 | Sparse vegetation | 0 | 0 | 0 | 0 | 0 | 0 | 0 |
| 127.1211 | 35.81155 | 1 | Urban | 0 | 0 | 0 | 0 | 1 | 0 | 0 |
| 126.8041 | 37.4436 | 1 | Urban | 0 | 0 | 0 | 0 | 1 | 0 | 0 |
| 126.6096 | 37.45734 | 1 | Urban | 0 | 0 | 0 | 0 | 1 | 0 | 0 |
| 126.8051 | 37.46247 | 1 | Urban | 0 | 0 | 0 | 0 | 1 | 0 | 0 |
| 126.8566 | 37.49967 | 1 | Urban | 0 | 0 | 0 | 0 | 1 | 0 | 0 |
| 126.7217 | 37.52809 | 1 | Urban | 0 | 0 | 0 | 0 | 1 | 0 | 0 |
| 126.8758 | 37.52942 | 1 | Urban | 0 | 0 | 0 | 0 | 1 | 0 | 0 |
| 126.6397 | 37.55594 | 1 | Urban | 0 | 0 | 0 | 0 | 1 | 0 | 0 |
| 126.7402 | 37.5589 | 1 | Urban | 0 | 0 | 0 | 0 | 1 | 0 | 0 |
| 126.9095 | 37.55917 | 1 | Urban | 0 | 0 | 0 | 0 | 1 | 0 | 0 |
| 126.8306 | 37.5708 | 1 | Urban | 0 | 0 | 0 | 0 | 1 | 0 | 0 |
| 126.8603 | 37.57284 | 1 | Urban | 0 | 0 | 0 | 0 | 1 | 0 | 0 |
| 126.6956 | 37.64513 | 1 | Urban | 0 | 0 | 0 | 0 | 1 | 0 | 0 |
| 126.7869 | 37.88312 | 1 | Urban | 0 | 0 | 0 | 0 | 1 | 0 | 0 |
| 125.7586 | 39.06334 | 1 | Urban | 0 | 0 | 0 | 0 | 1 | 0 | 0 |
| 125.7716 | 39.00839 | 1 | Urban | 0 | 0 | 0 | 0 | 1 | 0 | 0 |
| 126.4592 | 36.76236 | 1 | Urban | 0 | 0 | 0 | 0 | 1 | 0 | 0 |
| 126.6121 | 37.4721 | 1 | Urban | 0 | 0 | 0 | 0 | 1 | 0 | 0 |
| 126.6746 | 37.50219 | 1 | Urban | 0 | 0 | 0 | 0 | 1 | 0 | 0 |
| 126.6774 | 37.53782 | 1 | Urban | 0 | 0 | 0 | 0 | 1 | 0 | 0 |
| 126.6869 | 37.46735 | 1 | Urban | 0 | 0 | 0 | 0 | 1 | 0 | 0 |
| 126.7059 | 37.42545 | 1 | Urban | 0 | 0 | 0 | 0 | 1 | 0 | 0 |
| 126.7153 | 37.53922 | 1 | Urban | 0 | 0 | 0 | 0 | 1 | 0 | 0 |
| 126.7262 | 37.65944 | 1 | Urban | 0 | 0 | 0 | 0 | 1 | 0 | 0 |
| 126.7277 | 37.53611 | 1 | Urban | 0 | 0 | 0 | 0 | 1 | 0 | 0 |
| 126.7379 | 37.61615 | 1 | Urban | 0 | 0 | 0 | 0 | 1 | 0 | 0 |
| 126.7392 | 37.4461 | 1 | Urban | 0 | 0 | 0 | 0 | 1 | 0 | 0 |
| 126.7406 | 37.69673 | 1 | Urban | 0 | 0 | 0 | 0 | 1 | 0 | 0 |
| 126.7613 | 37.48151 | 1 | Urban | 0 | 0 | 0 | 0 | 1 | 0 | 0 |
| 126.7673 | 37.52249 | 1 | Urban | 0 | 0 | 0 | 0 | 1 | 0 | 0 |
| 126.8369 | 37.41147 | 1 | Urban | 0 | 0 | 0 | 0 | 1 | 0 | 0 |
| 126.8415 | 37.64291 | 1 | Urban | 0 | 0 | 0 | 0 | 1 | 0 | 0 |
| 126.8725 | 37.52066 | 1 | Urban | 0 | 0 | 0 | 0 | 1 | 0 | 0 |
| 126.8847 | 37.29204 | 1 | Urban | 0 | 0 | 0 | 0 | 1 | 0 | 0 |
| 126.892 | 37.4915 | 1 | Urban | 0 | 0 | 0 | 0 | 1 | 0 | 0 |
| 126.8994 | 37.53599 | 1 | Urban | 0 | 0 | 0 | 0 | 1 | 0 | 0 |
| 126.9068 | 37.49108 | 1 | Urban | 0 | 0 | 0 | 0 | 1 | 0 | 0 |
| 126.9244 | 37.47463 | 1 | Urban | 0 | 0 | 0 | 0 | 1 | 0 | 0 |
| 126.9478 | 37.57154 | 1 | Urban | 0 | 0 | 0 | 0 | 1 | 0 | 0 |
| 126.954 | 37.5417 | 1 | Urban | 0 | 0 | 0 | 0 | 1 | 0 | 0 |
| 126.9651 | 37.48821 | 1 | Urban | 0 | 0 | 0 | 0 | 1 | 0 | 0 |
| 126.971 | 37.47904 | 1 | Urban | 0 | 0 | 0 | 0 | 1 | 0 | 0 |
| 126.977 | 37.4094 | 1 | Urban | 0 | 0 | 0 | 0 | 1 | 0 | 0 |
| 126.994 | 37.5887 | 1 | Urban | 0 | 0 | 0 | 0 | 1 | 0 | 0 |
| 126.9963 | 37.18717 | 1 | Urban | 0 | 0 | 0 | 0 | 1 | 0 | 0 |
| 126.998 | 37.54692 | 1 | Urban | 0 | 0 | 0 | 0 | 1 | 0 | 0 |
| 126.998 | 36.79815 | 1 | Urban | 0 | 0 | 0 | 0 | 1 | 0 | 0 |
| 127.1123 | 35.83262 | 1 | Urban | 0 | 0 | 0 | 0 | 1 | 0 | 0 |
| 127.1192 | 37.10764 | 1 | Urban | 0 | 0 | 0 | 0 | 1 | 0 | 0 |
| 127.1473 | 35.85099 | 1 | Urban | 0 | 0 | 0 | 0 | 1 | 0 | 0 |
| 127.298 | 36.5901 | 1 | Urban | 0 | 0 | 0 | 0 | 1 | 0 | 0 |
| 127.3024 | 36.40579 | 1 | Urban | 0 | 0 | 0 | 0 | 1 | 0 | 0 |
| 127.3379 | 36.40157 | 1 | Urban | 0 | 0 | 0 | 0 | 1 | 0 | 0 |
| 126.8872 | 35.23361 | 0 | Urban | 0 | 0 | 0 | 0 | 1 | 0 | 0 |
| 128.7921 | 35.33743 | 0 | Urban | 0 | 0 | 0 | 0 | 1 | 0 | 0 |
| 126.9858 | 35.93972 | 0 | Urban | 0 | 0 | 0 | 0 | 1 | 0 | 0 |
| 126.745 | 35.95997 | 0 | Urban | 0 | 0 | 0 | 0 | 1 | 0 | 0 |
| 128.3493 | 36.13595 | 0 | Urban | 0 | 0 | 0 | 0 | 1 | 0 | 0 |
| 128.3244 | 36.13831 | 0 | Urban | 0 | 0 | 0 | 0 | 1 | 0 | 0 |
| 128.3636 | 36.14836 | 0 | Urban | 0 | 0 | 0 | 0 | 1 | 0 | 0 |
| 128.1721 | 36.41058 | 0 | Urban | 0 | 0 | 0 | 0 | 1 | 0 | 0 |
| 126.8446 | 36.70156 | 0 | Urban | 0 | 0 | 0 | 0 | 1 | 0 | 0 |
| 127.4407 | 36.86586 | 0 | Urban | 0 | 0 | 0 | 0 | 1 | 0 | 0 |
| 127.9023 | 36.9558 | 0 | Urban | 0 | 0 | 0 | 0 | 1 | 0 | 0 |
| 127.9228 | 36.96253 | 0 | Urban | 0 | 0 | 0 | 0 | 1 | 0 | 0 |
| 127.9114 | 36.96514 | 0 | Urban | 0 | 0 | 0 | 0 | 1 | 0 | 0 |
| 127.9062 | 36.9735 | 0 | Urban | 0 | 0 | 0 | 0 | 1 | 0 | 0 |
| 127.0799 | 36.97364 | 0 | Urban | 0 | 0 | 0 | 0 | 1 | 0 | 0 |
| 127.9092 | 36.978 | 0 | Urban | 0 | 0 | 0 | 0 | 1 | 0 | 0 |
| 127.5936 | 36.98576 | 0 | Urban | 0 | 0 | 0 | 0 | 1 | 0 | 0 |
| 127.2586 | 37.015 | 0 | Urban | 0 | 0 | 0 | 0 | 1 | 0 | 0 |
| 127.0383 | 37.06428 | 0 | Urban | 0 | 0 | 0 | 0 | 1 | 0 | 0 |
| 127.6299 | 37.11017 | 0 | Urban | 0 | 0 | 0 | 0 | 1 | 0 | 0 |
| 127.6232 | 37.11874 | 0 | Urban | 0 | 0 | 0 | 0 | 1 | 0 | 0 |
| 128.2091 | 37.15309 | 0 | Urban | 0 | 0 | 0 | 0 | 1 | 0 | 0 |
| 126.9842 | 37.23911 | 0 | Urban | 0 | 0 | 0 | 0 | 1 | 0 | 0 |
| 126.989 | 37.26709 | 0 | Urban | 0 | 0 | 0 | 0 | 1 | 0 | 0 |
| 126.8008 | 37.305 | 0 | Urban | 0 | 0 | 0 | 0 | 1 | 0 | 0 |
| 126.8017 | 37.39861 | 0 | Urban | 0 | 0 | 0 | 0 | 1 | 0 | 0 |
| 126.6787 | 37.40951 | 0 | Urban | 0 | 0 | 0 | 0 | 1 | 0 | 0 |
| 126.9086 | 37.41299 | 0 | Urban | 0 | 0 | 0 | 0 | 1 | 0 | 0 |
| 126.8094 | 37.41333 | 0 | Urban | 0 | 0 | 0 | 0 | 1 | 0 | 0 |
| 126.795 | 37.42028 | 0 | Urban | 0 | 0 | 0 | 0 | 1 | 0 | 0 |
| 126.8933 | 37.43023 | 0 | Urban | 0 | 0 | 0 | 0 | 1 | 0 | 0 |
| 126.9938 | 37.43374 | 0 | Urban | 0 | 0 | 0 | 0 | 1 | 0 | 0 |
| 126.6241 | 37.43633 | 0 | Urban | 0 | 0 | 0 | 0 | 1 | 0 | 0 |
| 126.6855 | 37.4499 | 0 | Urban | 0 | 0 | 0 | 0 | 1 | 0 | 0 |
| 126.8779 | 37.45689 | 0 | Urban | 0 | 0 | 0 | 0 | 1 | 0 | 0 |
| 126.9518 | 37.45824 | 0 | Urban | 0 | 0 | 0 | 0 | 1 | 0 | 0 |
| 126.7236 | 37.45889 | 0 | Urban | 0 | 0 | 0 | 0 | 1 | 0 | 0 |
| 126.6404 | 37.45919 | 0 | Urban | 0 | 0 | 0 | 0 | 1 | 0 | 0 |
| 126.7234 | 37.48716 | 0 | Urban | 0 | 0 | 0 | 0 | 1 | 0 | 0 |
| 126.8743 | 37.49773 | 0 | Urban | 0 | 0 | 0 | 0 | 1 | 0 | 0 |
| 126.9648 | 37.51898 | 0 | Urban | 0 | 0 | 0 | 0 | 1 | 0 | 0 |
| 126.8309 | 37.52887 | 0 | Urban | 0 | 0 | 0 | 0 | 1 | 0 | 0 |
| 127.0065 | 37.53498 | 0 | Urban | 0 | 0 | 0 | 0 | 1 | 0 | 0 |
| 126.7452 | 37.5441 | 0 | Urban | 0 | 0 | 0 | 0 | 1 | 0 | 0 |
| 126.684 | 37.55763 | 0 | Urban | 0 | 0 | 0 | 0 | 1 | 0 | 0 |
| 126.9472 | 37.5618 | 0 | Urban | 0 | 0 | 0 | 0 | 1 | 0 | 0 |
| 126.8058 | 37.56222 | 0 | Urban | 0 | 0 | 0 | 0 | 1 | 0 | 0 |
| 126.8003 | 37.56528 | 0 | Urban | 0 | 0 | 0 | 0 | 1 | 0 | 0 |
| 126.8061 | 37.5725 | 0 | Urban | 0 | 0 | 0 | 0 | 1 | 0 | 0 |
| 126.9533 | 37.5812 | 0 | Urban | 0 | 0 | 0 | 0 | 1 | 0 | 0 |
| 127.1308 | 37.58726 | 0 | Urban | 0 | 0 | 0 | 0 | 1 | 0 | 0 |
| 126.907 | 37.58943 | 0 | Urban | 0 | 0 | 0 | 0 | 1 | 0 | 0 |
| 126.6117 | 37.64228 | 0 | Urban | 0 | 0 | 0 | 0 | 1 | 0 | 0 |
| 126.6992 | 37.64303 | 0 | Urban | 0 | 0 | 0 | 0 | 1 | 0 | 0 |
| 126.61 | 37.65061 | 0 | Urban | 0 | 0 | 0 | 0 | 1 | 0 | 0 |
| 126.6131 | 37.65754 | 0 | Urban | 0 | 0 | 0 | 0 | 1 | 0 | 0 |
| 126.6522 | 37.66297 | 0 | Urban | 0 | 0 | 0 | 0 | 1 | 0 | 0 |
| 126.7575 | 37.66371 | 0 | Urban | 0 | 0 | 0 | 0 | 1 | 0 | 0 |
| 126.7071 | 37.66947 | 0 | Urban | 0 | 0 | 0 | 0 | 1 | 0 | 0 |
| 126.7163 | 37.67308 | 0 | Urban | 0 | 0 | 0 | 0 | 1 | 0 | 0 |
| 126.6971 | 37.67388 | 0 | Urban | 0 | 0 | 0 | 0 | 1 | 0 | 0 |
| 126.7281 | 37.67576 | 0 | Urban | 0 | 0 | 0 | 0 | 1 | 0 | 0 |
| 126.7161 | 37.67809 | 0 | Urban | 0 | 0 | 0 | 0 | 1 | 0 | 0 |
| 126.7244 | 37.6796 | 0 | Urban | 0 | 0 | 0 | 0 | 1 | 0 | 0 |
| 126.6932 | 37.68001 | 0 | Urban | 0 | 0 | 0 | 0 | 1 | 0 | 0 |
| 126.7032 | 37.68047 | 0 | Urban | 0 | 0 | 0 | 0 | 1 | 0 | 0 |
| 126.7004 | 37.6845 | 0 | Urban | 0 | 0 | 0 | 0 | 1 | 0 | 0 |
| 126.7395 | 37.69184 | 0 | Urban | 0 | 0 | 0 | 0 | 1 | 0 | 0 |
| 126.7607 | 37.70424 | 0 | Urban | 0 | 0 | 0 | 0 | 1 | 0 | 0 |
| 126.6982 | 37.71997 | 0 | Urban | 0 | 0 | 0 | 0 | 1 | 0 | 0 |
| 126.7186 | 37.72915 | 0 | Urban | 0 | 0 | 0 | 0 | 1 | 0 | 0 |
| 126.767 | 37.74774 | 0 | Urban | 0 | 0 | 0 | 0 | 1 | 0 | 0 |
| 126.7918 | 37.75572 | 0 | Urban | 0 | 0 | 0 | 0 | 1 | 0 | 0 |
| 127.7292 | 37.87457 | 0 | Urban | 0 | 0 | 0 | 0 | 1 | 0 | 0 |
| 123.5121 | 41.84869 | 0 | Urban | 0 | 0 | 0 | 0 | 1 | 0 | 0 |
| 123.3804 | 41.85733 | 0 | Urban | 0 | 0 | 0 | 0 | 1 | 0 | 0 |
| 126.9218 | 36.9148 | 1 | Water bodies | 0 | 0 | 0 | 0 | 0 | 1 | 0 |
| 126.2217 | 37.65264 | 1 | Water bodies | 0 | 0 | 0 | 0 | 0 | 1 | 0 |
| 126.5355 | 37.77511 | 1 | Water bodies | 0 | 0 | 0 | 0 | 0 | 1 | 0 |
| 126.476 | 36.609 | 1 | Water bodies | 0 | 0 | 0 | 0 | 0 | 1 | 0 |
| 126.5652 | 37.76496 | 1 | Water bodies | 0 | 0 | 0 | 0 | 0 | 1 | 0 |
| 126.8413 | 36.84637 | 1 | Water bodies | 0 | 0 | 0 | 0 | 0 | 1 | 0 |
| 126.8414 | 36.82297 | 1 | Water bodies | 0 | 0 | 0 | 0 | 0 | 1 | 0 |
| 126.8579 | 36.86608 | 1 | Water bodies | 0 | 0 | 0 | 0 | 0 | 1 | 0 |
| 126.9766 | 36.95796 | 1 | Water bodies | 0 | 0 | 0 | 0 | 0 | 1 | 0 |
| 128.9622 | 35.13796 | 0 | Water bodies | 0 | 0 | 0 | 0 | 0 | 1 | 0 |
| 126.8775 | 36.08821 | 0 | Water bodies | 0 | 0 | 0 | 0 | 0 | 1 | 0 |
| 126.8301 | 36.85265 | 0 | Water bodies | 0 | 0 | 0 | 0 | 0 | 1 | 0 |
| 126.9603 | 36.91333 | 0 | Water bodies | 0 | 0 | 0 | 0 | 0 | 1 | 0 |
| 126.6104 | 37.00589 | 0 | Water bodies | 0 | 0 | 0 | 0 | 0 | 1 | 0 |
| 126.8528 | 37.57034 | 0 | Water bodies | 0 | 0 | 0 | 0 | 0 | 1 | 0 |
| 126.7947 | 37.60663 | 0 | Water bodies | 0 | 0 | 0 | 0 | 0 | 1 | 0 |
| 126.6894 | 37.66305 | 0 | Water bodies | 0 | 0 | 0 | 0 | 0 | 1 | 0 |
| 126.2042 | 37.6673 | 0 | Water bodies | 0 | 0 | 0 | 0 | 0 | 1 | 0 |
| 126.6648 | 37.70209 | 0 | Water bodies | 0 | 0 | 0 | 0 | 0 | 1 | 0 |
| 126.686 | 37.73024 | 0 | Water bodies | 0 | 0 | 0 | 0 | 0 | 1 | 0 |
| 126.5283 | 37.7677 | 0 | Water bodies | 0 | 0 | 0 | 0 | 0 | 1 | 0 |
| 126.5475 | 37.77258 | 0 | Water bodies | 0 | 0 | 0 | 0 | 0 | 1 | 0 |
| 126.6266 | 37.78224 | 0 | Water bodies | 0 | 0 | 0 | 0 | 0 | 1 | 0 |
| 126.6098 | 37.78262 | 0 | Water bodies | 0 | 0 | 0 | 0 | 0 | 1 | 0 |
| 126.6872 | 37.84053 | 0 | Water bodies | 0 | 0 | 0 | 0 | 0 | 1 | 0 |
| 127.7082 | 37.88421 | 0 | Water bodies | 0 | 0 | 0 | 0 | 0 | 1 | 0 |
| 124.2939 | 39.84526 | 0 | Water bodies | 0 | 0 | 0 | 0 | 0 | 1 | 0 |
| 130.4011 | 42.26152 | 0 | Water bodies | 0 | 0 | 0 | 0 | 0 | 1 | 0 |
| 130.5843 | 42.32588 | 0 | Water bodies | 0 | 0 | 0 | 0 | 0 | 1 | 0 |
| 130.5774 | 42.3318 | 0 | Water bodies | 0 | 0 | 0 | 0 | 0 | 1 | 0 |
