## Supplementary File 2 for "Korean endemic species no more: on the occurrence of *Pelophylax chosenicus* in China"

Supplementary File 2. Information on molecular samples used to reconstruct the phylogenetic trees in this study (*n* taxa = 34; 32 ingroup taxa and 2 outgroup taxa - *Pelophylax bedrigae* and *Pelophylax kurtmuelleri*).

| Taxa | Voucher ID/  reference ID in tree | Country: localities | 16S rRNA accession number | Reference |
| --- | --- | --- | --- | --- |
| Pelophylax *chosenicus* | 17PcDD001 | China: Dandong | ON054286 | This study |
| Pelophylax *chosenicus* | 17PcDD002 | China: Dandong | ON054287 | This study |
| Pelophylax *chosenicus* | 17PcDD003 | China: Dandong | ON054288 | This study |
| Pelophylax *chosenicus* | 17PcDD004 | China: Dandong | ON054289 | This study |
| Pelophylax *chosenicus* | 17PcDD005 | China: Dandong | ON054290 | This study |
| *Pelophylax chosenicus* | Pc_MMS523_Sk | Republic of Korea | EU386941 | Direct submission by Min et al. (2016) |
| *Pelophylax chosenicus* | Pc_MMS513_Sk | Republic of Korea | EU387004 | Direct submission by Min et al. (2016) |
| *Pelophylax chosenicus* | Pc_MMS176_Sk | Republic of Korea | EU386945 | Direct submission by Min et al. (2016) |
| *Pelophylax chosenicus* | Pc_MMS179_Sk | Republic of Korea | EU386932 | Direct submission by Min et al. (2016) |
| *Pelophylax chosenicus* | Pc_MMS533_Sk | Republic of Korea | EU386947 | Direct submission by Min et al. (2016) |
| *Pelophylax chosenicus* | Pc_MMS432_Sk | Republic of Korea | EU386948 | Direct submission by Min et al. (2016) |
| *Pelophylax chosenicus* | Pc_MMS431_Sk | Republic of Korea | EU386935 | Direct submission by Min et al. (2016) |
| *Pelophylax chosenicus* | Pc_MMS454_Sk | Republic of Korea | EU386949 | Direct submission by Min et al. (2016) |
| *Pelophylax chosenicus* | Pc_MMS428_Sk | Republic of Korea | EU386950 | Direct submission by Min et al. (2016) |
| *Pelophylax chosenicus* | Pc_MMS440_Sk | Republic of Korea | EU386857 | Direct submission by Min et al. (2016) |
| *Pelophylax chosenicus* | Pc_MMS446_Sk | Republic of Korea | EU386958 | Direct submission by Min et al. (2016) |
| *Pelophylax chosenicus* | Pc_MMS524_Sk | Republic of Korea | EU386959 | Direct submission by Min et al. (2016) |
| *Pelophylax chosenicus* | Pc_MMS531_Sk | Republic of Korea | EU386943 | Direct submission by Min et al. (2016) |
| *Pelophylax chosenicus* | Pc_MMS189_Sk | Republic of Korea | EU386944 | Direct submission by Min et al. (2016) |
| *Pelophylax chosenicus* | Pc_MMS171_Sk | Republic of Korea | EU386946 | Direct submission by Min et al. (2016) |
| *Pelophylax chosenicus* | Pc_MMS102_Sk | Republic of Korea | EU386914 | Direct submission by Min et al. (2016) |
| *Pelophylax chosenicus* | Pc_MMS510_Sk | Republic of Korea | EU386908 | Direct submission by Min et al. (2016) |
| *Pelophylax chosenicus* | Pc_fullmtDNA | Republic of Korea | JF730436 | Ryu and Hwang (2011) |
| *Pelophylax plancyi* | Pp_MMS301_Ch | Republic of Korea | EU386960 | Direct submission by Min et al. (2016) |
| *Pelophylax plancyi* | Pp_NIBRAM0000000038_Sk | Chungcheongbuk-do Goesan-gun | JQ815307 | Jeong et al. (2013) |
| *Pelophylax plancyi* | Pp_fullmtDNA | China | EF196679 | Direct submission by Nie et al. (2016) |
| *Pelophylax nigromaculatus* | Pn_Pn3_J | Aichi: Japan | LC389203 | Tokumoto et al. (2019) |
| *Pelophylax nigromaculatus* | Pn_Pn5_J | Aichi: Japan | LC389205 | Tokumoto et al. (2019) |
| *Pelophylax nigromaculatus* | Pn_fullmtDNA | China | KT878718 | Jiang et al. (2017) |
| *Pelophylax nigromaculatus* | Pn_FMNH 232879_C | China: Sichuan | DQ283137 | Frost et al. (2006) |
| *Pelophylax nigromaculatus* | Pn_SCUM045199CJ | China | KX269216 | Yuan et al. (2016) |
| *Pelophylax hubeiensis* | Phub_C1 | China: Anhui | AF315137 | Jiang and Zhao (2005) |
| *Pelophylax kurtmuelleri* | *Pelophylax kurtmuelleri* | Greece: Skala | KP814011 | Hofman et al. (2016) |
| *Pelophylax bedrigae* | *Pelophylax bedrigae* | N/A | KP260934 | Direct submission by Hofman et al. (2014) |
